# Role of FMRP in rapid antidepressant effects and synapse regulation

**DOI:** 10.1101/2020.08.25.266833

**Authors:** Chelcie F. Heaney, Sanjeev V. Namjoshi, Ayse Uneri, Eva C. Bach, Jeffrey L. Weiner, Kimberly F. Raab-Graham

## Abstract

Rapid antidepressants are novel treatments for major depressive disorder (MDD) and work by blocking N-methyl-aspartate receptors (NMDAR), which, in turn, activate the protein synthesis pathway regulated by mechanistic/mammalian target of rapamycin complex 1 (mTORC1). Our recent work demonstrates that the RNA-binding protein Fragile X Mental Retardation Protein (FMRP) is downregulated in dendrites upon treatment with a rapid antidepressant. Here, we show that the behavioral effects of the rapid antidepressant Ro-25-6981 require FMRP expression, and treatment promotes differential mRNA binding to FMRP in an mTORC1-dependent manner. Further, these mRNAs are identified to regulate transsynaptic signaling. Using a novel technique, we show that synapse formation underlying the behavioral effects of Ro-25-6981 requires GABA_B_R-mediated mTORC1 activity in WT animals. Finally, we demonstrate that in an animal model that lacks FMRP expression and has clinical relevance for Fragile X Syndrome (FXS), GABA_B_R activity is detrimental to the effects of Ro-25-6981. These effects are rescued with the combined therapy of blocking GABA_B_Rs and NMDARs, indicating that rapid antidepressants alone may not be an effective treatment for people with comorbid FXS and MDD.

## Introduction

Over the past 50 years, treatment for major depressive disorder (MDD) has been limited^1^, with selective serotonin reuptake inhibitors (SSRIs) as the most commonly prescribed treatment for the general population. Recently, MDD treatment entered a new era with the discovery that glutamate N-methyl-D-aspartate receptor (NMDAR) antagonists produce antidepressant-like effects^2^, perhaps through blocking NMDARs containing the 2B subunit (GluN2B)^3^. Importantly, these drugs have a rapid onset and work in a treatment-resistant population, with relief of MDD symptoms within hours of a single infusion that can last up to a week^4–6^. Thus, NMDAR antagonists are considered paradigm-shifting treatments for MDD^7^.

Activation of mechanistic/mammalian target of rapamycin complex 1 (mTORC1), a serine-threonine kinase that regulates protein synthesis and promotes new synapse formation, is key to reducing MDD symptoms through NMDAR antagonism^8^. mTORC1 regulates protein synthesis in dendrites by utilizing mRNAs that are targeted and translated in a site-specific and temporal fashion. The coordinated expression of these mRNAs, which code for proteins involved in synaptic and structural plasticity^9,10^, may be regulated by RNA-binding proteins (RBPs). Our recent work suggests that NMDAR antagonism rapidly downregulates dendritic expression of the RBP Fragile X Mental Retardation Protein (FMRP)^11^. While FMRP is extensively characterized in the context of ASD, its role in MDD and MDD treatments is unclear.

FMRP is one of the best characterized RBPs, affecting mRNA trafficking^12^, mRNA repression in dendrites^13^, and is implicated in Fragile X Syndrome (FXS), the leading single-gene cause of autism spectrum disorder (ASD)^14,15^. Loss-of-function mutations in the FMR1 gene, which codes for FMRP^16^, lead to FXS. Notably, people who are FMR1 premutation carriers exhibit higher rates of MDD and depressed mood compared to the general population^17–19^. People with ASD have an increased risk of developing MDD by the time they are 30 years old compared to people without ASD^20^, and caretakers of people with FXS identify mood and depression as a priority for treatment development^21^. Moreover, SSRIs do not produce an antidepressant-like effect in an animal model of FXS^22^, suggesting that people with comorbid FXS and MDD may be treatment-resistant to traditional MDD pharmacotherapies.

Critical questions arise in light of these data: (1) is FMRP required for the rapid antidepressant properties of NMDAR antagonists? (2) Given the importance of mTORC1 signaling in rapid antidepressant efficacy, do mTORC1 and FMRP regulate specific cellular functions? And (3), are rapid antidepressants an effective treatment for MDD in FXS?

Herein, we demonstrate that rapid antidepressant-mediated transsynaptic signaling and presynaptic engagement are mTORC1-dependent and are required for new synapse formation. These mechanisms underlying the rapid antidepressant efficacy of the GluN2B-specific antagonist Ro-25-6981 require activity-dependent FMRP expression. Moreover, we demonstrate that, unlike wild-type (WT) mice, treatment with Ro-25-6981 alone is insufficient to promote new synapse formation or antidepressant-like effects on behavior in *Fmr1* knockout (KO) mice, similar to SSRIs^22^. Finally, we provide data that γ-aminobutyric acid B receptor (GABA_B_R) signaling prevents the actions of Ro-25-6981 in *Fmr1* KO mice. Blocking GABA_B_Rs with the specific antagonist CGP35348 restores the antidepressant-like effect on behavior and increases synapse number when coadministered with Ro-25-6981 in *Fmr1* KO mice. Importantly, our findings propose a new combination therapy effective to treat MDD in FXS.

## Materials and Methods

### Detailed Materials and Methods are outlined in the Supplement

#### Animals

Male and female WT (C57BL/6, Jackson Laboratory) and *Fmr1* KO (B.6129P2-Fmr1tm1Cgr/J, Jackson Laboratory) mice between the ages of 2-5 months were used. All animals were randomly assigned to treatment groups. We found no significant effect of sex or sex X treatment interaction (data presented in Supplement). All experiments were carried out in accordance with the National Institutes of Health’s Guide for the Care and Use of Laboratory Animals and approved by the Wake Forest Institutional Animal Care and Use Committee (IACUC).

#### Pharmacology

For behavioral and *in vivo* experiments, animals received intraperitoneal (i.p.) injections of Ro-25-6981 (10 mg/kg, Tocris 1594) with or without CGP35348 (100 mg/kg; Tocris, 1245), or 0.9% physiological saline vehicle (0.01 ml/g, LabChem). Animals in the RIP-Seq experiment received either Ro-25-6981 (10 mg/kg) with or without rapamycin (6 mg/kg; LC Laboratories, #R-5000), or 0.9% physiological saline (200 μl). For *in vitro* experiments, Ro-25-6981 (10 μM), rapamycin (200 nM), and baclofen (50 μM; Tocris, #0796) were used. Vehicle controls cells were treated with water (vehicle for Ro-25-6981 and baclofen) and/or 0.1% DMSO (vehicle for rapamycin).

#### Forced swim test

Animals were placed into a glass cylinder filled with water 23-25°C for six minutes; animals were quickly removed at the end, dried, and returned to their warmed homecage. Each animal was run in two FST sessions (see Fig. 1A for timeline); the first session acutely stressed the animals, and the second session, 24 hours after the first, was video recorded and scored for immobility in the last four minutes using Ethovision (version 11.5, Noldus).

**Figure 1.**
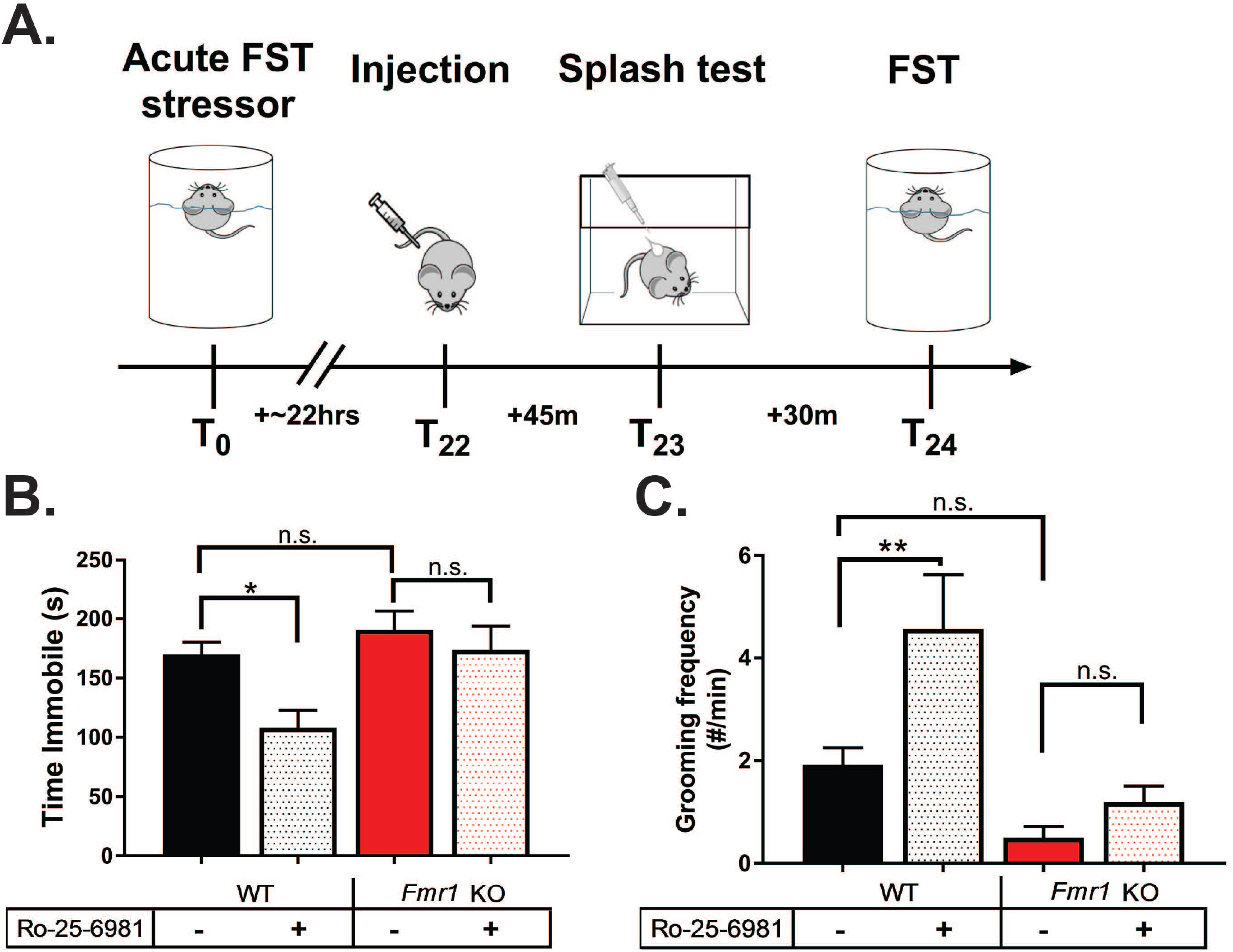
The rapid antidepressant behavioral effects of Ro-25-6981 require FMRP expression. **A.** Representative timeline of behavioral procedures. Twenty-four hours prior to the recorded forced swim test (FST), animals were acutely stressed with FST. Approximately 22 hours later, animals received a single i.p. injection of their randomly assigned treatment. Forty-five minutes after the injection, animals underwent the splash test, and 30 minutes after that, they were exposed to a second FST. **B.** WT animals treated with Ro-25-6981 significantly decreased immobility in the FST, but *Fmr1* KO mice did not respond to treatment. Two-way ANOVA revealed a significant main effect of treatment (F_1,58_=6.152, *p*=0.0161) and genotype (F_1,58_=7.386, *p*=0.0087) but did not find a significant genotype X treatment interaction. Newman-Keuls post-hoc revealed a significant difference between WT control (169.9±10.4 s, n=14) vs. WT Ro-25-6981 (108.1±14.7 s, n=17), but no significant difference between KO control (190.5±16.0 s, n=16) and KO Ro-25-6981 (173.7±20.0 s, n=15). KO control and KO Ro-25-6981 were significantly more immobile than WT Ro-25-6981. **C.** Similarly, Ro-25-6981-treated WT animals increased grooming frequency but treated *Fmr1* KO mice did not. Two-way ANOVA revealed a significant main effect of treatment (F_1,57_=7.29, *p=*0.0091) and genotype (F_1,57_=15.16, *p=*0.0003), but there was not a significant genotype X treatment interaction. Newman-Keuls post-hoc revealed a significant difference between WT control (1.92±0.32 times per minute, n=13;) vs. WT Ro-25-6981 (4.56±1.06, n=16), but no significant difference between KO control (0.5±0.22 times per minute, n=16) and KO Ro-25-6981 (1.18±0.31, n=16). Bars represent mean±SEM. *=*p*<0.05, **=*p*<0.01, n.s.=not significant.

#### Splash test

Animals were individually assessed in their homecage by applying 200 μl of 10% sucrose to their dorsal coat. Animals were video-recorded for five minutes and later scored for grooming behavior (licking, grooming with forepaws, and scratching).

#### RNA Immunoprecipitation (RIP)

Animals were sacrificed 45 minutes after drug administration, and cortices were rapidly dissected out and flash frozen on dry ice (Fig. S1A). RIP-Seq was modified from previous methods shown to significantly reduce background mRNA binding^23,24^.

#### Immunoblotting

Animals were sacrificed 45 minutes after drug administration, and cortices were processed according to previously published methods^11,25–27^.

#### Immunohistochemistry and *in vivo* proximity ligation assay (PLA)

Animals were deeply anesthetized 45 minutes after drug administration and underwent transcardial perfusion. Brains were sectioned 25 μm thick on a Leica SM2010F sliding microtome and put into cryoprotectant. Slices were permeabilized and blocked before overnight primary antibody incubation at 4°C. Slices underwent a two-hour incubation at 37°C (PLA) or RT (immunohistochemistry) in secondary antibodies, then incubated in ligase for 30 minutes at 37°C (PLA). Finally, slices underwent the amplification/polymerase step for two hours at 37°C (PLA).

#### Cell culture and *in vitro* PLA

Primary neuronal hippocampal cultures were prepared from postnatal day 0-3 mouse pups from WT or *Fmr1* KO mice according to previously published methods^11,25–27^. Cells were plated between 70-100,000 per 12 mm on glass coverslips, and cultures were used at 18-21 days *in vitro*. Drug treatments were done in media. Fixed neurons were blocked and permeabilized and then incubated with primary antibodies for one hour at RT, followed by secondary antibody incubation for one hour at RT (immunocytochemistry) or 37°C (PLA). PLA coverslips were then incubated with ligation solution for 30 minutes at 37°C, followed by incubation in amplification/polymerase buffer for 100 minutes at 37°C.

#### Microscopy and analysis

All images within a given experiment were acquired and analyzed using the same settings with a Nikon A1plus confocal microscope. Images were analyzed with Nikon NIS-Elements AR (version 4.40.00).

#### Statistical Analyses

Prism (version 7.05, GraphPad) was used for all statistical analyses. Statistical comparisons were made using Student’s t-test, one-way analysis of variance (ANOVA), or two-way ANOVA where appropriate. Newman-Keuls post-hoc tests were performed in the event of significant ANOVAs. Outliers were determined using Grubbs’ test (α=0.05). All data are expressed as mean ± standard error of the mean (SEM).

## Results

### The rapid antidepressant behavioral effects of Ro-25-6981 require FMRP expression

We recently showed that dendritic FMRP expression is rapidly reduced in WT dendrites after treatment with the GluN2B-specific antagonist Ro-25-6981, suggesting that activity-dependent expression of FMRP and FMRP-dependent translation in dendrites may be necessary for antidepressant efficacy^11^. To test this hypothesis, we treated acutely stressed WT and *Fmr1* KO mice with a single i.p. injection of Ro-25-6981 (see Fig. 1A for timeline). We then analyzed performance in two well-characterized behavioral tasks that measure antidepressant efficacy – the forced swim test (FST), a measure of behavioral despair, and the splash test, a measure of motivated self-care^2,28–31^. Because the splash test uses sucrose, one could argue that this task measures reward-related behavior. However, WT and *Fmr1* KO mice do not display baseline differences in sucrose preference^32^; further, others show that the splash test is sensitive enough to detect behavioral changes due to stress and antidepressant treatment without showing changes to sucrose preference^33^. Consistent with our previous results^25,26^, Ro-25-6981-treated WT animals are mobile about 60s more in the FST (Fig. 1B) and groom about twice as frequently in the splash test (Fig. 1C), indicative of an antidepressant-like effect. However, Ro-25-6981 does not affect performance in either task when administered to *Fmr1* KO mice. Together, these results confirm our hypothesis that the RBP FMRP is required for the behavioral effects of rapid antidepressants.

### mTORC1-sensitive FMRP-target mRNAs are regulated by Ro-25-6981 treatment

Next, we set out to identify the FMRP-target mRNAs regulated by Ro-25-6981, which may underlie its antidepressant efficacy. We treated WT and *Fmr1* KO mice with Ro-25-6981 and collected brains 45 minutes post-administration, a time point where behavioral changes^25^ and mTORC1 activation^2^ are observed. Cortices were processed for RNA immunoprecipitation (RIP), followed by RNASeq (RIP-Seq), using a specific antibody against FMRP^34^ (see Supplemental Methods, Fig. S1-S3, and Tables S1-S3 for RIP-Seq data preprocessing, filtering, and quality control). As a control for nonspecific binding, FMRP-RIP was performed, in parallel, on *Fmr1* KO cortices. Ro-25-6981 treatment promoted the differential binding of 283 transcripts to FMRP relative to control (Fig 2A; light blue circle). These data indicate that FMRP regulates the availability of mRNAs for translation in response to Ro-25-6981 treatment.

**Figure 2.**
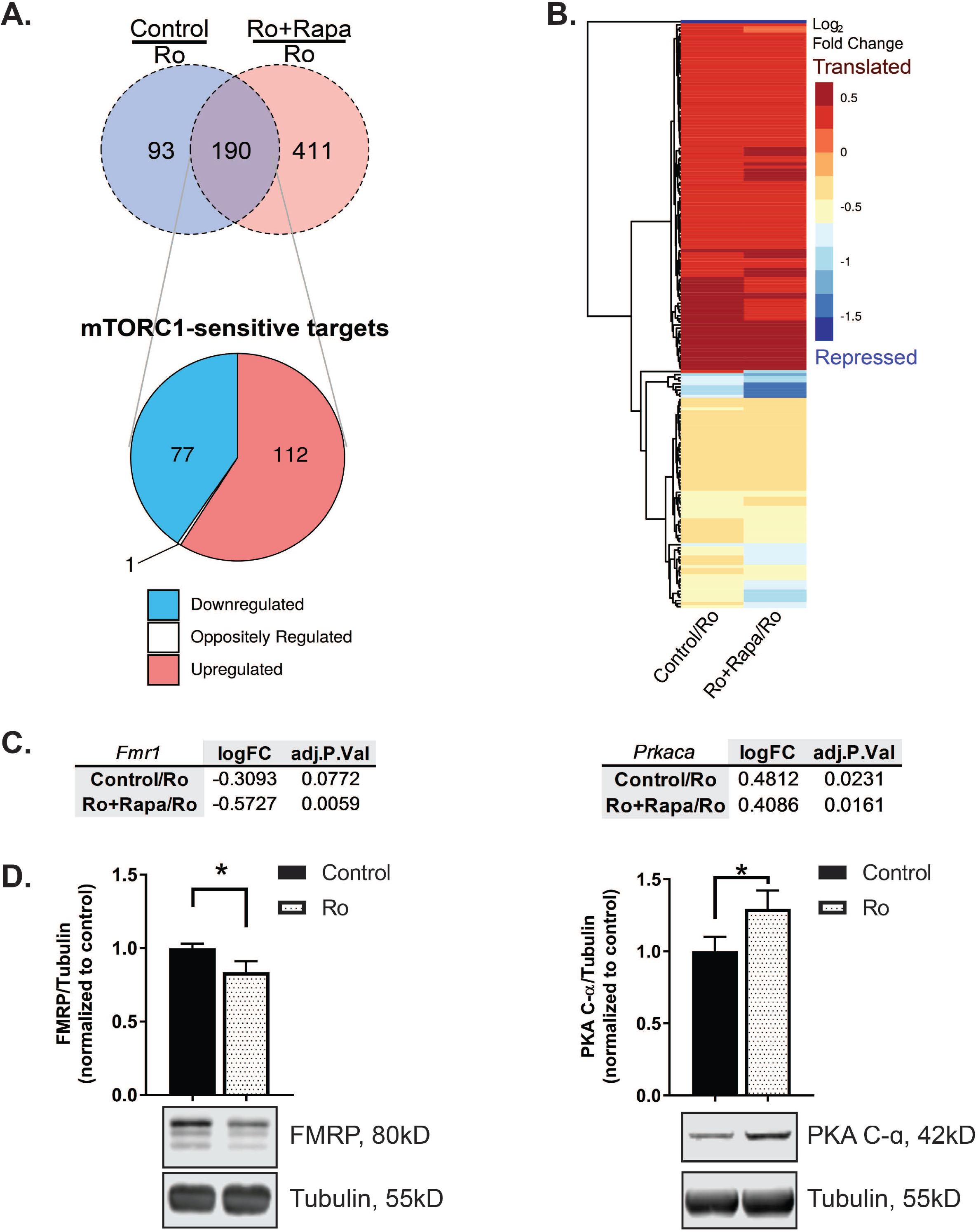
mTORC1-sensitive FMRP-target mRNAs are regulated by Ro-25-6981 treatment. **A.** *Top:* Venn diagram of differentially expressed mRNAs isolated in Control/Ro-25-6981 or Ro-25-6981+Rapamycin/Ro-25-6981 conditions with the FDR cutoff of 0.1 (same mRNAs as in Fig. 2B). *Bottom*: Breakdown of log_2_ fold-change direction of mRNAs in the intersection of the Venn diagram, indicative of the mRNAs that are sensitive to both treatment of Ro-25-6981 and mTORC1 activity. See Tables S4 and S5 for a complete list of genes and fold-changes. The single mRNA that is oppositely regulated is Dbp, a transcription factor implicated in regulating circadian rhythm. **B.** Clustered dendrogram and heatmap for log_2_ fold-change for the treatments. Targets are the same as those in the pie chart in Fig. 2A. Dendrogram created using the average linkage method and Euclidean distance metric. Positive fold-changes indicate mRNAs are released from FMRP and presumably translated with Ro-25-6981 treatment; negative fold-changes indicate that more mRNAs are bound to FMRP with Ro-25-6981 treatment compared to the control (indicative of mRNA repression with Ro-25-6981 treatment) or Ro+Rapa (indicative that mTORC1 activity is needed for mRNAs to be released with Ro-25-6981 treatment) conditions. **C.** In both Control/Ro-25-6981 and Ro-25-6981+Rapamycin/Ro-25-6981 conditions, *Fmr1* has a negative fold-change, indicating *Fmr1* mRNAs remain bound to FMRP with Ro-25-6981 treatment and is mTORC1 sensitive. Conversely, *Prkaca* has a positive fold-change in both Control/Ro-25-6981 and Ro-25-6981+Rapamycin/Ro-256981 conditions, indicating that *Prkaca* mRNAs are released from FMRP with Ro-25-6981 treatment and is mTORC1 sensitive. Significance determined with the FDR cutoff of 0.1. **D.** Verification that protein expression of FMRP decreases^11^ (control=1.0±0.03 normalized optical density, n=8; Ro-25-6981=0.83±0.07, n=8; t_14_=2.008, *p=*0.0322) and PKA C-α increases (control=1.0±0.10 normalized optical density, n=7; Ro-25-6981=1.29±0.12, n=7; t_12_=1.797, p=0.0488) with Ro-25-6981 treatment in cortical lysates 45 minutes after treatment. *=*p*<0.05

We and others show that rapid antidepressants, including Ro-25-6981, increase mTORC1-dependent synthesis of several synaptic proteins necessary for its antidepressant effects^2,25,26^. To identify the mTORC1-sensitive transcripts among the 283 FMRP-target mRNAs we identified, we performed RIP-Seq on cortices from mice co-injected with Ro-25-6981 plus the mTORC1 inhibitor rapamycin. We then compared the differentially regulated FMRP-target mRNAs sensitive to both Ro-25-681 treatment (Fig. 2A, light blue circle) and mTORC1 activity (Fig. 2A, light red circle). Differential gene expression analysis between these two treatments revealed 190 overlapping transcripts (false discovery rate cutoff of 0.1; Fig. 2A, 2B; Table S4, S5^35^). Of these overlapping FMRP-target mRNAs sensitive to both Ro-25-6981 treatment and mTORC1 activity, 99.5% change in the same direction (58.9% upregulated/released, 40.5% downregulated/bound) (Fig. S1B). FMRP is best known for repressing mRNA translation^36^, and we expected that treatment with Ro-25-6981 would induce FMRP to release select transcripts to be translated (~59% are released/upregulated). We were surprised to find that FMRP also bound a new set of target mRNAs (~41% become bound/downregulated). To verify our RIP-seq results, we identified target mTORC1-sensitive mRNAs that were either bound to (*Fmr1*) or released from (*Prkaca*) FMRP after Ro-25-6981 treatment (Fig. 2C, Table S5). We analyzed FMRP and PKA C-α protein expression *in vivo* 45 minutes after treatment. As expected, we found that mice treated with Ro-25-6981 express significantly less FMRP, in line with our previous results^11^, and more PKA C-α in cortical lysates by Western blot analysis (Fig. 2D). Together, these data verify our RIP-Seq results and suggest that Ro-25-6981 regulates the dynamic expression of FMRP and PKA C-α.

### FMRP-target mRNAs regulate transsynaptic signaling

Next, we used the Database for Annotation, Visualization and Integrated Discovery (DAVID 6.8)^37,38^ to cluster the resulting transcripts based on their coordinated biological processes (gene ontology, GO), to identify critical mechanisms mediated by Ro-25-6981/mTORC1/FMRP (Fig. 3A, 3B). These analyses revealed transsynaptic signaling to be the most significantly enriched GO cluster in both conditions (Fig. 3A, control/Ro; Fig. 3B, Ro+Rapa/Ro). We used biomaRt^39–41^ to identify the genes associated with transsynaptic signaling, which were then intersected with our data set to determine the fold-changes for these genes based on condition. Notably, of the 190 Ro-25-6981/mTORC1-sensitive FMRP target mRNAs (Fig. 2A), ~60% are transsynaptic (Fig. 3C, *upper right quadrant*, Fig. S4D, Table S5). These analyses suggest that regulating transsynaptic signaling is a specific role for the FMRP-target mRNAs sensitive to both mTORC1 activity and Ro-25-6981 treatment.

**Figure 3.**
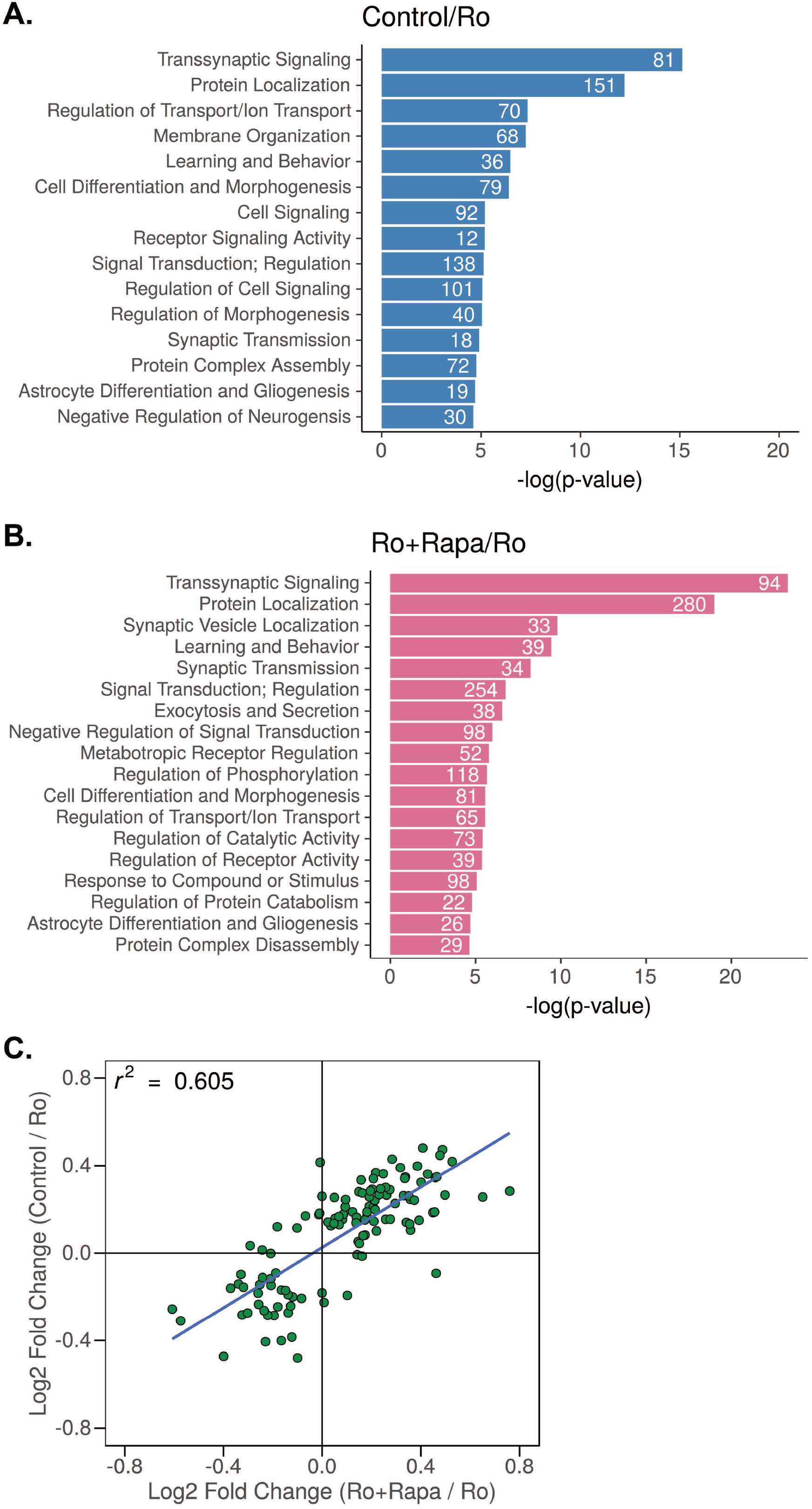
FMRP-target mRNAs regulate transsynaptic signaling. **A.** Uniquely enriched GO biological process clusters for FMRP targets identified by RIP-Seq where log_2_(Control/Ro-25-6981 fold-change) > 0 (*blue)* and **B** log_2_(Ro-25-6981+Rapa/Ro-25-6981 fold-change) > 0 (*pink*). Transsynaptic signaling is the most significantly enriched GO cluster in both conditions. **C.** Scatterplot showing log_2_(Control/Ro-25-6981 fold-change) > 0 log_2_(Ro-25-6981+Rapa/Ro-25-6981 fold-change) > 0 of the mTORC1-sensitive transsynaptic targets (*upper right quadrant*). Supplemental Tables S4 and S5 contain the complete list of identified targets.

### Transsynaptic signaling requires both NMDAR blockade and GABA_B_R-mediated mTORC1 activation

Transsynaptic signaling facilitates the coordinated communication of pre-and postsynaptic terminals^42^. Additionally, new synapse formation is suggested to underlie rapid antidepressant efficacy^2^. We have previously shown that activation of traditionally inhibitory GABA_B_Rs in the presence of an NMDAR antagonist unexpectedly increases L-type calcium channel activity, which thereby increases dendritic mTORC1 activation and downstream protein synthesis^25,26^. To validate that Ro-25-6981 affects transsynaptic signaling, we capitalized on the ability to tightly control mTORC1 activity in the cell culture system. We measured the expression of two prototypical synaptic proteins that specifically localize to post- and presynaptic terminals: postsynaptic density 95 (PSD-95) and synapsin-1 (SYN1), respectively^2,34^. Notably, both NMDAR antagonism and GABA_B_R activity are required to activate mTORC1^25^. Thus, we treated cultured primary hippocampal neurons (see Fig. 4A for timeline) with Ro-25-6981 (no mTORC1 activity *in vitro*), Ro-25-6981 with the GABAB agonist baclofen (mTORC1 activity *in vitro*), or Ro-25-6981 with baclofen and the mTORC1 inhibitor rapamycin (mTORC1 blocked *in vitro*). We found that GluN2B-specific antagonism increased the expression of the postsynaptic marker PSD-95 in an mTORC1-independent manner (Fig. 4B, 4C). In contrast, the expression of the presynaptic marker SYN1 increased only in the neurons with mTORC1 activity (Ro-25-6981+Baclofen; Fig. 4B, 4D; S5A). When we measured colocalization of PSD-95 with SYN1, however, we did not find a significant effect of treatment (Fig. 4E), likely due to these proteins being juxtaposed versus overlapping. Realizing the limitations of colocalization as a correlate of synapse number, we developed a fluorescent assay that could detect pre- and postsynaptic engagement and, thus, indicate new synapse formation. Array tomography studies demonstrate that postsynaptic PSD-95 and presynaptic SYN1 localize within 40 nm of each other^43^. We surmised we could utilize a proximity ligation assay (PLA) to detect protein-protein interactions between proximal pre- and postsynaptic proteins located across the synaptic cleft within 0-40 nm. Consistent with our immunostaining data, NMDAR antagonism alone (Ro-25-6981) did not increase synapse formation, as indicated by no change in PLA puncta number relative to control (Fig. 4F, 4G). However, when we activated mTORC1 (Ro-25-6981+baclofen), the number of detectable PLA puncta in dendrites increased by ~ 65% (Fig. 4F, 4G). Further, when we inhibited mTORC1 with rapamycin (Ro-25-6981+Rapa+Bac), the number of detected synapses decreased back to baseline (Fig. 4F, 4G). Thus, GABA_B_R-mediated mTORC1 activation is required for presynaptic engagement for synapse formation with Ro-25-6981.

**Figure 4.**
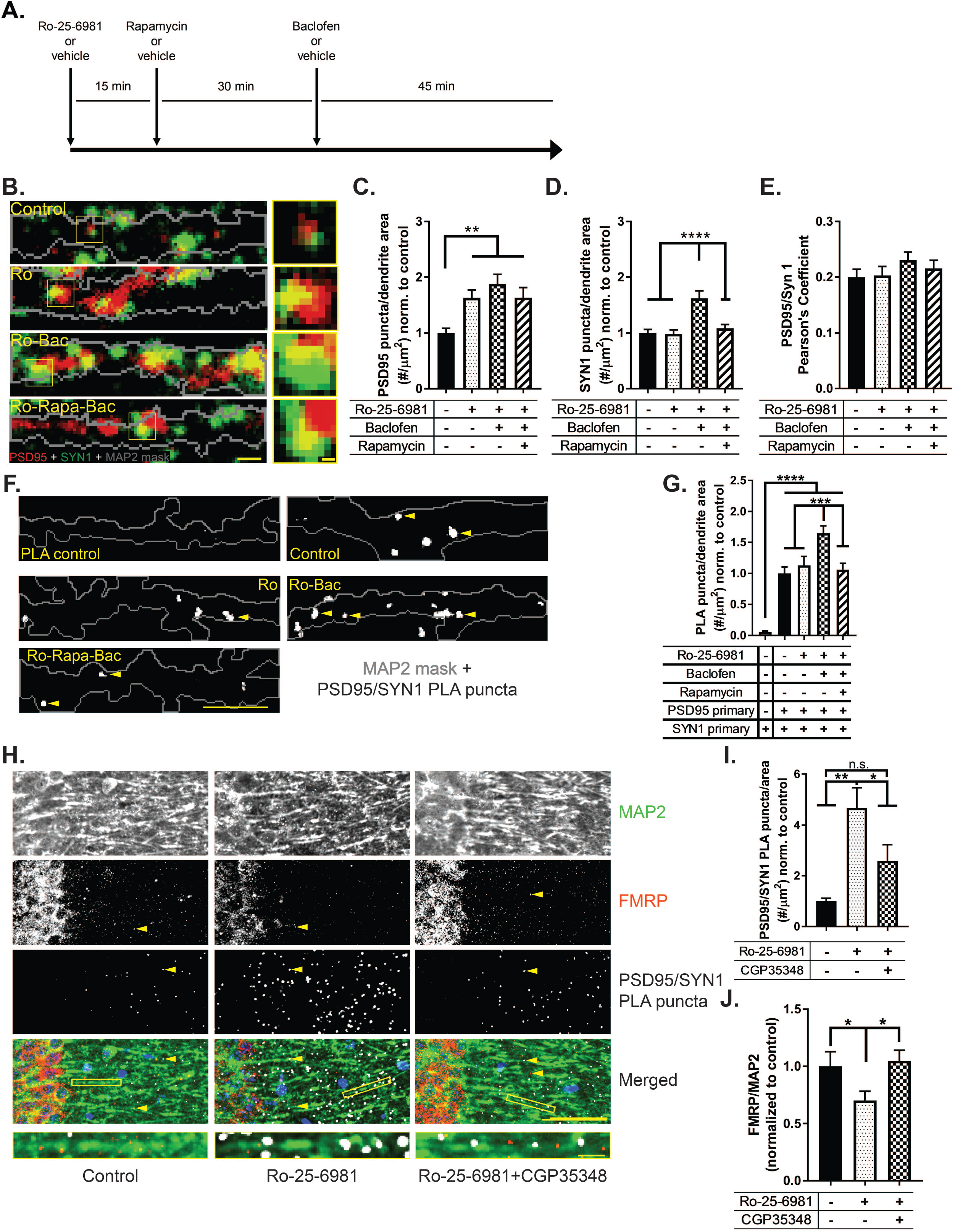
Transsynaptic signaling requires both NMDAR blockade and GABA_B_R-mediated mTORC1 activation. **A.** Timeline of treatments for neuronal culture experiments in 4B and 4F. **B.** Representative immunofluorescence images of treated WT hippocampal neurons stained for PSD-95 (red), SYN1 (green), and MAP2 (grey outline). Panels depict 10 μm segments of dendrites, at least 20 μm from the soma; scale bar=1 μm. Highlighted boxes indicate representative SYN1 and PSD-95 staining; scale bar=0.2 μm. **C.** Quantification of average number of PSD-95 puncta within dendritic MAP2 area by treatment. PSD-95 expression increased in all treatments in an mTORC1-independent manner. One-way ANOVA revealed a significant main effect of treatment (F_3,184_=6.303, *p=*0.0004). Newman-Keuls post-hoc revealed a significant difference between control (1.0±0.08 normalized optical density, n=49) vs. Ro-25-6981 (1.63±0.14, n=46), control vs. Ro-25-6981+Baclofen (1.88±0.17, n=41), and control vs. Ro-25-6981+Rapamycin+Baclofen (1.63±0.17, n=52). **D.** Quantification of average number of SYN1 puncta within dendritic MAP2 area by treatment. One-way ANOVA revealed a significant main effect of treatment (F_3,184_=11.14, *p*<0.0001). Newman-Keuls post-hoc revealed Ro-25-6981+Baclofen (1.61±0.13 normalized optical density, n=41) significantly increased the number of SYN1 puncta compared to control-treated neurons (1.0±0.06, n=49), as well as neurons treated with Ro-25-6981 (0.98±0.07, n=46) and Ro-25-6981+Rapamycin+Baclofen (1.08±0.06, n=52), indicating an mTORC1-dependent effect. **E.** Quantification of Pearson’s correlation coefficient of PSD-95 and SYN1 by treatment. One-way ANOVA did not reveal a significant effect of treatment for the correlation between PSD-95 and SYN1 (F_3,186_=0.8282, *p*=0.4799). **F.** Representative dendritic traces of treated hippocampal neurons. White puncta represent proximity-detected PSD-95 and SYN1 proteins by PLA (indicated by yellow arrowheads), and grey outline represents MAP2 staining of the neuron. Puncta at least 12.5 μm from the soma were analyzed. Scale bar=10 μm. **G.** Quantification of the average number of PSD-95/SYN1 PLA puncta per dendritic MAP2 area. One-way ANOVA revealed a significant main effect of treatment (F_4,581_=18.25, *p*<0.0001). Newman-Keuls post-hoc revealed Ro-25-6981+Baclofen (1.65±0.11 puncta/μm^2^ normalized to control, n=132) significantly increased the number of PSD-95/SYN1 PLA puncta compared to control-treated (1.0±0.10 puncta/μm^2^ normalized, n=119), Ro-25-6981-treated (1.12±0.14 puncta/μm^2^ normalized, n=125), and Ro-2506981+Rapamycin+Baclofen-treated (1.06±0.10 puncta/μm^2^ normalized, n=143) neurons. Additionally, PLA control neurons (0.05±0.01 puncta/μm^2^ normalized, n=67) showed a significant lack of puncta compared to all other treatments. **H.** Representative images of treated WT CA1 stratum radiatum dendrites. White puncta represent proximity-detected PSD-95 and SYN1 proteins by PLA, and red puncta represent FMRP (yellow arrowheads indicate representative puncta of each). **I.** Quantification of the average number of PSD-95/SYN1 PLA puncta per area of CA1 stratum radiatum dendrites. Puncta at least 10 μm from the pyramidal layer were analyzed. One-way ANOVA revealed a significant main effect of treatment (F_2,52_=6.79, *p*=0.0024). Newman-Keuls post-hoc revealed that treatment with Ro-25-6981 (4.68±0.78 puncta/μm^2^ normalized to control, n=23) significantly increased PSD-95/SYN1 PLA puncta 45 minutes after treatment compared to control (1.0±0.11 puncta/μm^2^ normalized, n=13) and Ro-25-6981+CGP35348 (2.58±0.64 puncta/μm^2^ normalized, n=19). This increase is blocked with the GABA_B_R antagonist CGP35348 (no significant difference between control and Ro-25-6981+CGP35348). **J.** Protein expression of FMRP decreases with Ro-25-6981 treatment in CA1 stratum radiatum dendrites 45 minutes after treatment (control=1.0±0.12 normalized optical density, n=19; Ro-25-6981=0.70±0.07, n=23; t40=2.026, *p=*0.0247). Scale bar=50 μm. Representative dendrites shown below to demonstrate the puncta are located on dendrites. Scale bar=5 μm. Bars represent mean±SEM. *=*p*<0.05, **=*p*<0.01, ***=*p*<0.001, ****=*p*<0.0001, n.s.=not significant.

We next verified that the change in synapse number detected *in vitro* is reflected *in vivo*. Because there are few inhibitory neurons in hippocampal cultures^44,45^, baclofen is necessary to activate GABA_B_Rs *in vitro*. However, *in vivo* treatment of ketamine or Ro-25-6981 is sufficient to increase endogenous GABA^46^ and promote mTORC1 activity and antidepressant-like effects on behavior^25,26^. Therefore, we inhibited GABA_B_Rs to investigate the effects of GABA_B_R-mediated mTORC1 activity *in vivo*. Thus, we treated WT mice with Ro-25-6981 or Ro-25-6981 and the specific GABA_B_R antagonist CGP35348 to block GABA_B_R signaling and downstream mTORC1 activation^25^. Indeed, consistent with our *in vitro* PLA results, we found that Ro-25-6981 treatment increased the number of detected synapses and that inhibiting mTORC1 activity with CGP35348 prevented this increase (Fig. 4H, 4I, S5B, S5C). Further, dendritic FMRP expression decreased in these same slices with Ro-25-6981 treatment, which returned to baseline expression when GABA_B_R signaling was blocked with Ro-25-6981+CGP35358 (Fig 4H, 4J).

### GABA_B_R activity is detrimental to synapse formation and rapid antidepressant efficacy of Ro-25-6981 in a preclinical mouse model of FXS

Next, we sought to validate the requirement of FMRP expression for mTORC1-dependent synapse formation upon treatment with Ro-25-6981. We replicated the PLA-synapse assay in hippocampal slices from *Fmr1* KO mice. We predicted that if the activity-dependent release of FMRP-target mRNAs is required for mTORC1-dependent transsynaptic signaling and synapse formation, then the relative increase in synapses detected by PLA upon treatment will be absent in the *Fmr1* KO mouse. Surprisingly, we instead found a significant decrease of about 43% in the number of detected synapses in *Fmr1* KO mice treated with Ro-25-6981 (Fig. 5A, 5B). Further, treatment with Ro-25-6981 and CGP35348 unexpectedly reversed this decrease and significantly increased the number of detected synapses by about 47% compared to control animals (Fig. 5A, 5B). These data suggest that in the presence of FMRP expression, GABA_B_R signaling is required for rapid antidepressant-mediated synaptogenesis; however when FMRP expression is absent, GABA_B_R signaling is detrimental and needs to be blocked for synaptogenesis to occur.

**Figure 5.**
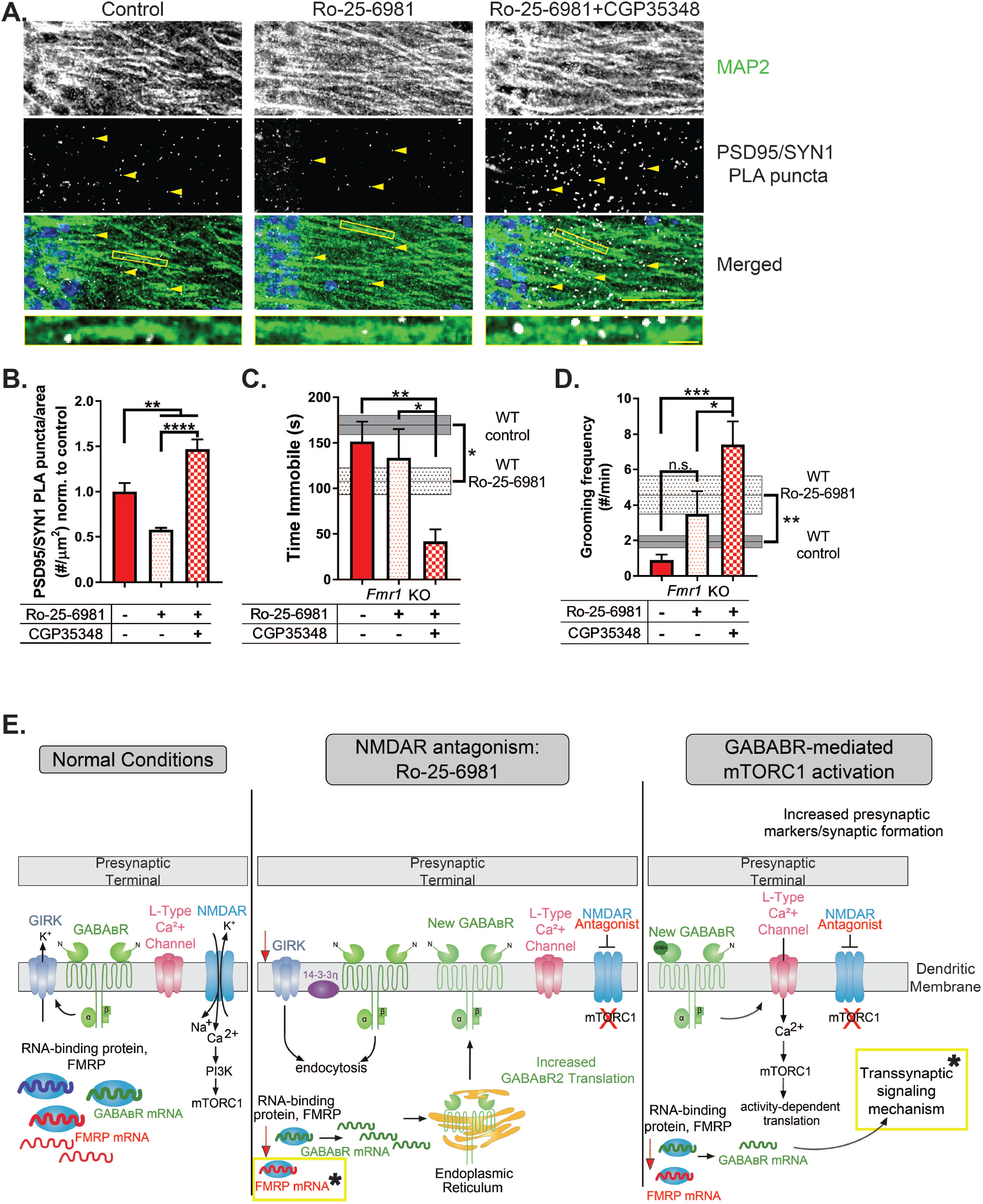
GABA_B_R activity is detrimental to synapse formation and rapid antidepressant efficacy of Ro-25-6981 in a preclinical mouse model of FXS. **A.** Representative images of treated KO CA1 stratum radiatum dendrites. White puncta represent proximity-detected PSD-95 and SYN1 proteins by PLA (indicated by yellow arrowheads). Scale bar=50 μm. Representative dendrites shown below to demonstrate the puncta are located on dendrites. Scale bar=5 μm. **B.** Quantification of the average number of PSD-95/SYN1 PLA puncta per area of CA1 stratum radiatum dendrites. Puncta at least 10 μm from the pyramidal layer were analyzed. One-way ANOVA revealed a significant main effect of treatment (F_2,10_=25.45, *p=*0.0001). Newman-Keuls post-hoc revealed Ro-25-6981 (0.57±0.02 puncta/μm^2^ normalized to control, n=4) significantly decreases PLA puncta compared to control (1.0±0.09 puncta/μm^2^ normalized, n=4), but blocking GABA_B_Rs with CGP35348 increases the number of puncta (1.47±0.10 puncta/μm^2^ normalized, n=5) compared to both other treatments. **C.** One-way ANOVA revealed a significant main effect of treatment (F_2,24_=6.157, *p=*0.0069) in the FST. Newman-Keuls post-hoc revealed *Fmr1* KO animals treated with Ro-25-6981+CGP35348 (41.6±13.7 s, n=9) significantly decrease immobility in the FST compared to control (151.5±21.9 s, n=9), whereas treatment with Ro-25-6981 alone does not (133.5±31.9 s, n=9). WT data from Figure 1 depicted on the y-axis for reference. **D.** Similarly, one-way ANOVA revealed a significant main effect of treatment (F_2,24_=9.616, *p=*0.0009) in the splash test. Newman-Keuls post-hoc revealed Ro-25-6981+CGP35348 (7.4±1.3 times per minute, n=10) significantly increases grooming frequency in the splash test compared to control (0.88±0.30 times per minute, n=9) but not Ro-25-6981-treatment only (3.5±1.2 times per minute, n=8) for *Fmr1* KO animals. WT data from Figure 1 depicted on the y-axis for reference. **E.** Updated rapid antidepressant signaling mechanism model^11,25,26^ to include RIP-Seq results. Under baseline conditions, activated NMDARs increase intracellular calcium, which activates mTORC1 (*left panel*). With NMDAR antagonists like ketamine and Ro-25-6981, NMDARs are blocked but activate the release or binding of mRNAs from the RNA-binding protein FMRP (*middle panel*). Among these released transcripts is GABA_B_R mRNA, which is translated into newly synthesize GABA_B_Rs^25^. Old GABA_B_Rs and the GIRK channels they are coupled to undergo endocytosis^26^. Newly synthesized GABA_B_Rs are coupled to L-type calcium channels and, upon activation, increase intracellular calcium, activating mTORC1 (*right panel*)^26^. mTORC1-dependent transcripts are released from or bound to FMRP, including those mediating a transsynaptic signal that recruits presynaptic engagement to help form new synapses. Yellow boxes with stars indicate the new steps added to the model based on the current data presented. Bars represent mean±SEM. *=*p*<0.05, **=*p*<0.01, ***=*p*<0.001, ****=*p*<0.0001.

If the increase in synapses in WT animals underlies the efficacy of Ro-25-6981 as a rapid antidepressant, we thus questioned whether blocking GABA_B_R activity in conjunction with Ro-25-6981 treatment would promote an antidepressant-like effect on behavior in *Fmr1* KO animals. We tested this hypothesis by examining the effect of Ro-25-6981 with or without the GABA_B_R antagonist CGP35348 on behavior of *Fmr1* KO mice in the FST and splash test (see Fig. 1A for timeline). Indeed, *Fmr1* KO animals treated with Ro-25-6981+CGP35348 decreased immobility in the FST by about 100s (Fig. 5C) and increased grooming frequency in the splash test by nearly eight times compared to control KO animals (Fig. 5D). Importantly, CGP35348 alone does not increase PSD/SYN1 puncta (Fig. S5D, S5E), nor does it affect performance in the FST^11^ or splash test (Fig. S5F) in *Fmr1* KO mice. Thus, our data indicate that the rapid antidepressant efficacy of Ro-25-6981 requires FMRP expression, and treatment with Ro-25-6981 regulates mTORC1/FMRP-target mRNAs that promote transsynaptic signaling and synaptogenesis. However, in cases where FMRP expression is absent, as in *Fmr1* KO animals, Ro-25-6981 is ineffective as a rapid antidepressant. Only when GABA_B_R activity is blocked in these animals does Ro-25-6981 increase synaptogenesis and promote antidepressant-like effects.

## Discussion

Previously, we discovered that the molecular mechanism underlying the behavioral efficacy of rapid antidepressants requires GABA_B_R-mediated mTORC1 activity (Fig. 5E)^11,25,26^. Here, we demonstrate that rapid antidepressants regulate mTORC1-sensitive FMRP-target mRNAs that are involved in transsynaptic signaling. While both FMRP and mTORC1 are independently implicated in transsynaptic signaling in *Drosophila* and cultured neurons ^47–49^, our results are the first to demonstrate their coordinated regulation of the transsynaptic signaling required for effective pharmacological therapy in a mammalian model. To this end, we show, with our newly developed assay that detects synapse formation, that rapid antidepressant treatment in conjunction with GABA_B_R activity promotes synaptogenesis *in vitro* and *in vivo* in WT animals, but not in a preclinical model of FXS (*Fmr1* KO mouse). These data collectively confirm the necessity of FMRP expression for the efficacy of rapid antidepressants.

Surprisingly, in *Fmr1* KO animals, GABA_B_R activity prevents the new synapse formation and behavioral effects observed with rapid antidepressant treatment. However, blocking GABA_B_Rs rescues these detrimental effects in *Frm1* KO mice and promotes the rapid antidepressant effects similar to what we see in WT animals. Paradoxically, clinical trials suggest that treatment with the GABA_B_R agonist baclofen improves irritability and social responsiveness in people with FXS and ASD; however, these results are mixed^50–52^. Moreover, baclofen can cause adverse side effects, including hyperactivity and anxiety in FXS patients ^53–55^. Our results raise the question of why GABA_B_R signaling is required for synaptogenesis and antidepressant efficacy in WT mice yet prohibitive in *Fmr1* KO mice. One clue comes from our previous data demonstrating that traditional inhibitory postsynaptic GABA_B_R-GIRK signaling is absent in *Fmr1* KO dendrites and may instead be constitutively excitatory^11^. By blocking GABA_B_R activity, we may be preventing excessive excitation^56^, and the synapse reduction seen in Ro-25-6981-only treated *Fmr1* KO mice (Fig. 5A and 5B).

Precision-based medicine holds promise for treatment-resistant MDD patients; however, understanding the underlying cause of treatment resistance is necessary for tailored pharmacotherapy. Herein, our data indicate that treating MDD in people with FXS or an FMR1 premutation may require combination therapy, as traditional^22^ and rapid antidepressants alone are not effective in animal models of FXS. FXS is not the only neurological disorder with overactive mTORC1. Tuberous Sclerosis Complex and Alzheimer’s Disease also exhibit comorbid MDD and are reported to have overactive mTORC1 signaling^57–59^. Moreover, a recent clinical report indicates that oral rapamycin treatment before ketamine prolongs the antidepressant effects of ketamine in a treatment-resistant population^60^. Together, these data support the idea that rapid antidepressants, combined with other drugs that modify mTORC1 activity, such as CGP35348 or rapamycin, may be more effective in populations with certain comorbid conditions. While NMDAR antagonists certainly have revolutionized the treatment of MDD in a general population, caution should be exercised when utilizing them to treat individuals with MDD and mTORC1-related diseases, and combination therapy should be considered.

## Acknowledgments

We thank Emma K. Erickson for generating preliminary data associated with this project, Farr Niere for his valuable comments and suggestions, and Olivia Cain for her help running analyses.

## Author contributions

KRG, CFH, and SVN designed the studies; CFH, SVN, AU, and ECB, performed the experiments; KRG, CFH, and SVN performed analyses; KRG, JLW, CFH, and SVN wrote and edited the manuscript.

## Data and materials availability

Sequencing data available in the NCBI Sequencing Read Archive (SRA) under submission SUB5029763. Code for RNAseq analysis available at https://github.com/snamjoshi.

### Added note of proof

Since the original submission of this manuscript a recent report utilizes the same PLA assay to detect changes in synapse number.^61^

Supplementary information is available at MP’s website.

