## Supplemental Materials and Methods for "Role of FMRP in rapid antidepressant effects and synapse regulation"

RNA Immunoprecipitation (RIP)

Animals were sacrificed 45 minutes after drug administration, and cortices were rapidly dissected out and flash frozen on dry ice (Fig. S1A). RIP-seq was modified from previous methods shown to significantly reduce background mRNA binding [1,2]. Tissue was homogenized and lysed in polysome lysis buffer (10 mM HEPES pH 7.0, 100 mM KCl, 25 mM EDTA, 5 mM MgCl2, 1 mM DTT, 0.5% NP-40) in a 1:1 tissue-buffer ratio. RNaseOUT (Thermo Fisher Scientific) and protease/phosphatase inhibitors (HALT Protease and Phosphatase Cocktail, Pierce Biotechnology) were added fresh to samples. Samples were rotated for 10 minutes at 4°C to induce swelling and then flash-frozen on dry ice. Samples were thawed at room temperature (RT) to lyse, and nuclei were pelleted at 3000 x g for 10 minutes. Lysates obtained above were pre-cleared by adding 50 µl of washed magnetic bead slurry (Protein A Dynabeads, Thermo Fisher Scientific), rotated for 30 minutes at 4°C. The magnetic beads slurry was washed and resuspended in 8 volumes of NT-2 buffer (50 mM Tris-HCl pH 7.4, 150 mM NaCl, 1 mM MgCl2, 1 mM DTT, 0.05% NP-40 with RNaseOUT/protease & phosphatase inhibitors added fresh) + 5% BSA. 10 µg of either FMRP (Abcam, ab17722) or IgG (Santa Cruz Biotechnologies, sc-2027) antibodies were added to the beads and rotated for 10 minutes at RT. Antibody-bound beads were washed four times with ice-cold NT-2 buffer. For the immunoprecipitation, 4.5 mg of protein from pre-cleared lysates was added to the antibody-bound beads. The antibody-bead-lysate mixture was then diluted at a ratio of 1:5 with NET-2 buffer (20 mM EDTA pH 8.0, and 1 mM DTT in NT-2 buffer; RNaseOUT and protease/phosphatase inhibitors added fresh) and rotated for one hour at RT. Beads were washed six times in ice-cold NT-2 buffer and immediately resuspended in 350 µl TRI Reagent® Solution (Ambion) for 10 minutes at RT. Beads were pelleted, and the supernatant was removed and resuspended in 350 µl of absolute ethanol. RNA was extracted by applying ethanol-resuspended samples to spin columns from the Direct-zol RNA MiniPrep Kit (Zymogen) according to the manufacturer's instructions. Eluted RNA (25 µl) was DNase treated using the TURBO DNA-free kit (Thermo Fisher Scientific).

*cDNA Synthesis and Quantitative Real-Time PCR (qRT-PCR)*

15% of the eluate from DNase-treated RNA samples was reverse-transcribed to cDNA using the iScript cDNA Synthesis Kit (Bio-Rad) in a 20 µl volume according to manufacturer's instructions. qRT-PCR was performed in 20 µl reaction volume using the iQ™ SYBR® Green Supermix (Bio-Rad) and primers for GABABR1, GABABR2, CaMKIIα, and Cacna2δ2 (GeneCopoeia). qRT-PCR was run with the following protocol: 95°C for 10:00, 40 cycles of 95°C for 0:15 followed by 60°C for 1:00, 95°C for 1:00, and 55°C for 1:00. Relative fold-enrichment was determined by the equation $\Delta\Delta Ct = 2^{-(Ct\ FMRP\ RNA\text{-}IP - Ct\ IgG\ RNA\text{-}IP) - (Ct\ FMRP\ input - Ct\ IgG\ input)}$.

*Library preparation*

85% of the eluate from DNase I treated samples were analyzed for quality using the BioAnalyzer (Agilent Genomics). The mean RIN score across all samples was 8.488 (IQR = 8.375 to 8.825) and the mean RNA mass recovered (ng) was 75.31 (IQR = 36.75 to 81.50) (Table S1). Recovered RNA was generally lower in the knockout samples but still had a high RIN. Two biological replicates for the Ro-25-6981 knockout samples had a RIN too low for library preparation and sequencing (RIN = 1.0 and 1.1 respectively). Library preparation was performed at the Genome Sequencing and Analysis Facility (GSAF, The University of Texas at Austin). After sample amplification to > 1 ug by PCR, rRNA was depleted using the Ribo-Zero rRNA Removal Kit (Human/Mouse/Rat) (Epicentre) according to the manufacturer's instructions. RNA was then purified using the RNeasy MinElute Cleanup Kit (Qiagen). Library preparation of the RNA was performed with the NEBNext Ultra Directional RNA Library Prep Kit for Illumina (NEB). Single-end sequencing of RNA (SE 1x50) was performed with the HiSeq4000 (Illumina).

*Quality control assessment of RNAseq libraries*

We sequenced four technical replicates for each RNAseq library resulting in WT libraries ranging from 16-27 million reads (mean = 22.1 million reads, s.d. = 4.3 million reads). Reads obtained in KO libraries were highly variable ranging from 8-35 million reads (mean = 19.6 million reads, s.d. = 10.3 million reads). However, reads from KO libraries mapped lower than those of their WT counterparts (mean KO library mapping rate = 57.64% ± 21.11% (s.d.); mean WT library mapping rate = 68.94% ± 18.59% (s.d)) (Table S2).

All preprocessing was performed in the R environment [3] and terminal commands were sent directly to the shell from R. Code for analysis can be found at https://github.com/snamjoshi. RNAseq libraries from all technical replicates were analyzed for quality using FastQC 0.11.5 [4] with standard parameter settings. We inspected the libraries for a high Q-score (at least 37) across all base-calls. All libraries showed an overall high Q-score as shown in the “per sequence quality scores” distribution of FastQC. However, multiple libraries showed a small drop in Q-score in either the first six base-calls or the last 2-3 base-calls. The interquartile range for these drops never exceeded a minimum Q-score of 32 demonstrating that overall, the sequencing was of excellent quality. In general, any repetitive sequences were below 1% of all sequences in a given library and usually even lower. A BLAST search from the NCBI website [5] revealed that they were primarily ribosomal RNA or adaptor sequences. The technical replicates from the Ro-25-6981 knockout library (RO_KO_2) showed the lowest library complexity as well as abnormal GC content and per base sequence content compared to other libraries. Duplicated sequences were shown to be as high as 2%. We analyzed the contents of this library for ribosomal contamination using RNA- SeQC 1.1.8 [6] but found them to be minimal (< 2% of library sequences). We, therefore, attribute the unusual sequencing results to poor starting material since these libraries showed the lowest RIN scores and RNA concentration. However, they still contain important sequence information, so we opted to keep these libraries for our analysis.

*Trimming of RNAseq libraries*

Since a few libraries showed a lower Q-score toward the beginning and end of the sequences, we applied a trimming tool to the libraries to remove these lower quality base-calls. Trimming would also remove any adaptor contamination since we saw this in a few of the samples. We used the tool Trim Galore 0.4.1 to trim the sequencers and adaptors. Trim Galore utilizes Python 2.7.11 and is wrapped around cutadapt 1.10 [7]. Trim Galore was run as follows: trim_galore –fastqc -o trimmed_fasqc *.gz. Since Trim Galore is pre-loaded with Illumina adaptors, we did not need to specify them as parameters. We ran FastQC again to verify that trimming and adaptor removal was carried out successfully and did not alter the quality of the libraries.

*Alignment of RNAseq libraries and calculating counts from BAM files*

Alignment was performed using Tophat2/Bowtie2 [8,9] using samtools 0.1.19 [10] with the following settings: tophat2 library-type fr-firststrand -G [GTF] p 4 -o [library] [index] [files]. The library-type parameter was chosen to be consistent with the library preparation method. Gene model annotations for M. musculus (mm9) were downloaded directly from Ensembl (assembly version GRCm38.84, Ensembl 84; Mar 2016) [11]. Pre- built reference indices for Bowtie2 were downloaded from the Illumina website. The alignment data shows that most libraries had a relatively strong alignment percentage (greater than 75%). The only exception is the Ro-25-6981 Knockout replicate 2 (16%). This result is not unexpected given the lower library complexity and slightly higher amount of ribosomal RNA contamination. A summary of all library information and alignment data can be found in Table S2.

Rsubread 1.20.6 [12] was used to calculate counts. Strandedness was maintained in counting using the “strandSpecific” parameter. Additionally, we used the “countMultiMappingReads = TRUE” parameter option so we could count ambiguously mapped reads. All BAM files for all four technical replicates were combined so that the final output contains counts for all of our libraries.

*Post-alignment biological replicate quality control*

Count data for each technical replicate was combined using the collapseReplicates() function from DESeq2 [13]. A regularized log transformation was applied to the data before visualization. Data were inspected to see the correlation in count number between different biological replicates (Fig. S2A), principal components analysis to see the similarity in variance among the replicates (Fig. S2B), and a Euclidean distance matrix to assess hierarchical clustering of different libraries based on the overall Euclidean distance between each gene (Fig. S2C).

To verify the validity of our sequencing results, we first took a list of FMRP mRNA targets and non-targets from a previous publication from the Darnell lab and compared the counts for these targets obtained from the WT or Fmr1-KO mice across treatments (Fig. S2D) [14]. We found the normalized counts for the targets were increased by 333.90% (Saline WT), 369.29% (Ro WT), and 365.35% (Ro + rapamycin WT) relative to non-targets in each treatment. This result supports the validity of our obtained libraries. Next, we performed qRT-PCR on our immunoprecipitated RNA using primers for GABABR1 and GABABR2, two FMRP targets we have previously characterized, under Ro or Saline treatments (Fig. S2E). This data confirms the presence of GABABR1/GABABR2 mRNA in our samples and shows that their levels decrease with Ro treatment.

*Filtration/normalization of RNA sequence libraries and differential gene expression analysis*

**Low count filtration, factor size normalization, and overlap comparison to other FMPR dataset**

Combined count data (all technical replicates collapsed) was filtered using HTSFilter 1.14.1 [15] with the standard settings. HTSFilter removes all low counts from the full list of raw counts. Annotation was performed using Bioconductor [16] with the org.Mm.eg.db [17] and AnnotationDbi [18] packages. DESeq2 factor size normalization was then applied to the data. Since this will scale across the entire library, the counts in the knockout libraries will be inflated. In order to correct for this, the ratio between extracted RNA concentrations from WT and *Fmr1* KO tissue (BioAnalyzer results) were used to scale back the KO tissue counts after DESeq2 normalization. WT normalized counts were unchanged. All biological replicates were then averaged together across treatments.

To filter the list further, we first sorted the list by the highest counts in the saline treatment and then compared to other known FMRP target data sets (see supplemental results). This includes two RIP-ChIP datasets [19,20], HITS-CLIP data [14], PAR-CLIP data [20], and APRA-NeuroArray [21]. Our initial analysis showed poor overlap with the Ascano et al. and Miyashiro et al. data sets; poor overlap with the Miyashiro dataset was also seen in Suhl et al. [22]. The Ascano data also contains a large number of RNA targets (> 5000) and may not be as useful for determining a proper cutoff. These sets were excluded from our analysis; however, consensus datasets showing overlap between the Ascano RIP-ChIP (A-RIP), Ascano PAR-CLIP (A-PAR), Brown

(B), and Darnell (D) datasets from Suhl et al. were included in our analysis. Cutoff was determined by sorting the list by highest normalized saline counts and then ranking the percent of matches between the (D), (B) data, and consensus data (B_D_A-RIP, B_D_A-PAR, B_D, and D_A-PAR). By including 4120 genes from our data, we have a 95% overlap with all genes obtained in the above data (Fig. S3A). Lastly, we filtered the data by removing mitochondrial and glial cells. These genes were derived from the Brain RNAseq database [23] and MitoMiner [24].

*Filtering background counts by WT-to-KO fold enrichment*

Background was assessed by two methods. 1) We computed the WT-to-KO fold enrichment for all major treatment libraries. Next, we took a list of FMRP non-targets [14] and plotted the WT-to-KO fold enrichment of these non-targets from our data against the WT-to-KO fold enrichment of the targets identified in the other datasets the overlapped in our data (Fig. S3B). The aim here was to determine a cutoff between the average WT-to-KO fold enrichment for targets versus non-targets, the latter of which were presumed to have a higher background and thus a lower value for the WT-to-KO fold ratio. We then took the average of the lower quartile of the WT-to-KO fold enrichment across all the different data sets (Fig. S3B, dotted red line) and used this as an estimation of the cutoff for background. 2) We used univariate K-means with the Ckmeans.1d.dp package [25] to quantify the distribution of signal and background counts into two clusters (Fig. S3C, S3D). Gaussian mixture modeling (GMM), a generalization of K-means, has been used previously to characterize signal-to-noise for RIP-SEQ/RIP- ChIP data [26,27], however, our data did not converge using GMM. We then averaged the cutoffs from the two methods used above for a WT-to-KO fold enrichment ratio cutoff of ~1.112788. We applied this value across all treatments to filter any data that did not meet this cutoff criterion.

*Motif analysis of FMRP target*

To further assess the validity of our data, we used an FMRP motif analysis similar to Suhl et al. [22]. For this analysis, we did not use the differentially expressed genes selected by Limma/voom because motif frequency/number was not significantly higher than that of the non-targets. All sequences were obtained from BioMart [28] using the biomaRt [29,30] package from Bioconductor. Sequences were obtained from the coding region because the majority of FMRP binding motifs appear to cluster in these parts of the gene [31]. Since some genes may return multiple sequences, we only used sequences of max length for our calculations. Motifs were counted using the count() function from the seqinr [32] package. The QFM sequence was counted using the matchPattern() function from the biostrings [33] package and a regular expression. Rho is computed using the rho() function from seqinr. To compute multiple rho, which is necessary for motifs for which a position can take on more than one nucleotide value (e.g., ACUK), a function called multipleRho() is used which computes rho manually according to the following formula:

$$\frac{[frequency(motif1) + frequency(motif2)]}{[expected(motif1) + expected(motif2)]}$$

Motif statistics were calculated using base R functions except for the effect size which uses the cohensD() function from the lsr package [34]. For all motif statistics (freq/kB, t- test, Wilcoxon-Mann-Whitney test, effect size) except rho, we followed the general presentation of data from Suhl et al. [22] (Figure S3E, Table S3).

*Differential gene expression analysis*

Differential gene expression (DGE) analysis was performed using Limma/voom [35,36]. Raw counts were imported into R and then normalized via TMM/CPM. The contrast matrix was set up to compare fold changes for either Saline/Ro-25-6981 (SAL_WT – RO_WT) or Ro-25-6981+Rapa/Ro-25-6981 (RO_RAP_WT – RO_WT). These values were then visualized using an MA plot (Fig. S4A, *top*, and S4B, *top*) and a volcano plot (Fig. S4A, *bottom*, and S4B, *bottom*). In these plots, the colored value represent mRNA outside the False Discovery Rate (FDR) cutoff of 0.1

We determined mTORC1-sensitive targets using a combination of a Venn diagram and pie chart (Fig. 2A). First, the output from Limma/voom was first filtered for a $\log_2$ fold-change between -2 and 2. To create the Venn diagram, the Control/Ro relative complement was obtained by filtering this data set with an FDR cutoff of 0.1. The Ro+Rapa/Ro relative complement was obtained by filtering this data set with an FDR cutoff of 0.1. The intersection was obtained by filtering both data sets with this filter. The differentially expressed genes in this intersection were then filtered for those with a fold-change greater than zero for both treatments (“Upregulated”) or less than zero for both treatments (“Downregulated”). One remaining gene was oppositely regulated between both conditions. A clustered dendrogram for these targets was produced using the pheatmap package (average cluster method,

Euclidean distance metric, k = 6 clusters) (Fig. 2B).

*Gene Ontology (GO) clustering*

The Database for Annotation, Visualization and Integrated Discovery (DAVID 6.8) [37,38] was queried for gene ontology annotation using the RDAVIDWebService package [39] (Fig. S4C). To cluster GO terms with high similarity, we used an approach similar to that of the Enrichment Map plugin [40,41] for Cytoscape 3.0 [42]. First, we obtained DAVID output from their functional annotation chart service from the "GOTERM_BP_ALL" ontology. The up-regulated and down-regulated genes were analyzed separately by DAVID. Then, we filtered these GO terms using a p-value cutoff of 0.001 and an FDR cutoff of 0.05. We then took the relative complement of either the up-regulated or down- regulated GO terms. We then use a clustering approach to group GO terms of high similarity (based on the number of overlapping genes) from each relative complement together in a dendrogram. To determine the degree of overlap between genes in any two ontologies we used the average of the Jaccard cutoff (JC) and the overlap coefficient (OC).

$$JC = \frac{|A \cap B|}{|A \cup B|}$$

$$OC = \frac{|A \cap B|}{Min(|A|, |B|)}$$

A higher overlap score indicates a greater number of genes in common. We next created a matrix of the average JC and OC for every GO-term against all other GO-terms. The matrix was filtered for a similarity cutoff of > 0.5. This matrix was then clustered to create a dendrogram using the factoextra package [43]. To determine optimal cluster number, we minimized the within-sum-of-squares (WSS). Each branch of the dendrogram indicates a degree of overlap between the different GO terms. Each clustered branch was then manually annotated for a general biological category that described the GO term. We then averaged the total number of genes in each of these clusters (Fig. 3A, 3B).

*Transsynaptic genes fold-change correlations*

First, we obtained a list of trans-synaptic genes using biomaRt using the “go_id” attribute and the “go_parent_term” filter for the following GO ids: postsynaptic density (GO:0014069), presynapse (GO:0098793), postsynapse (GO:0098794), and trans-synaptic signaling (GO:0099537). The genes associated with trans-synaptic signaling were then intersected with our data set to determine the fold-changes for these genes based on condition. We then plotted the log2 fold changes for the transsynaptic genes in both conditions. In Figure 3C only the transsynaptic genes are shown. In Figure S4D all of our filtered FMRP targets are shown. Targets in the interquartile range were colored grey to indicate “background” or no change. All data outside of the range of -2 to 2 log2 fold-change were removed before plotting. Only data with an adjusted p-value less than 0.1 were included.

*Other R packages for analysis*

In addition to the packages already mentioned, we utilized a number of other packages for our analysis. For plotting: ggplot2 [44], ggthemes [45], RColorBrewer [46], and gridExtra [47]. For annotation: mgu74a.db [48] and org.Hs.eg.db [49]. For functions used in

DGE analysis: edgeR [50]. For data import: xlsx [51] and R.utils [52]. For data transformation and processing: reshape2 [53], magrittr [54], dplyr [55], and gdata [56].

Immunoblotting

Animals were sacrificed 45 minutes after drug administration, and cortices were rapidly dissected out and flash frozen on dry ice. Cortices were homogenized and lysed in RIPA buffer (150 mM NaCl, 10 mM Tris pH 7.4, 0.1% SDS, 1% Triton X-100, 1% deoxycholate, 5 mM EDTA, and 1x HALT Protease and Phosphatase Cocktail) with a motorized pestle. Samples were rotated at 4ºC for one hour, then centrifuged 14,000 x g for 20 minutes at 4ºC. The resulting supernatants were analyzed with Pierce BCA Protein assay kit (Thermo Fisher Scientific) to determine protein concentration. An equal amount of protein was loaded into a 10% SDS-polyacrylamide gel, separated by molecular weight, and transferred to a 0.22 μm nitrocellulose membrane. Membranes were blocked in 5% bovine serum albumin (Sigma Aldrich) in tris-buffered saline (TBS) for one hour. Primary antibodies (rabbit anti-FMRP (1:1000, Abcam ab17722), rabbit-anti-PKA C-α (1:1000, Cell Signaling #4782) rabbit or mouse anti-tubulin (1:10,000)) were applied overnight at 4ºC in TBS + 0.1% Tween20 (TBST). Membranes were then washed three times for 10 minutes each in TBST before being incubated in secondary antibodies in TBST for one hour (goat anti-rabbit IRDye 680 (1:4,000, LICOR) or goat anti-mouse IRDye 800 (1:4000, LICOR)) at RT. Membranes were again washed three times for 10 minutes in TBST, followed by two TBS washes for five minutes before being imaged using a LICOR Odyssey imaging system. Densitometry analysis was performed in LICOR Image Studio Lite.

Immunohistochemistry and *in vivo* proximity ligation assay

Animals were deeply anesthetized 45 minutes after drug administration and underwent transcardial perfusion of cold phosphate-buffered saline (PBS), followed by cold 4% paraformaldehyde (PFA). Whole brains were then dissected out and stored in 4% PFA overnight at 4ºC, followed by overnight at 4ºC in 15% sucrose, and overnight at 4ºC in 30% sucrose. Brains were sectioned 25 μm thick on a Leica SM2010F sliding microtome and put into cryoprotectant (15% ethylene glycol, 10% glycerol in 0.05 M PBS). Slices were washed three times for 10 minutes each in PBS and 0.75% glycine before being permeabilized and blocked (10% normal donkey serum (NDS), 0.25% Tween20 in PBS) for two hours at RT. Slices used for PLA were incubated overnight in primary antibodies (mouse anti-PSD95 (1:400, Neuromab 75-028), goat anti-synapsin1 (1:200, Santa Cruz s.c.7379), chicken anti-MAP2 (1:500, Abcam ab5392)) at 4ºC in blocking buffer; slices used for immunohistochemistry were incubated overnight (rabbit anti-FMRP (1:400, Abcam ab17722), chicken anti-MAP2 (1:500, Abcam ab5392)). Slices were washed three times for 10 minutes in PBS/glycine, followed by a two-hour incubation at 37ºC in secondary antibodies (PLA: donkey anti-mouse PLUS (1:5, Sigma-Aldrich DUO92001), donkey anti-goat MINUS (1:5, Sigma-Aldrich DUO92006), immunohistochemistry: donkey anti-rabbit 555 (1:400, AlexaFluor), donkey anti-chicken 488, (1:400, Invitrogen SA1-72000)). PLA slices went through two, ten-minute washes in Duolink Buffer A, then incubated in ligase for 30 minutes at 37ºC and again washed twice for 10 minutes in Duolink Buffer A. Slices then underwent the amplification/polymerase step for two hours at 37ºC, followed by two 20-minute washes in Duolink Buffer B and a single one minute wash in 1% Duolink Buffer B. Slices were mounted to slides with Duolink mounting media (Sigma-Aldrich DUO82040). Immunohistochemistry slices were washed two times for 10 minutes in PBS/glycine, followed by a 20 minute DAPI wash; then, slices were mounted to slides with Aqua- Poly/Mount (Polysciences, 18606-20).

Cell culture and *in vitro* immunocytochemistry and proximity ligation assay

Primary neuronal hippocampal cultures were prepared from postnatal day 0-3 mouse pups from WT or *Fmr1* KO mice according to previously published methods [57-60]. Cells were plated between 70-100,000 per 12 mm on glass coverslips coated with 50 µg/mL poly-D-lysine and 100 µg/mL laminin. After four hours *in vitro*, the cell culture medium was replaced with Neurobasal Medium (Thermo Fisher Scientific) supplemented with 2% B27 supplement and 0.5 mM GlutaMAX™ and maintained in 5% $CO_2$ at 37°C. After four days *in vitro*, cytosine arabinofuranoside (2 µM) was added to the cultures. Neuronal cultures were used at 18-21 days *in vitro*. Drug treatments were done in media. After treatment, cells were fixed in 4% PFA for 15 minutes at RT and stored in fresh PBS at 4°C until used for immunocytochemistry or proximity ligation assay. Neurons were blocked and permeabilized with 10% NDS and 0.25% Tween20 in PBS for 30 minutes, then were incubated with primary antibodies (1:500 mouse anti-PSD-95, Neuromab; 1:500 goat anti-synapsin-1, Santa Cruz; 1:1000 chicken anti-MAP2, Aves MAP) for one hour at RT. After being washed with PBS three times for ten minutes, cells were incubated in secondary antibodies (immunofluorescence: 1:400 Alexa Fluor 405 goat anti-mouse, Life Technologies

A31553; 1:400 Alexa Fluor 647 goat anti-rabbit, Life Technologies, A21245; 1:400 Alex Fluor 488 goat anti-chicken, A11039; PLA: 1:5 donkey anti-mouse PLUS, Sigma-Aldrich; 1:5 donkey anti-goat MINUS, Sigma-Aldrich; 1:400 donkey anti-chicken 488, Invitrogen), in blocking buffer for one hour at RT (immunofluorescence) or 37°C (PLA). Immunofluorescence coverslips were washed in PBS (two times for ten minutes each, plus one ten minute DAPI wash) before being slide-mounted. PLA coverslips were then incubated with ligation solution for 30 minutes at 37°C. Cells were washed in Duolink Buffer A twice for five minutes each, followed by incubation in amplification buffer for 100 minutes at 37°C. Cells were washed in Duolink Buffer B twice for ten minutes, and one final one minute 1% Duolink Buffer B wash. Coverslips were slide-mounted using Duolink mounting media.

Microscopy and analysis

All images within a given experiment were acquired and analyzed using the same settings on a Nikon A1plus confocal microscope. Hippocampal slices were imaged using an air 20x lens at 1024x104 pixels in a given Z plane where the hippocampus was in focus based on MAP2 staining. All slides were labeled without treatment information. Experimenter was blind to specific treatments until statistical analysis. To create a region of interest (ROI), the Nikon NIS-Elements AR (version 4.40.00) “Draw Bezier ROI” function was used to trace over the CA1 stratum radiatum. PLA puncta were identified using the “spot detection” function and converted to a binary mask. Puncta within the drawn ROI were counted with the “binary in ROI” function. Average intensities within the drawn ROI for MAP2 and FMRP staining were obtained using the “ROI data” function. Neurons were imaged using an oil-immersion 60X lens with sequential scanning. For immunofluorescence images, dendrites were imaged at 1024x1024 pixels in a given Z plane. Puncta masks for immunofluorescence images were created based on intensity after thresholding images. ROIs were chosen based on MAP2 signal, extending at least 20 μm from the cell body, and included primary, secondary, or tertiary branches. Puncta within each ROI were counted with the "binary in ROI" function and normalized to the area of the dendritic ROI. For PLA, a max projection of a 3 x 1 μm Z-stack of 1024 x 1024 pixels was obtained. PLA puncta were chosen based on a standard intensity above background applied to all images and converted to a binary mask. Puncta within dendritic ROIs identified by MAP2 signal at least 12.5 μm from the soma were counted with the "binary in ROI" function and normalized to the area of the dendritic ROI.

**Supplementary Text**

Reproducibility

Animals were pseudo-randomly assigned treatments, such that, in a given cage, each treatment was represented at least once. At the time of randomization, animals were assigned a unique identifier and tail color marking. These identifiers were used to identify animals during scoring of behavior, and to which experimenter was blind.

Figure 1 and 5: Behavioral experiments shown in Figure 1 and Figure 5 were repeated three times, with an n=8-9 per group for WT and n=3-9 for KO mice. Individual mice were removed from forced swim analyses if they exhibited poor swimming ability.

Figure 2 and Figure 3: The RIP-Seq experiment was run once, with an n=3 for WT and KO for each treatment group (total experimental n=18). Tissue for Western blot analyses was obtained from one experimental iteration (n=8/treatment); data were removed from analyses if a statistical outlier. Immunofluorescence analysis (n=8-9 animals per treatment, 2-3 slices per animal analyzed; slices were removed from analysis if MAP2 staining was poor) was from two experimental iterations. Figure 4: Immunofluorescence experiment was from two independent culture with two separate coverslips treated per treatment. 41-52 dendrites per treatment were analyzed, with no more than three dendrites analyzed from a single neuron. PLA analyses were conducted on two independent cultures with two separate coverslips per treatment. 67- 143 dendrites were analyzed from 8-10 neurons per coverslip. Slice PLA analyses were from two independent experiments, n=3-9 animals per treatment, 2-3 slices per animal analyzed; slices were removed from analysis if MAP2 staining was poor. Figure 5: Slice PLA analyses were from one experimental iteration, n=3/treatment with 3 slices per animal analyzed; slices were removed from analysis if MAP2 staining was poor. Sample sizes were based on previous experiments and were limited to availability within our colony.

### Supplementary Results

#### *Filtration of RNA-IP data results in a high-quality dataset enriched for multiple neuronal GO clusters and differentially expressed FMRP targets*

Attempts to characterize FMRP binding motifs have identified multiple common patterns among targets including the G-quadraplex motif, U-rich sequences, and other common motif patterns [14,20,22,31,61-67]. Although there is no consensus on which patterns are common to all FMRP targets, Suhl et al. have attempted to statistically characterize the occurrence of all these motifs within known FMRP datasets. Our approach was to partially replicate their overall analysis strategy with our targets to see if we obtained consistent results as a way to validate our FMRP target set compared to previously identified FMRP targets.

We characterized the enrichment of putative FMRP target motifs across our filtered dataset compared to genes that had been filtered out of our dataset in previous processing steps described above. We also calculated the occurrence of the motif within sequences that were filtered out of the list ("non-targets"). We consider a motif to be enriched in our FMRP targets (n = 2696) if the frequency of occurrence per kilobase (kB) was significantly higher than the occurrence in non-targets (n = 11222). Sequences were used specifically from the coding region because the majority of FMRP motifs are found here [31]. Additionally, we also calculated ρ which tells us if the occurrence of a DNA string is over- or under-represented according to what we would expect by chance. For example, ρ = 10 would indicate the occurrence of the sequence is 10 times more likely that we would expect by chance.

Data for motif analysis is summarized in Figure S3E and Table S3. The mean frequency/kB of the ACUK motif in the FMRP targets was not significantly enriched compared to the non-targets (targets = 8.036, non-targets = 8.483). High enrichment was seen for the GAC motif (targets = 16.787, non-targets = 16.066), GACR (targets = 7.905, non-targets = 7.444), and GACARG motifs (targets = 0.875, non-targets = 0.773). While the WGGA motif occurred at a high frequency/kB (freq./kB = 16.531) there was significant difference in motif frequency compared to non-targets (freq./kB = 16.666). Finally, the QFM motif showed a significant enriched motif frequency per gene compared to non-targets (targets = 1.170, non-targets = 1.074). Importantly, the effect size (Cohen's d) is small in all cases indicating that although there is a significant difference in motif frequency/kB between targets (n = 2696) and non-targets (n = 11222) for all motifs except WGGA, the relationship may have moderately low practical significance. However, similar effect values were reported by Suhl et al. [22]. Finally, ρ was relatively small for all motifs (mean $\rho_{targets}$ = 0.371, mean $\rho_{non\text{-}targets}$ = 0.370) but in all cases where there was significant enrichment, the motif occurred with a greater frequency then we would expect by chance. The GAC motif showed the strongest motif frequency compared to chance ($\rho_{targets}$ = 0.955, $\rho_{non\text{-}targets}$ = 0.936). Overall, the results of our analysis were similar to those seen in the analysis of the FMRP consensus data sets by Suhl et al., [22] though there was some variability in the exact motif frequency in our data compared to the consensus data sets. Taken together, these results support the validity of our methodology and confirm that the motif frequency in the mRNA we isolated have similar enrichment over non-targets to previously identified FMRP targets.

*No behavioral differences between mice by sex*

We determined there were no significant differences between sex or sex by treatment interactions for the mice for the behavioral tasks by two-way ANOVA. See table below for statistics.

| | Main effect of sex | | Interaction | |
|---|---|---|---|---|
| | FST | Splash test | FST | Splash test |
| WT (Fig. 1) | $F_{1,27}$=0.81, p=0.3761 | $F_{1,25}$=2.898, p=0.1011 | $F_{1,27}$=1.524, p=0.2277 | $F_{1,25}$=0.2707, p=0.6074 |
| KO (Fig. 1) | $F_{1,27}$=8.249, p=0.0078 NK post-hoc does not show any significant comparisons | $F_{1,28}$=3.351, p=0.0778 | $F_{1,27}$=0.1772, p=0.6771 | $F_{1,28}$=1.123, p=0.2983 |
| KO (Fig. 5) | $F_{2,21}$=6.794, p=0.0165 NK post-hoc does not show any significant comparisons | $F_{2,21}$=2.721, p=0.1139 | $F_{2,21}$=1.35, p=0.2808 | $F_{2,21}$=0.49, p=0.6195 |

14 Darnell JC, Van Driesche SJ, Zhang C, Hung KYS, Mele A, Fraser CE *et al.*

FMRP stalls ribosomal translocation on mRNAs linked to synaptic function and autism. *Cell* 2011; **146**: 247–261.

15 Rau A, Gallopin M, Celeux G, Jaffrézic F. Data-based filtering for replicated high- throughput transcriptome sequencing experiments. *Bioinformatics* 2013; **29**: 2146– 2152.

16 Huber W, Carey VJ, Gentleman R, Anders S, Carlson M, Carvalho BS *et al.* Orchestrating high-throughput genomic analysis with Bioconductor. *Nat Methods* 2015; **12**: 115–121.

17 Carlson M. org.Mm.eg.db: Genome wide annotation for Mouse. 2016.

18 Pages H, Carlson M, Falcon S, Li N. AnnotationDbi: Annottion Database Interface. 2017.

19 Brown V, Jin P, Ceman S, Darnell JC, O'Donnell WT, Tenenbaum SA *et al.* Microarray identification of FMRP-associated brain mRNAs and altered mRNA translational profiles in fragile X syndrome. *Cell* 2001; **107**: 477–487.

20 Ascano M, Mukherjee N, Bandaru P, Miller JB, Nusbaum JD, Corcoran DL *et al.* FMRP targets distinct mRNA sequence elements to regulate protein expression. *Nature* 2012; **492**: 382–386.

21 Miyashiro KY, Beckel-Mitchener A, Purk TP, Becker KG, Barret T, Liu L *et al.* RNA cargoes associating with FMRP reveal deficits in cellular functioning in Fmr1 null mice. *Neuron* 2003; **37**: 417–431.

22 Suhl JA, Chopra P, Anderson BR, Bassell GJ, Warren ST. Analysis of FMRP mRNA target datasets reveals highly associated mRNAs mediated by G-quadruplex structures formed via clustered WGGA sequences. *Hum Mol Genet* 2014; **23**: 5479–5491.

23 Zhang Y, Chen K, Sloan SA, Bennett ML, Scholze AR, O'Keeffe S *et al.* An RNA- sequencing transcriptome and splicing database of glia, neurons, and vascular cells of the cerebral cortex. *J Neurosci* 2014; **34**: 11929–11947.

24 Smith AC, Blackshaw JA, Robinson AJ. MitoMiner: a data warehouse for mitochondrial proteomics data. *Nucleic Acids Res* 2012; **40**: D1160–D1167.

25 Wang H, Song M. Ckmeans.1d.dp: Optimal k-means Clustering in One Dimension by Dynamic Programming. *R J* 2011; **3**: 29–33.

26 Erhard F, Dölken L, Zimmer R. RIP-chip enrichment analysis. *Bioinformatics* 2013; **29**: 77–83.

27 Lu Z, Guan X, Schmidt CA, Matera AG. RIP-seq analysis of eukaryotic Sm proteins identifies three major categories of Sm-containing ribonucleoproteins.

*Genome Biol* 2014; **15**: R7.

28 Smedley D, Haider S, Durinck S, Pandini L, Provero P, Allen J *et al.* The BioMart community portal: an innovative alternative to large, centralized data repositories. *Nucleic Acids Res* 2015; **43**: W589–98.

29 Durinck S, Moreau Y, Kasprzyk A, Davis S, De Moor B, Brazma A *et al.* BioMart and Bioconductor: a powerful link between biological databases and microarray data analysis. *Bioinformatics* 2005; **21**: 3439–3440.

30 Durinck S, Spellman PT, Birney E, Huber W. Mapping identifiers for the integration of genomic datasets with the R/Bioconductor package biomaRt. *Nat Protoc* 2009; **4**: 1184–1191.

31 Anderson BR, Chopra P, Suhl JA, Warren ST, Bassell GJ. Identification of consensus binding sites clarifies FMRP binding determinants. *Nucleic Acids Res* 2016; **44**: 6649–6659.

32 Charif D, Lobry JR. SeqinR 1.0-2: A Contributed Package to the R Project for Statistical Computing Devoted to Biological Sequences Retrieval and Analysis. In: *Biological and Medical Physics, Biomedical Engineering*. Springer Berlin Heidelberg: Berlin, Heidelberg, 2007, pp 207–232.

33 Pages H, Aboyoun P, Gentleman R, Debroy S. Biostrings: String objects representing biological sequences, and matching algorithms. 2016.

34 Navarro D. *Learning statistics with R*. Lulu. com, 2015 http://books.google.com/books?id=6ScxuwEACAAJ&dq=learning+statistics+with+r+a+tutorial+for+pscyhology+students+and+other+beginners&hl=&cd=2&source=gbs_api.

35 Law CW, Chen Y, Shi W, Smyth GK. voom: Precision weights unlock linear model analysis tools for RNA-seq read counts. *Genome Biol* 2014; **15**: R29.

36 Ritchie ME, Phipson B, Wu D, Hu Y, Law CW, Shi W *et al.* limma powers differential expression analyses for RNA-sequencing and microarray studies. *Nucleic Acids Res* 2015; **43**: e47–e47.

37 Huang DW, Sherman BT, Lempicki RA. Bioinformatics enrichment tools: paths toward the comprehensive functional analysis of large gene lists. *Nucleic Acids Res* 2009; **37**: 1–13.

38 Huang DW, Sherman BT, Lempicki RA. Systematic and integrative analysis of large gene lists using DAVID bioinformatics resources. *Nat Protoc* 2009; **4**: 44–57.

39 Fresno C, Fernández EA. RDAVIDWebService: a versatile R interface to DAVID. *Bioinformatics* 2013; **29**: 2810–2811.

40 Merico D, Isserlin R, Stueker O, Emili A, Bader GD. Enrichment map: a network- based method for gene-set enrichment visualization and interpretation. *PLoS ONE* 2010; **5**: e13984.

41 Merico D, Isserlin R, Bader GD. Visualizing gene-set enrichment results using the Cytoscape plug-in enrichment map. *Methods Mol Biol* 2011; **781**: 257–277.

42 Shannon P, Markiel A, Ozier O, Baliga NS, Wang JT, Ramage D *et al.* Cytoscape: a software environment for integrated models of biomolecular interaction networks. *Genome Res* 2003; **13**: 2498–2504.

43 Kassambara A, Mundt F. factoextra: Extract and Visualize the Results of Multivariate Data Analyses. 2016.

44 Wickham H. ggplot2: Elegant Graphics for Data Analysis. 2009; : 213.

45 Arnold JB. ggthemes: extra themes, scales and geoms for ‘ggplot2’. *R package version* 2013; **2**: e79.

46 Neuwirth E. RColorBrewer: ColorBrewer Palettes. 2014.

47 Auguie B. gridExtra: Miscellaneous functions for ‘grid’ graphics. R package version 2.2. 1. 2016.scholar.google.com.

48 Carlson M. mgu74a.db: Affymetrix Murine Genome U74v2 annotation data (chip mgu74a). 2016.

49 Carlson M. org.Hs.eg.db: Genome wide annotation for Human. 2016.

50 Robinson MD, McCarthy DJ, Smyth GK. edgeR: a Bioconductor package for differential expression analysis of digital gene expression data. *Bioinformatics* 2010; **26**: 139–140.

51 Dragulescu AA. xlsx: Read, write, format Excel 2007 and Excel 97/2000/XP/2003 files. 2014.

52 Bengtsson H. R. utils: Various programming utilities. *R package version* 2010; **1.**scholar.google.com.

53 Wickham H. Reshaping Data with the reshapePackage. *J Stat Softw* 2007; **21**. doi:10.18637/jss.v021.i12.

54 Bache SM, Wickham H. magrittr: a forward-pipe operator for R. R package version. 2014; **1**.scholar.google.com.

55 Wickham H, Francois R. dplyr: A Grammar of Data Manipulation. 2016.

56 Warnes GR, Bolker B, Gorjanc G, Grothendieck G, Korosec A, Lumley T *et al.*

gdata: Various R Programming Tools for Data Manipulation. 2015.

57 Workman ER, Niere F, Raab-Graham KF. mTORC1-dependent protein synthesis underlying rapid antidepressant effect requires GABABR signaling. *Neuropharmacology* 2013; **73**: 192–203.

58 Workman ER, Haddick PCG, Bush K, Dilly GA, Niere F, Zemelman BV *et al.* Rapid antidepressants stimulate the decoupling of GABA(B) receptors from GIRK/Kir3 channels through increased protein stability of 14-3-3η. *Mol Psychiatry* 2015; **20**: 298–310.

59 Wolfe SA, Workman ER, Heaney CF, Niere F, Namjoshi S, Cacheaux LP *et al.* FMRP regulates an ethanol-dependent shift in GABABR function and expression with rapid antidepressant properties. *Nat Commun* 2016; **7**: 12867.

60 Niere F, Namjoshi S, Song E, Dilly GA, Schoenhard G, Zemelman BV *et al.* Analysis of Proteins That Rapidly Change Upon Mechanistic/Mammalian Target of Rapamycin Complex 1 (mTORC1) Repression Identifies Parkinson Protein 7 (PARK7) as a Novel Protein Aberrantly Expressed in Tuberous Sclerosis Complex (TSC). *Mol Cell Proteomics* 2016; **15**: 426–444.

61 Schaeffer C, Bardoni B, Mandel JL, Ehresmann B, Ehresmann C, Moine H. The fragile X mental retardation protein binds specifically to its mRNA via a purine quartet motif. *EMBO J* 2001; **20**: 4803–4813.

62 Dolzhanskaya N, Sung YJ, Conti J, Currie JR, Denman RB. The fragile X mental retardation protein interacts with U-rich RNAs in a yeast three-hybrid system. *Biochem Biophys Res Commun* 2003; **305**: 434–441.

63 Menon L, Mihailescu M-R. Interactions of the G quartet forming semaphorin 3F RNA with the RGG box domain of the fragile X protein family. *Nucleic Acids Res* 2007; **35**: 5379–5392.

64 Menon L, Mader SA, Mihailescu M-R. Fragile X mental retardation protein interactions with the microtubule associated protein 1B RNA. *RNA* 2008; **14**: 1644– 1655.

65 Phan AT, Kuryavyi V, Darnell JC, Serganov A, Majumdar A, Ilin S *et al.* Structure-function studies of FMRP RGG peptide recognition of an RNA duplex-quadruplex junction. *Nat Struct Mol Biol* 2011; **18**: 796–804.

66 Ray D, Kazan H, Cook KB, Weirauch MT, Najafabadi HS, Li X *et al.* A compendium of RNA-binding motifs for decoding gene regulation. *Nature* 2013; **499**: 172–177.

67 Chen E, Joseph S. Fragile X mental retardation protein: A paradigm for translational control by RNA-binding proteins. *Biochimie* 2015; **114**: 147–154.

### Supplemental Figure Legends

**Fig. S1. A.** Workflow for RIP-Seq experiment and analysis. **B**. A detailed description of the experimental analysis. Our three experimental conditions under which FMRP was isolated during RIP-Seq (Control, Ro-25-6981, or Ro-25-6981+Rapa) are hypothesized to isolate three different populations. Left: In the control condition, the global condition, we will pull down FMRP transcripts not Ro-25-6981-released (A) as well as Ro-25-6981-released FMRP transcripts (B). Thus, the control condition is represented as the union between these two sets. Middle: Upon Ro-25-6981 injection, a number of the transcripts isolated in the control condition will be released and, possibly, translated. Thus, we will not pull them down on the beads. Note in this middle panel that there is an empty dotted white box. This box indicates the absence of Ro-released FMRP transcripts (B) because they were released upon injection of Ro-25-6981. In order to recover the Ro-25-6981-released FMRP transcripts (B), we need to subtract out population A from the control condition (which is the union of A and B). Right: Finally, in the Ro-25-6981+Rapa condition, there is a subpopulation within the Ro-25-6981-released transcripts that is also mTORC1-dependent which we can recover by inhibiting mTORC1 with rapamycin (C). Thus, the Ro-25-6981+Rapa condition is the union of the sets A and C. By subtracting out A from this union, we can isolate C. Note that this model is idealized in that we are likely to recover transcripts outside these boxes including transcripts pulled down and sequenced that were recovered due to experimental noise or chosen significant cutoffs. In this way, the true representation from the experiment is actually a three-way Venn diagram rather than an Euler diagram as depicted here. However, for our purposes in this paper we have focused only on the portions of this Venn diagram that are subsets of each other.

**Fig. S2. A**. Correlation of RIP-Seq libraries indicating high correlation between targets. KO libraries were used to determine nonspecific interactions with the FMRP antibody. Briefly, the KO counts were used to generate a WT-to-KO fold change ratio between all libraries to distinguish FMRP targets from FMRP non-targets. This cutoff was used to filter that we used to distinguish signal from background. More detail is given in the methods as well as in Figure S3. Note that SAL_KO1 is correlated against itself because there is only one replicate. This is due to the fact that not RNA was recovered from these beads to amplify for sequencing indicating low background binding for these libraries (see methods for more detail). This and the following figures indicate the reliability and validity of our sequencing approach. **B**. Principal component analysis (PCA) of observations in the RIP-Seq libraries. Color shades indicate WT (dark shade) or KO (light shade) libraries. Here the PCA shows that replicates within the same library cluster together indicating similarity. Furthermore, WT libraries typically cluster closer together than KO libraries as expected. **C**. Euclidean distance matrix shows high similarity between RIP-seq libraries using the complete linkage method for clustering. Similar to figure S2B, the distance matrix also shows that WT libraries are more similar to other WT libraries (the same is true for the KO libraries). Furthermore, replicates within libraries are also more similar to each other than to replicates within other libraries. **D**. Normalized counts (scaled between 0 to 100) for FMRP targets and non-targets (list derived from (22)) across treatments. These targets (purple) and non-targets (grey) are examples that are well-characterized in the literature so they can be used to assess the validity of the sequencing experiment. As expected, non-targets typically had extremely low counts compared to targets indicating that few were pulled down on the beads and resemble background binding. Furthermore, counts for targets generally decrease in the KO condition as expected since they should not be pulled down in the absence of FMRP. **E**. Relative fold-enrichment as determined by real-time qPCR relative to 10% input control ($\Delta\Delta Ct=2^{-(Ct\ FMRP\ RIP-Ct\ IgG\ RIP)-(Ct\ FMRP\ input-Ct\ IgG\ input)}$). FMRP binds to GABABR1 and GABABR2 (known FMRP targets) in Control or Ro-25-6981 treated animals but not in Fmr1 KO animals from RNA derived from RIP of FMRP. This result is expected and indicates the validity of using the KO animals to assess nonspecific binding.

**Fig. S3. A**. Top: Distribution of FMRP targets from other datasets compared to our data. The purpose of this comparison is to determine if our filtered data agrees with past FMRP target data sets in the literature. Each position along the X-axis represents a gene found in the Saline RIP-seq library. The gene with the highest count has a position near the origin while the gene with the lowest count has a position at the other end of the axis. The Y-axis indicates a data from the Ascano PAR-CLIP data (A-PAR), Ascano RIP-seq data (A-RIP), Darnell data (D), Brown data (B), or Miyashiro data. See supplemental methods for data citations and details. Consensus data sets indicate overlap between the above data (e.g. B_D_A-PAR indicates the intersection between the Brown, Darnell, and Ascano PAR-CLIP data). These overlap sets were obtained from the supplemental material in Suhl et al. [22]. Dotted red line indicates cutoff that was used in our data. Everything to the left of the line is considered a putative FMRP target and everything to the right is considered a non-target. Bottom: Line graph indicating the percent of consensus FMRP targets (excluding the Ascano and Miyashiro data) that overlap with our data for each position in the corresponding graph above. Dotted red line indicates the same as in A. Top. For all putative FMRP targets (left of dotted red line), 95% of consensus FMRP targets are captured. **B**. WT-to-KO fold enrichment for genes in our data that overlap with a data set indicated on the X-axis. Non-targets were obtained from [14]. Dotted red line indicates the average bottom quartile for all FMRP target data sets. According to this cutoff, everything above the line is a putative target and everything below is considered background binding. Statistics: ***, $p < 0.001$; Student's t-test. **C**. Scatterplot showing the relationship between the WT and KO counts in the $\log_2$ control RIP-Seq library. Cutoff between "signal" or "background" determined by univariate K-means. **D**. Univariate K-means for $\log_2$ control counts. The two clusters generated (signal, green; background, red) are overlaid on a histogram of the full saline distribution (grey). The max value from the background cluster was chosen as the cutoff point for the distinction between signal and background. This cutoff was averaged with the cutoff described in S3B to distinguish signal from background counts. **E**. Boxplot representing data summarized in Table S1.

**Fig. S4. A.** MA plots showing distribution of counts by fold-change for each of the $\log_2$ transformed fold-change conditions under consideration. Non-significant genes are shown in grey according to an FDR cutoff of 0.1. **B.** Volcano plots for the fold-change conditions under consideration. Non-significant genes are shown in grey according to an FDR cutoff of 0.1. **C.** Clustered dendrogram for each RIP-seq library after regularized log (rlog) transformation. Scale represents rlog transformed raw counts. Clustering performed using the average linkage method and Euclidean distance. Colored box on the right side correspond to GO biological process clusters named below the chart. GO clustering performed with DAVID. **D.** Same scatterplot from Fig. 3B with non-transsynaptic signaling mRNA added in (blue circles). Transsynaptic signaling mRNA shown in red circles. Top left quadrant indicates mRNA translated when mTORC1 is ON and bottom left quadrant indicates mRNA repressed when mTORC1 is ON.

**Fig. S5. A.** Quantification of rat hippocampal neurons treated with Ro-25-6981 (NR2B antagonist) or Ro-25-6981+Baclofen. The postsynaptic marker phalloidin increases with both treatments, but only Ro-25-6981+Baclofen significantly increases synapsin. Scale=1.5 µm. Bars represent mean±SEM. *=p<0.05, ****=p<0.0001. **B.** Representative images of treated WT CA1 stratum radiatum dendrites, including a negative control. White puncta represent proximity-detected PSD-95 and SYN1 proteins by PLA (indicated by yellow arrowheads). **C.** Quantification of the average number of PSD-95/SYN1 PLA puncta per area of CA1 stratum radiatum dendrites, with negative control mapped onto y-axis. Puncta at least 10 µm from the pyramidal layer were analyzed. One-way ANOVA revealed a significant main effect of treatment ($F_{2,52}$=6.79, *p*=0.0024). Newman-Keuls post-hoc revealed that treatment with Ro-25-6981 (4.68±0.78 puncta/µm$^2$ normalized to control, n=23) significantly increased PSD-95/SYN1 PLA puncta 45 minutes after treatment compared to control (1.0±0.11 puncta/µm$^2$ normalized, n=13) and Ro-25-6981+CGP35348 (2.58±0.64 puncta/µm$^2$ normalized, n=19). This increase is blocked with the $GABA_B$R antagonist CGP35348 (no significant difference between control and Ro-25-6981+CGP35348). Scale bar=50 µm. Representative dendrites shown below to demonstrate the puncta are located on dendrites. Scale bar=5 µm. Bars represent mean±SEM. *=*p*<0.05, **=*p*<0.01, ***=*p*<0.001, ****=*p*<0.0001, n.s.=not significant. **D.** Representative images of treated KO CA1 stratum radiatum dendrites. White puncta represent proximity-detected PSD-95 and SYN1 proteins by PLA (indicated by yellow arrowheads). **E.** Quantification of the average number of PSD-95/SYN1 PLA puncta per area of CA1 stratum radiatum dendrites, with negative control mapped onto y-axis. Puncta at least 10 μm from the pyramidal layer were analyzed. Two-tailed t-test ($t_{21}$=2.235, $p$=0.0364) revealed that CGP35348 (0.61±0.15 puncta/$\mu m^2$ normalized to control, n=10) significantly decreased in PSD-95/SYN1 PLA puncta 45 minutes after treatment compared to control (1.0±0.09 puncta/$\mu m^2$ normalized, n=13). Scale bar=50 μm. Representative dendrites shown below to demonstrate the puncta are located on dendrites. Scale bar=5 μm. Bars represent mean±SEM. *=$p$<0.05, **=$p$<0.01, ***=$p$<0.001, ****=$p$<0.0001, n.s.=not significant. **F.** Treatment with CGP35348 does not significantly affect performance in the splash test (see [59] for FST data) for *Fmr1* KO mice (control: 1.1±0.7 times per minute, n=6; CGP35348: 0.6±0.3 times per minute, n=6; $t_{10}$=0.6098, $p$=0.5556).
