## Supplementary figures and images for "Role of FMRP in rapid antidepressant effects and synapse regulation"

### Supplemental Figures 1-5

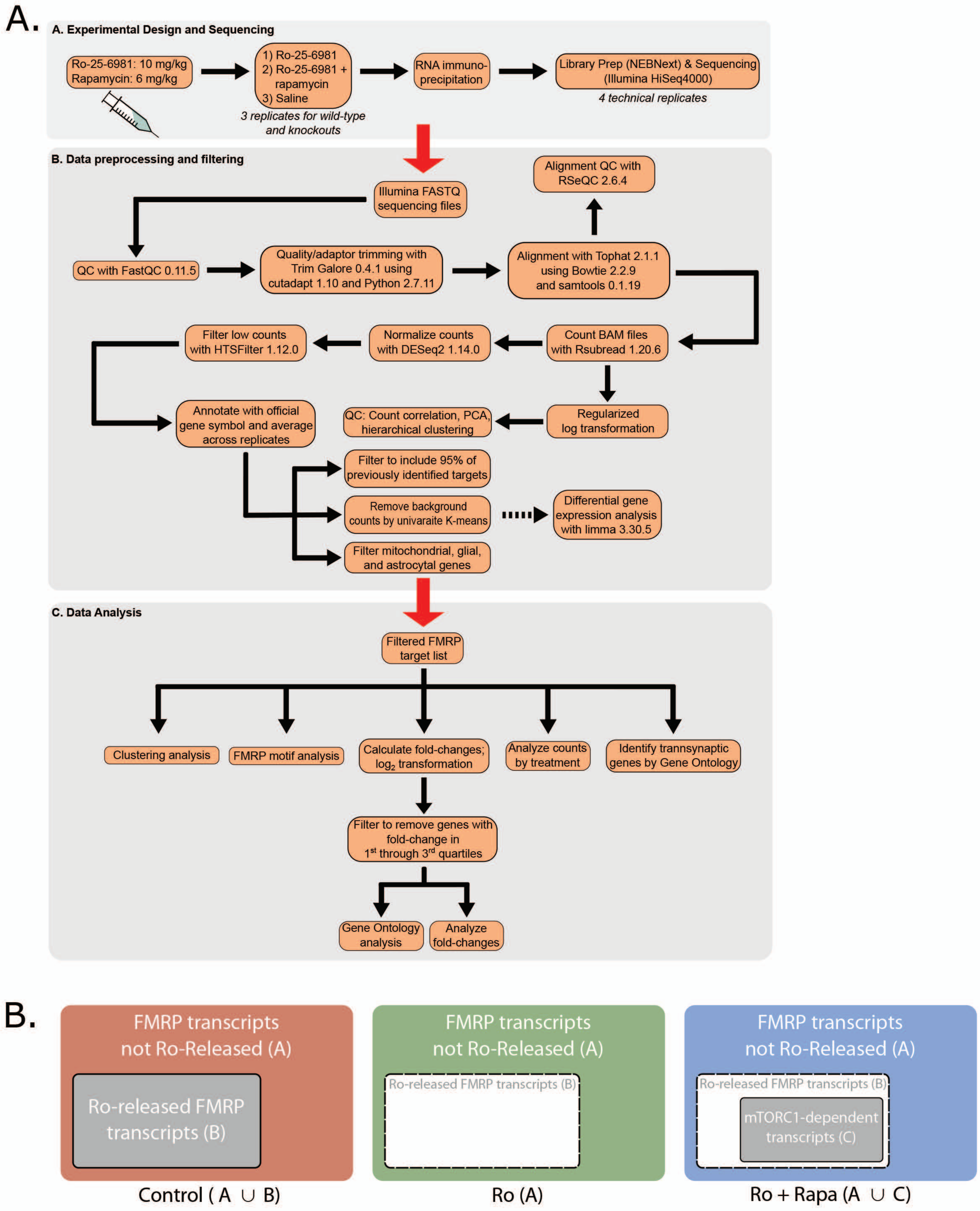

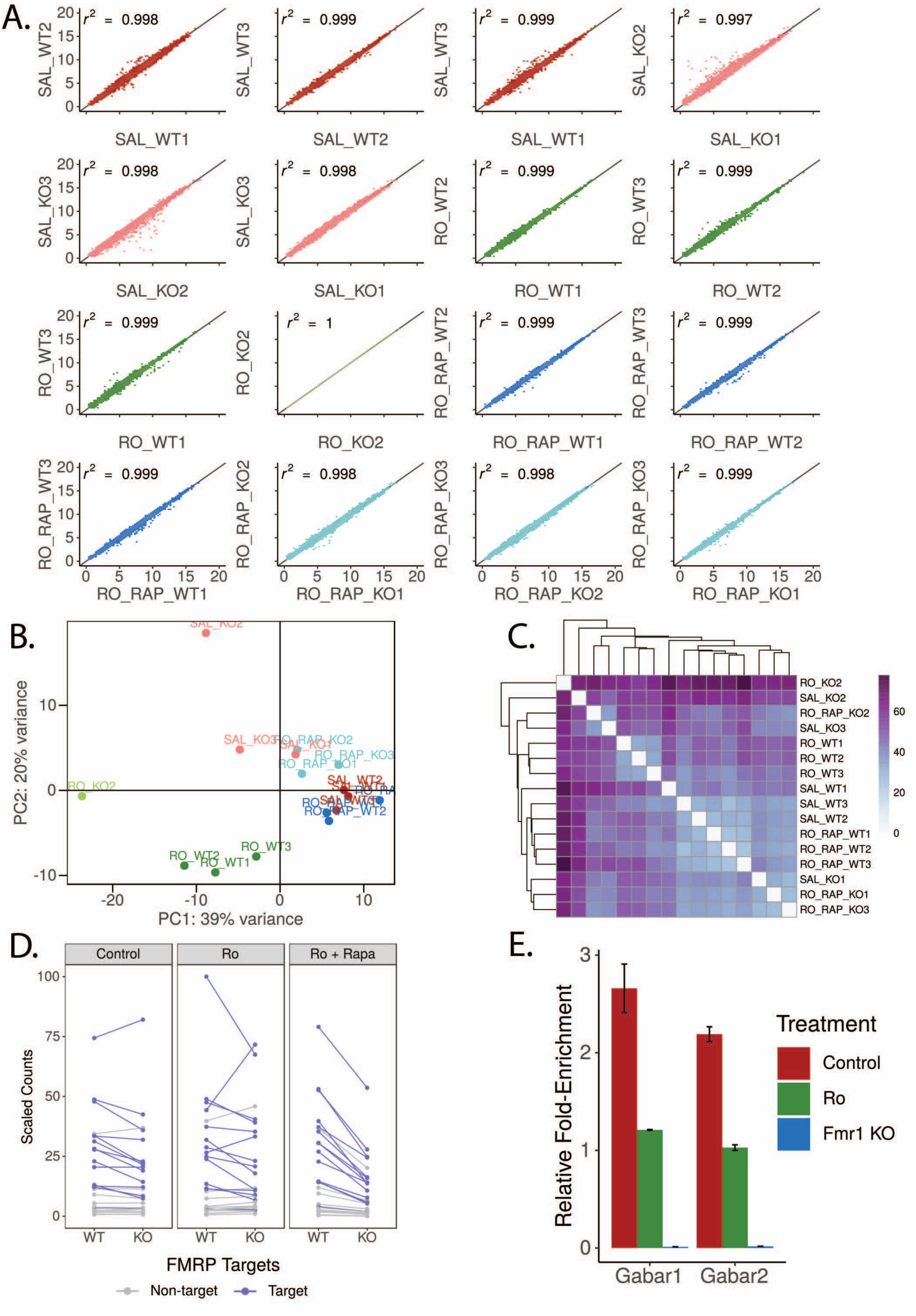

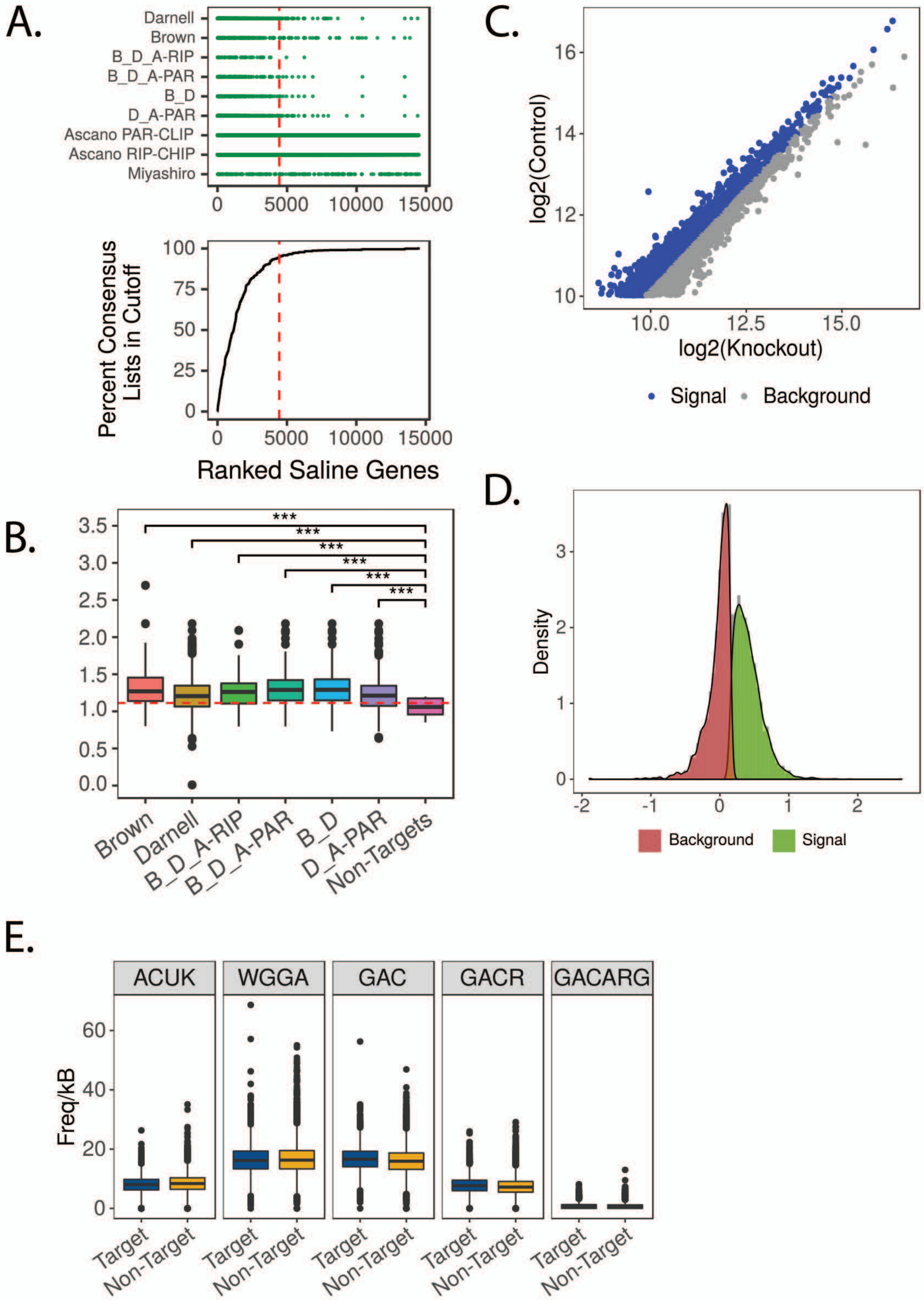

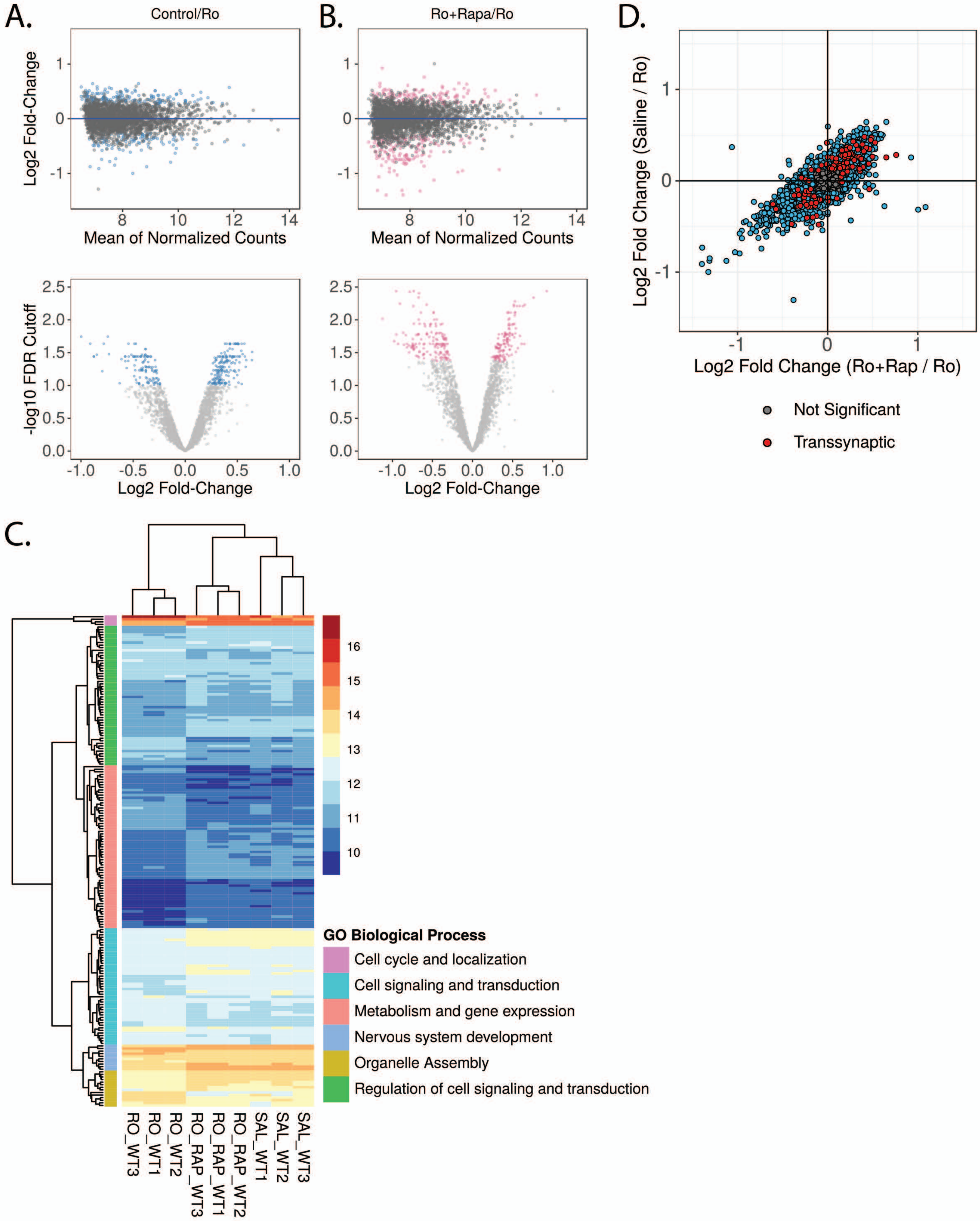

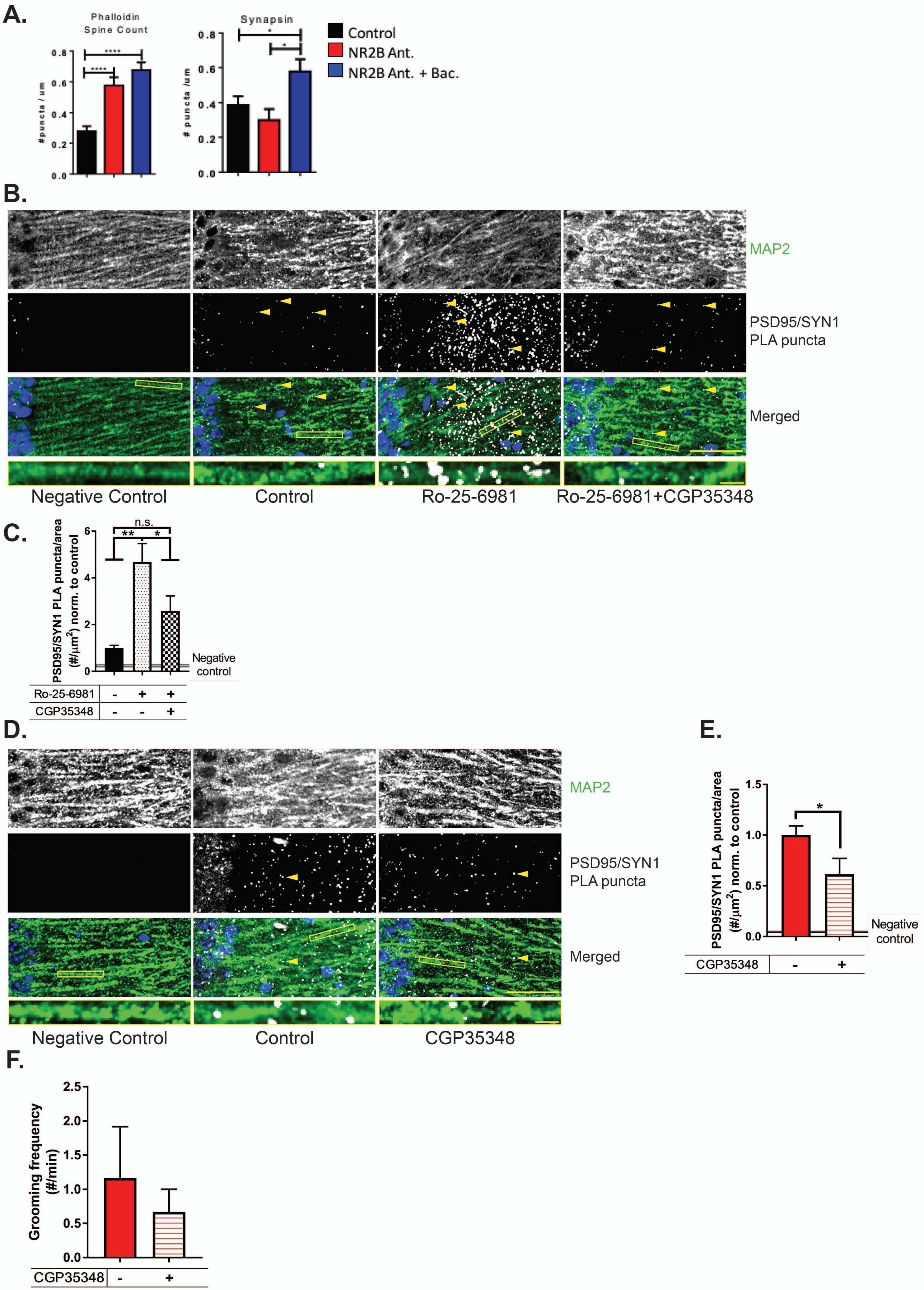
