## Supplemental Table 1 for "Role of FMRP in rapid antidepressant effects and synapse regulation"

**Table S1 - RNA mass recovered and RIN values for RNA immunoprecipitation of FMRP targets.** BioAnalyzer readings for RNA recovered in FMRP RIP-Seq experiment.

| Treatment | Wild-type (Replicate) | | | Knockout (replicate) | | |
|---|---|---|---|---|---|---|
| | 1 | 2 | 3 | 1 | 2 | 3 |
| **RNA mass recovered (ng)** | | | | | | |
| Ro-25-6981 | 36 | 67 | 48 | 3 | 5 | 3 |
| Ro-25-6981 + Rapamycin | 56 | 101 | 75 | 47 | 48 | 102 |
| Control | 54 | 326 | 139 | 29 | 37 | 35 |
| **RNA integrity number (RIN)** | | | | | | |
| Ro-25-6981 | 9 | 8.6 | 9.1 | 1 | 8.4 | 1.1 |
| Ro-25-6981 + Rapamycin | 8.6 | 8.8 | 8.8 | 9 | 7.2 | 8.9 |
| Control | 8.6 | 8.3 | 6.7 | 8.7 | 8.8 | 8.3 |
