## Supplemental Table 2 for "Role of FMRP in rapid antidepressant effects and synapse regulation"

### Table S2 - RIP-seq library and alignment data

| Library | Length (nt) | Input_Reads | Mapping_Rate | Multiple_Alignments |
|---|---|---|---|---|
| RO_KO2_T1 | 20-51 | 436,922 | 16.10% | 15.60% |
| RO_KO2_T2 | 20-51 | 399,555 | 16.10% | 15.50% |
| RO_KO2_T3 | 20-51 | 4,787,614 | 16.60% | 15.00% |
| RO_KO2_T4 | 20-51 | 3,990,679 | 16.40% | 15.10% |
| RO_RAP_KO1_T1 | 20-51 | 8,253,643 | 72.90% | 4.40% |
| RO_RAP_KO1_T2 | 20-51 | 8,140,014 | 73.00% | 4.40% |
| RO_RAP_KO1_T3 | 20-51 | 9,524,091 | 73.50% | 4.40% |
| RO_RAP_KO1_T4 | 20-51 | 8,983,390 | 73.20% | 4.40% |
| RO_RAP_KO2_T1 | 20-51 | 1,539,793 | 76.00% | 3.60% |
| RO_RAP_KO2_T2 | 20-51 | 1,447,787 | 76.00% | 3.60% |
| RO_RAP_KO2_T3 | 20-51 | 5,235,945 | 75.40% | 3.60% |
| RO_RAP_KO2_T4 | 20-51 | 4,919,303 | 75.30% | 3.60% |
| RO_RAP_KO3_T1 | 20-51 | 8,735,516 | 73.30% | 4.00% |
| RO_RAP_KO3_T2 | 20-51 | 8,640,851 | 73.30% | 4.00% |
| RO_RAP_KO3_T3 | 20-51 | 230,114 | 73.60% | 4.00% |
| RO_RAP_KO3_T4 | 20-51 | 9,691,348 | 73.30% | 4.00% |
| RO_RAP_WT1_T1 | 20-51 | 9,425,580 | 81.90% | 3.90% |
| RO_RAP_WT1_T2 | 20-51 | 9,329,940 | 82.00% | 3.90% |
| RO_RAP_WT1_T3 | 20-51 | 3,634,272 | 82.50% | 3.90% |
| RO_RAP_WT1_T4 | 20-51 | 2,678,561 | 82.30% | 3.90% |
| RO_RAP_WT2_T1 | 20-51 | 9,389,927 | 86.50% | 4.30% |
| RO_RAP_WT2_T2 | 20-51 | 9,330,571 | 86.60% | 4.30% |
| RO_RAP_WT2_T3 | 20-51 | 3,847,304 | 86.70% | 4.30% |
| RO_RAP_WT2_T4 | 20-51 | 2,873,233 | 86.60% | 4.30% |
| RO_RAP_WT3_T1 | 20-51 | 7,040,228 | 86.40% | 4.10% |
| RO_RAP_WT3_T2 | 20-51 | 7,030,839 | 86.50% | 4.10% |
| RO_RAP_WT3_T3 | 20-51 | 1,221,288 | 86.70% | 4.10% |
| RO_RAP_WT3_T4 | 20-51 | 504,767 | 86.70% | 4.10% |
| RO_WT1_T1 | 20-51 | 8,568,455 | 39.00% | 5.10% |
| RO_WT1_T2 | 20-51 | 8,499,710 | 39.00% | 5.00% |
| RO_WT1_T3 | 20-51 | 5,282,085 | 40.10% | 5.00% |
| RO_WT1_T4 | 20-51 | 4,971,366 | 39.90% | 5.00% |
| RO_WT2_T1 | 20-51 | 973,892 | 60.30% | 5.60% |
| RO_WT2_T2 | 20-51 | 915,550 | 60.40% | 5.60% |
| RO_WT2_T3 | 20-51 | 9,052,870 | 61.30% | 5.50% |
| RO_WT2_T4 | 20-51 | 8,498,721 | 61.00% | 5.60% |
| RO_WT3_T1 | 20-51 | 7,787,301 | 55.40% | 5.00% |
| RO_WT3_T2 | 20-51 | 7,826,769 | 55.40% | 5.00% |
| RO_WT3_T3 | 20-51 | 5,785,648 | 56.40% | 4.90% |
| RO_WT3_T4 | 20-51 | 4,770,198 | 56.10% | 4.90% |
| SAL_KO1_T2 | 20-51 | 3,360,188 | 62.30% | 4.30% |
| SAL_KO1_T1 | 20-51 | 3,511,776 | 62.20% | 4.40% |
| SAL_KO1_T3 | 20-51 | 975,474 | 63.20% | 4.30% |
| SAL_KO1_T4 | 20-51 | 329,254 | 63.00% | 4.30% |
| SAL_KO2_T1 | 20-51 | 7,887,843 | 46.10% | 4.70% |
| SAL_KO2_T2 | 20-51 | 7,762,342 | 46.00% | 4.70% |
| SAL_KO2_T3 | 20-51 | 6,240,194 | 47.10% | 4.60% |
| SAL_KO2_T4 | 20-51 | 5,855,520 | 46.90% | 4.60% |
| SAL_KO3_T1 | 20-51 | 141,624 | 55.60% | 4.40% |
| SAL_KO3_T2 | 20-51 | 38,776 | 55.70% | 4.40% |
| SAL_KO3_T3 | 20-51 | 8,361,756 | 56.10% | 4.30% |
| SAL_KO3_T4 | 20-51 | 7,898,366 | 55.70% | 4.30% |
| SAL_WT1_T1 | 20-51 | 7,366,745 | 46.40% | 4.60% |
| SAL_WT1_T2 | 20-51 | 7,395,787 | 46.50% | 4.60% |
| SAL_WT1_T3 | 20-51 | 4,801,547 | 47.60% | 4.60% |
| SAL_WT1_T4 | 20-51 | 3,949,727 | 47.20% | 4.60% |
| SAL_WT2_T1 | 20-51 | 7,165,746 | 74.50% | 3.90% |
| SAL_WT2_T2 | 20-51 | 7,629,193 | 74.40% | 3.90% |
| SAL_WT2_T3 | 20-51 | 961,778 | 74.70% | 3.90% |
| SAL_WT2_T4 | 20-51 | 208,678 | 74.50% | 3.90% |
| SAL_WT3_T1 | 20-51 | 6,961,446 | 87.40% | 4.20% |
| SAL_WT3_T2 | 20-51 | 6,952,007 | 87.50% | 4.20% |
| SAL_WT3_T3 | 20-51 | 3,979,856 | 87.70% | 4.20% |
| SAL_WT3_T4 | 20-51 | 2,940,141 | 87.60% | 4.20% |
