## Supplemental Table 3 for "Role of FMRP in rapid antidepressant effects and synapse regulation"

**Table S3 - Statistics for Motif Analysis** - Motif analysis was performed on the filtered FMRP target list and compared to the background target which did not make the cutoff. Motif analysis here replicated analysis performed in Suhl et al., 2014 (see supplemental methods for details and reference). See also Fig. S3E.

| Motif | Freq/kB (Targets) | Freq/kB (Non-Targets) | ρ (Targets) | ρ (Non-Targets) | *t*-test | Wilcoxon-Mann-Whitney Test | Cohen's *d* |
|---|---|---|---|---|---|---|---|
| ACUK | 8.036 | 8.483 | 0.455 | 0.477 | $p < 0.0001$ | $p < 0.0001$ | 0.144 |
| WGGA | 16.531 | 16.666 | 0.2 | 0.201 | $p = 0.222$ | $p = 0.239$ | 0.026 |
| GAC | 16.787 | 16.066 | 0.955 | 0.936 | $p < 0.0001$ | $p < 0.0001$ | 0.162 |
| GACR | 7.905 | 7.444 | 0.222 | 0.214 | $p < 0.0001$ | $p < 0.0001$ | 0.152 |
| GACARG | 0.875 | 0.773 | 0.024 | 0.022 | $p < 0.0001$ | $p < 0.0001$ | 0.115 |
| | **Freq/gene (Targets)** | **Freq/gene (Non-Targets)** | | | | | |
| QFM | 1.17 | 1.074 | — | — | $p < 0.0001$ | $p < 0.0001$ | — |
