## Supplemental Table 4 for "Role of FMRP in rapid antidepressant effects and synapse regulation"

**Table S4 - Fold-changes for significant differentially expressed mRNA** - Significant genes from tables S6 common to both Saline/Ro or Ro+Rapa/Ro.

| | Downregulated | |
|---|---|---|
| | Saline/Ro | Ro+Rapa/Ro |
| Abl2 | -0.43436 | -0.38726416 |
| Aftph | -0.26261 | -0.231876431 |
| Arap2 | -0.35174 | -0.279920205 |
| Arhgap5 | -0.4083 | -0.599667989 |
| Arl4a | -0.37215 | -0.667965952 |
| Birc2 | -0.48585 | -0.372748665 |
| Brk1 | -0.48934 | -0.638627333 |
| Cops2 | -0.48618 | -0.658996644 |
| Cyb5r4 | -0.41355 | -0.303658326 |
| Dlg1 | -0.28423 | -0.220252656 |
| Epc2 | -0.28193 | -0.34399462 |
| Erbin | -0.29841 | -0.520719424 |
| Fam212b | -0.57418 | -0.496884618 |
| Fat3 | -0.3847 | -0.37766726 |
| Fmr1 | -0.30928 | -0.572730162 |
| Fnta | -0.43999 | -0.566148083 |
| Grin2a | -0.47266 | -0.398899063 |
| Hecw2 | -0.40886 | -0.78717654 |
| Hipk2 | -0.44046 | -0.600233671 |
| Irs1 | -0.49388 | -0.446565263 |
| Kcnh1 | -0.40746 | -0.433014636 |
| 8-Mar | -0.33225 | -0.348614242 |
| Mdga2 | -0.49782 | -0.783963377 |
| Mobp | -0.79525 | -0.976740288 |
| Mtmr6 | -0.30382 | -0.263539733 |
| Myo9a | -0.35188 | -0.316236661 |
| Noct | -0.54166 | -0.452055352 |
| Nr3c1 | -0.27353 | -0.301765564 |
| Ntrk3 | -0.42193 | -0.455448392 |
| Pkn2 | -0.36062 | -0.334632449 |
| Pou3f3 | -0.35013 | -0.225108073 |
| Ppp1r16b | -0.34132 | -0.288288243 |
| Prex2 | -0.51595 | -0.450162633 |
| Psmc2 | -0.3155 | -0.47817906 |
| Pum2 | -0.37258 | -0.439313411 |
| Rab11fip2 | -0.24175 | -0.375494441 |
| Rb1cc1 | -0.36528 | -0.270752652 |
| Rlf | -0.48715 | -0.486156664 |
| Rn7sk | -1.6027 | -1.719252197 |
| Rnf19a | -0.38029 | -0.349158498 |
| Rpl12 | -0.99781 | -1.323264047 |
| Rpl15 | -0.42498 | -0.56902287 |
| Rpl17 | -0.44159 | -0.670744785 |
| Rpl18a | -0.54644 | -0.952063778 |
| Rpl31 | -0.78155 | -1.027366335 |
| Rpl37 | -0.44237 | -0.905967611 |
| Rpl38 | -0.91097 | -1.394633057 |
| Rps11 | -0.47261 | -0.679761807 |
| Rps17 | -0.57772 | -0.975559148 |
| Rps21 | -0.87629 | -1.311337622 |
| Rps23 | -0.84822 | -1.30905756 |
| Rps27a | -0.73107 | -1.391710613 |
| Rps3 | -0.62908 | -0.950790151 |
| Rps7 | -0.87653 | -1.122987432 |
| Rps9 | -0.56176 | -0.702194291 |
| Sacs | -0.4779 | -0.400056484 |
| Scn1a | -0.31725 | -0.333170908 |
| Sema7a | -0.29155 | -0.301598062 |
| Setx | -0.36416 | -0.338488765 |
| Slitrk4 | -0.39474 | -0.683279034 |
| Sptbn1 | -0.42231 | -0.324099459 |
| Tmem178b | -0.37667 | -0.489810452 |
| Tpt1 | -0.35141 | -0.643622346 |
| Trps1 | -0.54409 | -0.671985571 |
| Tsc1 | -0.31237 | -0.236180563 |
| Ttbk2 | -0.39241 | -0.387571962 |
| Tug1 | -0.38375 | -0.485768197 |
| Usf3 | -0.61145 | -0.680321449 |
| Wnk1 | -0.46506 | -0.436614545 |
| X26105071 | -0.38043 | -0.312179716 |
| X5031439( | -0.29995 | -0.326871562 |
| Zbtb20 | -0.74257 | -0.734619508 |
| Zbtb38 | -0.36291 | -0.363983976 |
| Zbtb41 | -0.37785 | -0.402111256 |
| Zfp644 | -0.38135 | -0.310148155 |
| Zfp651 | -0.28536 | -0.279722796 |
| Zzz3 | -0.41395 | -0.362745899 |

| | Upregulated | |
|---|---|---|
| | Saline/Ro | Ro+Rapa/Ro |
| Acot7 | 0.308754 | 0.30733071 |
| Actb | 0.485217 | 0.404576241 |
| Actg1 | 0.331121 | 0.203542298 |
| Adprh | 0.422655 | 0.312890527 |
| Aes | 0.64283 | 0.430016665 |
| Agpat1 | 0.449145 | 0.492469249 |
| Aldoa | 0.386551 | 0.516077238 |
| Anxa7 | 0.400459 | 0.312303192 |
| Aplp1 | 0.426868 | 0.28991054 |
| Arhgdia | 0.463436 | 0.436234853 |
| Arl8a | 0.568026 | 0.390414739 |
| Blcap | 0.336392 | 0.246419777 |
| C1ql3 | 0.323339 | 0.432342006 |
| C2cd2l | 0.51119 | 0.528047372 |
| Calm3 | 0.572607 | 0.434583938 |
| Cbx4 | 0.325703 | 0.435240512 |
| Cd81 | 0.378516 | 0.369640581 |
| Cdh13 | 0.454846 | 0.508994617 |
| Cdk5r2 | 0.392782 | 0.279017337 |
| Clu | 0.514066 | 0.56628776 |
| Cops7a | 0.392759 | 0.328383935 |
| Cplx1 | 0.473379 | 0.488131978 |
| Cpne4 | 0.465008 | 0.590657457 |
| Cpne6 | 0.585619 | 0.541719438 |
| Ctbp1 | 0.283172 | 0.286975439 |
| Ctsf | 0.290717 | 0.263852901 |
| Cyp46a1 | 0.43233 | 0.446770894 |
| Ddost | 0.29122 | 0.397378239 |
| Dpp6 | 0.298406 | 0.300889987 |
| Emc10 | 0.405392 | 0.430753766 |
| Eno1 | 0.381889 | 0.31945415 |
| Epn1 | 0.407066 | 0.475448191 |
| Fads1 | 0.398065 | 0.384402354 |
| Fads2 | 0.318479 | 0.285916534 |
| Fam19a2 | 0.538283 | 0.296555418 |
| Fam219a | 0.351256 | 0.282277219 |
| Fjx1 | 0.484422 | 0.271973854 |
| Gpr37l1 | 0.333059 | 0.280281668 |
| Grina | 0.401184 | 0.346157798 |
| Gsk3a | 0.501088 | 0.437092674 |
| Jph4 | 0.390843 | 0.31750816 |
| Jund | 0.525758 | 0.5206927 |
| Lamp1 | 0.272765 | 0.253915197 |
| Lancl1 | 0.322424 | 0.323845294 |
| Ldb1 | 0.316352 | 0.265428143 |
| Lmo3 | 0.500112 | 0.334366657 |
| Map2k1 | 0.401794 | 0.410391411 |
| Marcks | 0.399603 | 0.435880446 |
| Mcrs1 | 0.284479 | 0.271548673 |
| Med24 | 0.342512 | 0.327153456 |
| Mfge8 | 0.500037 | 0.611335586 |
| Mgrn1 | 0.33748 | 0.325533556 |
| Ncdn | 0.344865 | 0.459426093 |
| Ncs1 | 0.348659 | 0.465042107 |
| Nov | 0.551015 | 0.328505004 |
| Nptxr | 0.517144 | 0.463119151 |
| Olfm1 | 0.307515 | 0.340798496 |
| Olfm2 | 0.321426 | 0.270471987 |
| Paqr4 | 0.380566 | 0.313970439 |
| Parp6 | 0.394949 | 0.368045025 |
| Phyhip | 0.429898 | 0.448644571 |
| Plpp3 | 0.503348 | 0.527024401 |
| Pnkd | 0.359615 | 0.426269139 |
| Ppp2r1a | 0.317487 | 0.387817637 |
| Ppp2r5b | 0.399102 | 0.386793792 |
| Prkaca | 0.481174 | 0.40860635 |
| Prkacb | 0.308283 | 0.30161676 |
| Prkar1b | 0.333412 | 0.307687245 |
| Psap | 0.298305 | 0.312994441 |
| Psd | 0.312573 | 0.308095426 |
| Rab3a | 0.429977 | 0.284181213 |
| Rgs12 | 0.270844 | 0.410915502 |
| Rnf187 | 0.333245 | 0.272394669 |
| Rnps1 | 0.397431 | 0.34691471 |
| Rogdi | 0.452137 | 0.38113819 |
| Rpn2 | 0.306785 | 0.456306734 |
| Rusc1 | 0.416991 | 0.373213559 |
| Scamp5 | 0.300056 | 0.191400719 |
| Scn1b | 0.427973 | 0.308969198 |
| Scn3b | 0.382086 | 0.460733266 |
| 5-Sep | 0.454082 | 0.555874155 |
| Sh3bp1 | 0.346244 | 0.282408875 |
| Slc17a7 | 0.362953 | 0.248134645 |
| Slc1a4 | 0.344575 | 0.524723939 |
| Smap2 | 0.363149 | 0.529263539 |
| Smarcd3 | 0.47802 | 0.485239236 |
| Smpd1 | 0.327084 | 0.302580945 |
| Snap47 | 0.300472 | 0.257941315 |
| Sqle | 0.263412 | 0.293035542 |
| Strn4 | 0.253189 | 0.296160023 |
| Stum | 0.314686 | 0.268020843 |
| Syngr1 | 0.266272 | 0.231613626 |
| Syngr3 | 0.332247 | 0.373986173 |
| Syt4 | 0.293864 | 0.23765091 |
| Thra | 0.465485 | 0.515716822 |
| Timp2 | 0.510016 | 0.591468071 |
| Tmem151a | 0.348289 | 0.349049986 |
| Tmem63b | 0.290547 | 0.277783787 |
| Tom1l2 | 0.222574 | 0.229386544 |
| Trappc12 | 0.253776 | 0.262509016 |
| Tspan5 | 0.342468 | 0.494832331 |
| Tspan7 | 0.442808 | 0.401724446 |
| Tuba4a | 0.231129 | 0.291203042 |
| Tubb5 | 0.353897 | 0.407234403 |
| Ubb | 0.282701 | 0.344682459 |
| Vapa | 0.393961 | 0.42843086 |
| Vps26b | 0.27882 | 0.283017802 |
| Wbp2 | 0.457144 | 0.404834063 |
| Wfs1 | 0.641086 | 0.647777549 |
| Wnt7b | 0.520782 | 0.422223131 |
| Wsb2 | 0.283162 | 0.243849259 |
| X2900011( | 0.356283 | 0.284083861 |

| | Oppositely regulated | |
|---|---|---|
| | Saline/Ro | Ro+Rapa/Ro |
| Dbp | 0.3688728 | -1.0617925 |
