## Supplemental Table 5 for "Role of FMRP in rapid antidepressant effects and synapse regulation"

### Table S5 - DGE anaylsis for FMRP targets using Limma

| | Control/Ro | | | | | | | Ro+Rapa/Ro | | | | | | | |
|---|---|---|---|---|---|---|---|---|---|---|---|---|---|---|---|
| | logFC | AveExpr | t | P.Value | adj.P.Val | B | Threshold | logFC | AveExpr | t | P.Value | adj.P.Val | B | Threshold | Transsynaptic? |
| Aak1 | -0.01215 | 9.631335 | -0.11382 | 0.911411 | 0.953049 | -6.37089 | Non-Significant | 0.051057 | 9.631335 | 0.475817 | 0.643434 | 0.777229 | -6.54411 | Non-Significant | FALSE |
| Aamp | -0.06605 | 8.290475 | -0.40097 | 0.696058 | 0.832906 | -6.2501 | Non-Significant | -0.08148 | 8.290475 | -0.50014 | 0.626745 | 0.76369 | -6.50357 | Non-Significant | FALSE |
| Aar2 | 0.124915 | 6.775716 | 1.378999 | 0.195055 | 0.402742 | -5.2654 | Non-Significant | 0.173432 | 6.775716 | 1.969055 | 0.074424 | 0.200711 | -4.72905 | Non-Significant | FALSE |
| Aars | 0.003939 | 8.660136 | 0.054361 | 0.957614 | 0.976205 | -6.35592 | Non-Significant | 0.098426 | 8.660136 | 1.367347 | 0.198573 | 0.372133 | -5.73005 | Non-Significant | FALSE |
| Aatk | -0.14149 | 9.895432 | -1.64787 | 0.127375 | 0.322411 | -5.0956 | Non-Significant | -0.06884 | 9.895432 | -0.79634 | 0.44254 | 0.619187 | -6.33214 | Non-Significant | FALSE |
| Abca2 | -0.07496 | 9.326768 | -0.74736 | 0.470387 | 0.673419 | -6.08747 | Non-Significant | -0.03079 | 9.326768 | -0.30631 | 0.765039 | 0.862279 | -6.61513 | Non-Significant | FALSE |
| Abce1 | 0.108167 | 7.323844 | 1.162122 | 0.26957 | 0.495137 | -5.59172 | Non-Significant | 0.076529 | 7.323844 | 0.829067 | 0.424541 | 0.606357 | -6.2019 | Non-Significant | FALSE |
| Abcf1 | -0.12678 | 7.054592 | -1.40105 | 0.18854 | 0.396629 | -5.27571 | Non-Significant | 0.166136 | 7.054592 | 1.886099 | 0.085712 | 0.220261 | -4.893 | Non-Significant | FALSE |
| Abcg1 | 0.134646 | 7.400859 | 1.674707 | 0.121911 | 0.317585 | -4.96037 | Non-Significant | 0.041989 | 7.400859 | 0.525178 | 0.609792 | 0.75071 | -6.41973 | Non-Significant | FALSE |
| Abhd12 | 0.20345 | 9.167735 | 2.316563 | 0.040633 | 0.178092 | -4.07023 | Non-Significant | 0.131737 | 9.167735 | 1.498788 | 0.161827 | 0.328527 | -5.56841 | Non-Significant | FALSE |
| Abhd16a | 0.262061 | 7.631207 | 1.727541 | 0.111761 | 0.302477 | -4.91069 | Non-Significant | -0.10372 | 7.631207 | -0.68251 | 0.508915 | 0.675025 | -6.34222 | Non-Significant | FALSE |
| Abhd17a | 0.14183 | 8.683966 | 0.980762 | 0.347609 | 0.568661 | -5.87432 | Non-Significant | -0.06766 | 8.683966 | -0.46951 | 0.647795 | 0.780796 | -6.53948 | Non-Significant | FALSE |
| Abhd2 | 0.154988 | 8.076169 | 1.08398 | 0.301363 | 0.52474 | -5.73094 | Non-Significant | -0.07454 | 8.076169 | -0.52489 | 0.609987 | 0.75071 | -6.4711 | Non-Significant | FALSE |
| Abhd3 | 0.132569 | 6.793738 | 0.903281 | 0.385565 | 0.600429 | -5.77744 | Non-Significant | 0.059315 | 6.793738 | 0.413157 | 0.687363 | 0.810907 | -6.40802 | Non-Significant | FALSE |
| Abhd8 | 0.151776 | 9.324054 | 1.461758 | 0.17154 | 0.380067 | -5.34204 | Non-Significant | 0.29337 | 9.324054 | 2.81536 | 0.016678 | 0.083638 | -3.47074 | Non-Significant | FALSE |
| Abi1 | -0.05796 | 7.928158 | -0.53307 | 0.604494 | 0.772714 | -6.16402 | Non-Significant | -0.11923 | 7.928158 | -1.10709 | 0.291675 | 0.471682 | -5.99009 | Non-Significant | FALSE |
| Abi2 | 0.137565 | 9.092968 | 1.319153 | 0.213689 | 0.429624 | -5.51885 | Non-Significant | 0.07555 | 9.092968 | 0.724433 | 0.483794 | 0.65175 | -6.38936 | Non-Significant | FALSE |
| Abl1 | -0.09777 | 7.302067 | -1.01447 | 0.331967 | 0.555978 | -5.7511 | Non-Significant | -0.14097 | 7.302067 | -1.47867 | 0.167045 | 0.335904 | -5.50349 | Non-Significant | FALSE |
| Abl2 | -0.43436 | 8.563785 | -3.40744 | 0.005781 | 0.075521 | -2.21153 | Significant | -0.38726 | 8.563785 | -3.07016 | 0.010556 | 0.06358 | -3.02407 | Non-Significant | FALSE |
| Ablim2 | -0.01566 | 7.481387 | -0.19893 | 0.845913 | 0.916533 | -6.25683 | Non-Significant | 0.139442 | 7.481387 | 1.793725 | 0.100121 | 0.243397 | -5.07316 | Non-Significant | FALSE |
| Abr | 0.152166 | 10.23824 | 1.313164 | 0.215633 | 0.430324 | -5.52567 | Non-Significant | 0.313748 | 10.23824 | 2.679166 | 0.021295 | 0.096987 | -3.69965 | Non-Significant | FALSE |
| Acat2 | 0.061453 | 7.060279 | 0.467474 | 0.649206 | 0.80008 | -6.11686 | Non-Significant | -0.01393 | 7.060279 | -0.10768 | 0.916174 | 0.958404 | -6.52905 | Non-Significant | FALSE |
| Ache | 0.207881 | 6.734819 | 1.094455 | 0.296942 | 0.519746 | -5.58429 | Non-Significant | 0.10256 | 6.734819 | 0.55042 | 0.592938 | 0.742426 | -6.32466 | Non-Significant | FALSE |
| Acot7 | 0.308754 | 8.106004 | 3.13557 | 0.009389 | 0.093468 | -2.64757 | Significant | 0.307331 | 8.106004 | 3.155961 | 0.009052 | 0.059416 | -2.83734 | Non-Significant | FALSE |
| Acp1 | -0.41894 | 7.022768 | -2.25321 | 0.045433 | 0.18661 | -4.07741 | Non-Significant | -0.73174 | 7.022768 | -3.97579 | 0.002138 | 0.028538 | -1.36838 | Significant | FALSE |
| Acp2 | 0.23561 | 6.791672 | 2.23287 | 0.047087 | 0.190019 | -4.05608 | Non-Significant | 0.03019 | 6.791672 | 0.290721 | 0.776626 | 0.868157 | -6.44883 | Non-Significant | FALSE |
| Acsbg1 | 0.029078 | 8.752955 | 0.277452 | 0.786534 | 0.88316 | -6.32136 | Non-Significant | 0.153145 | 8.752955 | 1.468756 | 0.169667 | 0.338576 | -5.60276 | Non-Significant | FALSE |
| Acsl3 | -0.18152 | 9.008753 | -1.84314 | 0.092158 | 0.270614 | -4.8096 | Non-Significant | -0.10994 | 9.008753 | -1.11848 | 0.286989 | 0.46772 | -6.03037 | Non-Significant | FALSE |
| Acsl5 | -0.07914 | 8.405852 | -0.73435 | 0.477968 | 0.677965 | -6.06391 | Non-Significant | -0.0156 | 8.405852 | -0.14637 | 0.886254 | 0.943694 | -6.63251 | Non-Significant | FALSE |
| Actb | 0.485217 | 10.8146 | 5.766926 | 0.000121 | 0.023095 | 1.511245 | Significant | 0.404576 | 10.8146 | 4.772216 | 0.000565 | 0.016801 | -0.0953 | Significant | FALSE |
| Actg1 | 0.331121 | 10.24807 | 4.44143 | 0.000972 | 0.036195 | -0.49211 | Significant | 0.203542 | 10.24807 | 2.715322 | 0.019958 | 0.094096 | -3.63925 | Non-Significant | FALSE |
| Actn4 | 0.087107 | 7.007508 | 0.582182 | 0.572082 | 0.74979 | -6.04785 | Non-Significant | -0.01378 | 7.007508 | -0.09374 | 0.926984 | 0.964063 | -6.52417 | Non-Significant | FALSE |
| Actr1a | 0.19487 | 8.413027 | 2.286505 | 0.042846 | 0.182624 | -4.08757 | Non-Significant | 0.246693 | 8.413027 | 2.920964 | 0.013797 | 0.075731 | -3.26365 | Non-Significant | FALSE |
| Actr1b | 0.212247 | 9.305414 | 2.151877 | 0.054258 | 0.202654 | -4.33612 | Non-Significant | 0.177667 | 9.305414 | 1.797298 | 0.099525 | 0.24296 | -5.14598 | Non-Significant | FALSE |
| Actr3 | -0.19212 | 9.019496 | -1.79331 | 0.10019 | 0.283027 | -4.88416 | Non-Significant | -0.3535 | 9.019496 | -3.30593 | 0.006925 | 0.051803 | -2.61415 | Non-Significant | FALSE |
| Actr3b | 0.01579 | 8.505662 | 0.122242 | 0.904893 | 0.950114 | -6.34827 | Non-Significant | -0.29724 | 8.505662 | -2.30915 | 0.041168 | 0.142074 | -4.32489 | Non-Significant | FALSE |
| Actr8 | -0.12415 | 7.038995 | -0.99906 | 0.339054 | 0.562899 | -5.72851 | Non-Significant | -0.04394 | 7.038995 | -0.36193 | 0.724205 | 0.834518 | -6.47137 | Non-Significant | FALSE |
| Acvr1b | 0.00836 | 8.817817 | 0.120198 | 0.906475 | 0.950114 | -6.36078 | Non-Significant | -0.09208 | 8.817817 | -1.32821 | 0.210778 | 0.386884 | -5.786 | Non-Significant | FALSE |
| Acvr2a | 0.168363 | 7.228459 | 1.421152 | 0.182759 | 0.389673 | -5.27394 | Non-Significant | 0.133226 | 7.228459 | 1.135703 | 0.280014 | 0.460934 | -5.89318 | Non-Significant | FALSE |
| Adam15 | 0.270028 | 6.829941 | 2.231478 | 0.047202 | 0.190019 | -4.05061 | Non-Significant | 0.319083 | 6.829941 | 2.698774 | 0.020559 | 0.095488 | -3.5275 | Non-Significant | FALSE |
| Adap1 | 0.174927 | 9.187475 | 2.366002 | 0.03723 | 0.170843 | -3.98946 | Non-Significant | 0.146094 | 9.187475 | 1.973768 | 0.073825 | 0.199434 | -4.87761 | Non-Significant | FALSE |
| Adar | -0.11155 | 8.657516 | -0.72361 | 0.484282 | 0.684023 | -6.08465 | Non-Significant | 0.214838 | 8.657516 | 1.406447 | 0.186972 | 0.357396 | -5.68063 | Non-Significant | FALSE |
| Adarb1 | -0.03859 | 9.383636 | -0.44121 | 0.667535 | 0.811466 | -6.27416 | Non-Significant | -0.02668 | 9.383636 | -0.30426 | 0.766558 | 0.862539 | -6.6159 | Non-Significant | FALSE |
| Adcy2 | 0.015708 | 9.088242 | 0.221935 | 0.828396 | 0.909571 | -6.34679 | Non-Significant | 0.085113 | 9.088242 | 1.202506 | 0.254198 | 0.434424 | -5.93592 | Non-Significant | FALSE |
| Adcy3 | 0.016425 | 7.85324 | 0.161624 | 0.874507 | 0.932668 | -6.28937 | Non-Significant | 0.282625 | 7.85324 | 2.828608 | 0.016286 | 0.0826 | -3.40346 | Non-Significant | FALSE |
| Adcy5 | -0.19065 | 10.04124 | -1.70545 | 0.115908 | 0.308551 | -5.01307 | Non-Significant | -0.12615 | 10.04124 | -1.12018 | 0.286294 | 0.467213 | -6.02344 | Non-Significant | FALSE |
| Adcy8 | 0.111843 | 6.953171 | 0.699999 | 0.49834 | 0.693636 | -5.95901 | Non-Significant | 0.081729 | 6.953171 | 0.52215 | 0.61183 | 0.751597 | -6.37522 | Non-Significant | FALSE |
| Add1 | 0.010009 | 9.594093 | 0.137268 | 0.893278 | 0.942182 | -6.36741 | Non-Significant | 0.095697 | 9.594093 | 1.30575 | 0.218058 | 0.3943 | -5.81418 | Non-Significant | FALSE |
| Add3 | 0.18116 | 7.688971 | 2.01394 | 0.068907 | 0.232996 | -4.48487 | Non-Significant | 0.129481 | 7.688971 | 1.450343 | 0.174633 | 0.345653 | -5.56479 | Non-Significant | FALSE |
| Adgra1 | 0.139371 | 7.316912 | 0.894662 | 0.38996 | 0.602604 | -5.84096 | Non-Significant | 0.404193 | 7.316912 | 2.636076 | 0.023004 | 0.101212 | -3.68549 | Non-Significant | FALSE |
| Adgrb1 | 0.114092 | 9.683646 | 1.280156 | 0.226603 | 0.444899 | -5.5686 | Non-Significant | 0.308843 | 9.683646 | 3.439489 | 0.005462 | 0.046269 | -2.37424 | Non-Significant | FALSE |
| Adgrb2 | -0.11278 | 10.31536 | -0.91269 | 0.380804 | 0.597069 | -5.94724 | Non-Significant | 0.053387 | 10.31536 | 0.427476 | 0.67721 | 0.802464 | -6.5559 | Non-Significant | FALSE |
| Adgrl1 | -0.16352 | 10.85145 | -1.3054 | 0.218172 | 0.432914 | -5.51971 | Non-Significant | -0.11761 | 10.85145 | -0.92967 | 0.372325 | 0.557925 | -6.1922 | Non-Significant | FALSE |
| Adgrl2 | -0.12894 | 8.047491 | -1.03149 | 0.324267 | 0.548917 | -5.78491 | Non-Significant | -0.22172 | 8.047491 | -1.79084 | 0.100604 | 0.243817 | -5.11532 | Non-Significant | FALSE |
| Adgrl3 | -0.15156 | 9.461798 | -1.76019 | 0.105874 | 0.2908 | -4.9363 | Non-Significant | -0.13966 | 9.461798 | -1.61714 | 0.133896 | 0.28994 | -5.40939 | Non-Significant | FALSE |
| Adh5 | -0.07392 | 6.845644 | -0.44473 | 0.665062 | 0.80993 | -6.10252 | Non-Significant | -0.15575 | 6.845644 | -0.95875 | 0.358108 | 0.54235 | -6.04565 | Non-Significant | FALSE |
| Adipor1 | 0.07272 | 7.206457 | 0.764635 | 0.46044 | 0.666327 | -5.94737 | Non-Significant | 0.054184 | 7.206457 | 0.576377 | 0.575864 | 0.731924 | -6.37238 | Non-Significant | FALSE |
| Adipor2 | -0.0164 | 7.357793 | -0.19654 | 0.847742 | 0.916533 | -6.25081 | Non-Significant | -0.06335 | 7.357793 | -0.76544 | 0.45998 | 0.633981 | -6.26427 | Non-Significant | FALSE |
| Adora2a | 0.266264 | 7.110145 | 1.931165 | 0.079397 | 0.249268 | -4.54736 | Non-Significant | 0.334455 | 7.110145 | 2.459533 | 0.031532 | 0.122407 | -3.96187 | Non-Significant | TRUE |
| Adprh | 0.422655 | 6.618995 | 3.858235 | 0.002619 | 0.051849 | -1.35504 | Significant | 0.312891 | 6.618995 | 2.913091 | 0.013993 | 0.076321 | -3.13056 | Non-Significant | FALSE |
| Adra2c | -0.14014 | 7.848135 | -1.39693 | 0.189742 | 0.396946 | -5.36177 | Non-Significant | -0.03417 | 7.848135 | -0.34504 | 0.736513 | 0.842538 | -6.54675 | Non-Significant | FALSE |
| Aes | 0.64283 | 8.56806 | 5.037658 | 0.000369 | 0.030902 | 0.449722 | Significant | 0.430017 | 8.56806 | 3.379617 | 0.006074 | 0.048848 | -2.45731 | Non-Significant | FALSE |
| Aff3 | -0.20151 | 7.217015 | -1.53801 | 0.152058 | 0.355954 | -5.12236 | Non-Significant | 0.156771 | 7.217015 | 1.225112 | 0.245901 | 0.425402 | -5.80514 | Non-Significant | FALSE |
| Aff4 | -0.18645 | 9.522132 | -1.6943 | 0.118055 | 0.31068 | -5.03113 | Non-Significant | -0.09107 | 9.522132 | -0.82413 | 0.427223 | 0.609213 | -6.31273 | Non-Significant | FALSE |
| Aftph | -0.26261 | 9.003654 | -3.27633 | 0.0073 | 0.084431 | -2.44468 | Significant | -0.23188 | 9.003654 | -2.89995 | 0.014327 | 0.077199 | -3.32688 | Non-Significant | FALSE |
| Agap1 | -0.14758 | 8.648417 | -1.3061 | 0.217944 | 0.432914 | -5.52259 | Non-Significant | -0.06377 | 8.648417 | -0.5687 | 0.580887 | 0.73482 | -6.48829 | Non-Significant | FALSE |
| Agap2 | -0.06475 | 11.50706 | -0.74806 | 0.469979 | 0.673419 | -6.04613 | Non-Significant | 0.062309 | 11.50706 | 0.710721 | 0.491925 | 0.658749 | -6.34832 | Non-Significant | FALSE |
| Agap3 | 0.087696 | 8.594066 | 1.012488 | 0.332872 | 0.557144 | -5.82805 | Non-Significant | 0.400905 | 8.594066 | 4.669191 | 0.000668 | 0.017878 | -0.26582 | Significant | FALSE |
| Aggf1 | -0.25313 | 7.944213 | -3.32257 | 0.006723 | 0.081098 | -2.3255 | Significant | -0.03699 | 7.944213 | -0.49345 | 0.631316 | 0.767437 | -6.48832 | Non-Significant | FALSE |
| Agl | -0.23432 | 6.92779 | -2.02352 | 0.06778 | 0.23154 | -4.41108 | Non-Significant | -0.34789 | 6.92779 | -3.07211 | 0.010519 | 0.06358 | -2.91925 | Non-Significant | FALSE |
| Ago1 | -0.05231 | 7.316405 | -0.39399 | 0.701057 | 0.836156 | -6.18505 | Non-Significant | -0.06203 | 7.316405 | -0.47203 | 0.646051 | 0.779335 | -6.44823 | Non-Significant | FALSE |
| Agpat1 | 0.449145 | 7.315488 | 4.393339 | 0.001053 | 0.036195 | -0.52429 | Significant | 0.492469 | 7.315488 | 4.85788 | 0.000492 | 0.016463 | 0.09933 | Significant | FALSE |
| Agtpbp1 | -0.13041 | 9.735743 | -1.58101 | 0.141937 | 0.34121 | -5.18825 | Non-Significant | 0.02516 | 9.735743 | 0.302982 | 0.767508 | 0.863245 | -6.61449 | Non-Significant | FALSE |
| Ahnak | -0.27872 | 8.433696 | -1.94657 | 0.077339 | 0.246581 | -4.63236 | Non-Significant | -0.16439 | 8.433696 | -1.16297 | 0.269241 | 0.450079 | -5.96759 | Non-Significant | FALSE |
| AI593442 | 0.241748 | 8.000289 | 2.114035 | 0.057954 | 0.210274 | -4.34825 | Non-Significant | -0.1915 | 8.000289 | -1.67583 | 0.121686 | 0.274537 | -5.27206 | Non-Significant | FALSE |
| AI837181 | 0.189295 | 6.966667 | 1.027432 | 0.326092 | 0.550927 | -5.69388 | Non-Significant | -0.09162 | 6.966667 | -0.50248 | 0.625155 | 0.762702 | -6.38739 | Non-Significant | FALSE |
| Ajap1 | -0.13875 | 7.721659 | -1.40419 | 0.187625 | 0.395016 | -5.34849 | Non-Significant | -0.1263 | 7.721659 | -1.28949 | 0.223456 | 0.399494 | -5.77345 | Non-Significant | FALSE |
| Ak1 | 0.238438 | 8.042807 | 1.706469 | 0.115715 | 0.308551 | -4.95704 | Non-Significant | 0.084756 | 8.042807 | 0.611611 | 0.553119 | 0.714844 | -6.41773 | Non-Significant | FALSE |
| Ak5 | 0.40264 | 6.988768 | 1.874262 | 0.087446 | 0.26224 | -4.61779 | Non-Significant | 0.291127 | 6.988768 | 1.373493 | 0.19671 | 0.369937 | -5.58001 | Non-Significant | FALSE |
| Akap11 | -0.20139 | 10.39274 | -1.82997 | 0.094223 | 0.273709 | -4.82529 | Non-Significant | -0.35286 | 10.39274 | -3.1905 | 0.00851 | 0.057404 | -2.80702 | Non-Significant | FALSE |
| Akap6 | -0.14954 | 10.15737 | -1.52341 | 0.155634 | 0.359737 | -5.26158 | Non-Significant | -0.08792 | 10.15737 | -0.8885 | 0.393123 | 0.579892 | -6.24919 | Non-Significant | FALSE |
| Akirin1 | -0.11412 | 6.904568 | -0.95797 | 0.358486 | 0.579918 | -5.75388 | Non-Significant | -0.17522 | 6.904568 | -1.50696 | 0.159747 | 0.325148 | -5.42741 | Non-Significant | FALSE |
| Akr1a1 | 0.325316 | 8.105301 | 3.207975 | 0.008248 | 0.090526 | -2.52665 | Significant | 0.039274 | 8.105301 | 0.389355 | 0.704382 | 0.823351 | -6.53519 | Non-Significant | FALSE |
| Akt1 | 0.193183 | 8.406866 | 2.381537 | 0.036219 | 0.170339 | -3.93691 | Non-Significant | 0.021411 | 8.406866 | 0.265481 | 0.795506 | 0.879953 | -6.59951 | Non-Significant | FALSE |
| Akt2 | -0.13978 | 8.38497 | -1.52146 | 0.156116 | 0.360317 | -5.23012 | Non-Significant | 0.090036 | 8.38497 | 0.99423 | 0.341296 | 0.525282 | -6.13552 | Non-Significant | FALSE |
| Akt3 | -0.06874 | 8.785808 | -0.83135 | 0.423305 | 0.635372 | -6.01374 | Non-Significant | -0.19347 | 8.785808 | -2.34864 | 0.038392 | 0.13564 | -4.27064 | Non-Significant | FALSE |
| Aktip | -0.27369 | 8.127777 | -2.38204 | 0.036186 | 0.170339 | -3.92781 | Non-Significant | -0.37746 | 8.127777 | -3.31685 | 0.006792 | 0.050947 | -2.56559 | Non-Significant | FALSE |
| Alcam | 0.103035 | 8.789709 | 1.445446 | 0.175975 | 0.38217 | -5.35311 | Non-Significant | 0.167478 | 8.789709 | 2.358384 | 0.037736 | 0.134921 | -4.24984 | Non-Significant | FALSE |
| Aldoa | 0.386551 | 11.06763 | 4.81279 | 0.000529 | 0.033981 | 0.101923 | Significant | 0.516077 | 11.06763 | 6.344923 | 5.27E-05 | 0.00588 | 2.273221 | Significant | FALSE |
| Aldoc | 0.222242 | 10.47849 | 2.696964 | 0.020626 | 0.129661 | -3.43154 | Non-Significant | 0.248944 | 10.47849 | 2.994111 | 0.012099 | 0.070312 | -3.14768 | Non-Significant | FALSE |
| Alg2 | 0.145718 | 7.295545 | 1.084165 | 0.301284 | 0.52474 | -5.67561 | Non-Significant | -0.07429 | 7.295545 | -0.55584 | 0.589351 | 0.74063 | -6.39436 | Non-Significant | FALSE |
| Alkbh5 | 0.051557 | 8.033744 | 0.577303 | 0.57526 | 0.751853 | -6.14265 | Non-Significant | -0.00625 | 8.033744 | -0.07072 | 0.944879 | 0.973063 | -6.61407 | Non-Significant | FALSE |
| Ambra1 | -0.07284 | 7.499566 | -0.80228 | 0.439235 | 0.647013 | -5.95526 | Non-Significant | -0.02251 | 7.499566 | -0.25033 | 0.806908 | 0.888528 | -6.5503 | Non-Significant | FALSE |
| Amer3 | -0.18671 | 6.829671 | -1.96557 | 0.074869 | 0.243247 | -4.47647 | Non-Significant | -0.13037 | 6.829671 | -1.41737 | 0.183834 | 0.354179 | -5.53589 | Non-Significant | FALSE |
| Amfr | 0.164965 | 9.200276 | 2.34165 | 0.03887 | 0.175946 | -4.02944 | Non-Significant | 0.156249 | 9.200276 | 2.214957 | 0.04859 | 0.157599 | -4.49181 | Non-Significant | FALSE |
| Amotl1 | -0.1731 | 6.938512 | -1.17549 | 0.264403 | 0.487652 | -5.52883 | Non-Significant | 0.029639 | 6.938512 | 0.207811 | 0.839141 | 0.909438 | -6.50605 | Non-Significant | FALSE |
| Amph | 0.075801 | 9.227155 | 0.948805 | 0.362925 | 0.582595 | -5.91613 | Non-Significant | 0.154741 | 9.227155 | 1.933232 | 0.079118 | 0.209888 | -4.94028 | Non-Significant | FALSE |
| Amz2 | -0.04615 | 6.790072 | -0.47245 | 0.645762 | 0.799033 | -6.07255 | Non-Significant | 0.01124 | 6.790072 | 0.118709 | 0.907628 | 0.953935 | -6.49937 | Non-Significant | FALSE |
| Anapc1 | -0.10408 | 8.319681 | -0.89564 | 0.38946 | 0.602369 | -5.9268 | Non-Significant | -0.04164 | 8.319681 | -0.3628 | 0.723572 | 0.834148 | -6.56856 | Non-Significant | FALSE |
| Anapc2 | 0.114119 | 6.740937 | 0.950066 | 0.362312 | 0.582396 | -5.73557 | Non-Significant | -0.10208 | 6.740937 | -0.86192 | 0.406974 | 0.591232 | -6.10783 | Non-Significant | TRUE |
| Anapc5 | 0.083736 | 8.787222 | 1.113326 | 0.289103 | 0.512048 | -5.74328 | Non-Significant | 0.122097 | 8.787222 | 1.629573 | 0.131223 | 0.287004 | -5.38481 | Non-Significant | FALSE |
| Ank | 0.231047 | 8.269222 | 2.340004 | 0.038983 | 0.17605 | -3.99105 | Non-Significant | 0.297248 | 8.269222 | 3.044943 | 0.011044 | 0.065726 | -3.03863 | Non-Significant | FALSE |
| Ankib1 | -0.13777 | 8.055697 | -1.62034 | 0.133204 | 0.329687 | -5.07933 | Non-Significant | -0.11377 | 8.055697 | -1.35475 | 0.202436 | 0.376737 | -5.71284 | Non-Significant | FALSE |
| Ankle2 | 0.045664 | 6.958264 | 0.519859 | 0.613374 | 0.777386 | -6.06802 | Non-Significant | 0.136201 | 6.958264 | 1.588162 | 0.140313 | 0.29941 | -5.31043 | Non-Significant | FALSE |
| Ankmy2 | 0.229118 | 6.700235 | 2.453942 | 0.031847 | 0.159712 | -3.68532 | Non-Significant | 0.170613 | 6.700235 | 1.871038 | 0.087924 | 0.224675 | -4.86434 | Non-Significant | FALSE |
| Ankrd11 | -0.16008 | 8.444572 | -1.22205 | 0.247012 | 0.467732 | -5.60665 | Non-Significant | -0.06004 | 8.444572 | -0.46379 | 0.651763 | 0.783268 | -6.53428 | Non-Significant | FALSE |
| Ankrd12 | -0.2807 | 8.454841 | -2.01897 | 0.068313 | 0.232309 | -4.53034 | Non-Significant | -0.52476 | 8.454841 | -3.79501 | 0.002923 | 0.034091 | -1.74842 | Significant | FALSE |
| Ankrd13c | -0.02925 | 8.094876 | -0.28265 | 0.782648 | 0.882126 | -6.27748 | Non-Significant | -0.03836 | 8.094876 | -0.37501 | 0.714719 | 0.828713 | -6.54671 | Non-Significant | FALSE |
| Ankrd17 | -0.23786 | 8.952677 | -2.36873 | 0.03705 | 0.170529 | -3.98338 | Non-Significant | -0.18675 | 8.952677 | -1.86543 | 0.08876 | 0.226166 | -5.04621 | Non-Significant | FALSE |
| Ankrd33b | -0.0395 | 8.621006 | -0.31803 | 0.756368 | 0.870875 | -6.30411 | Non-Significant | -0.00766 | 8.621006 | -0.06214 | 0.951557 | 0.974467 | -6.65357 | Non-Significant | FALSE |
| Ankrd40 | 0.013877 | 7.879181 | 0.167765 | 0.869787 | 0.929978 | -6.29391 | Non-Significant | -0.00727 | 7.879181 | -0.08881 | 0.930815 | 0.966921 | -6.6052 | Non-Significant | FALSE |
| Ankrd50 | -0.35818 | 8.844383 | -2.91488 | 0.013948 | 0.111252 | -3.06065 | Non-Significant | -0.05283 | 8.844383 | -0.4328 | 0.673431 | 0.799559 | -6.56533 | Non-Significant | FALSE |
| Ankrd6 | 0.069665 | 7.637119 | 0.840631 | 0.418299 | 0.631762 | -5.92179 | Non-Significant | 0.460988 | 7.637119 | 5.674953 | 0.000139 | 0.009525 | 1.328154 | Significant | FALSE |
| Ankrd63 | 0.012024 | 6.886345 | 0.08304 | 0.935298 | 0.965916 | -6.18416 | Non-Significant | 0.320773 | 6.886345 | 2.290743 | 0.042527 | 0.144162 | -4.22885 | Non-Significant | FALSE |
| Anks1b | 0.068972 | 9.044406 | 0.805194 | 0.43762 | 0.647013 | -6.03907 | Non-Significant | 0.060318 | 9.044406 | 0.704552 | 0.49561 | 0.661966 | -6.40403 | Non-Significant | FALSE |
| Ano3 | -0.18016 | 8.856221 | -2.14782 | 0.054643 | 0.203463 | -4.33979 | Non-Significant | -0.3044 | 8.856221 | -3.64152 | 0.003824 | 0.038495 | -2.0243 | Significant | FALSE |
| Anxa6 | 0.141909 | 7.874994 | 1.689856 | 0.11892 | 0.311916 | -4.97061 | Non-Significant | 0.259554 | 7.874994 | 3.134112 | 0.009413 | 0.060523 | -2.86888 | Non-Significant | FALSE |
| Anxa7 | 0.400459 | 6.807315 | 4.158368 | 0.001564 | 0.041477 | -0.88133 | Significant | 0.312303 | 6.807315 | 3.303392 | 0.006957 | 0.051893 | -2.4763 | Non-Significant | FALSE |
| Ap1ar | -0.00079 | 7.975114 | -0.00591 | 0.995388 | 0.99725 | -6.31374 | Non-Significant | -0.07321 | 7.975114 | -0.55622 | 0.589099 | 0.74063 | -6.45061 | Non-Significant | FALSE |
| Ap1b1 | 0.208283 | 8.658992 | 2.181052 | 0.051563 | 0.19713 | -4.27395 | Non-Significant | 0.135655 | 8.658992 | 1.42691 | 0.181131 | 0.35201 | -5.65038 | Non-Significant | FALSE |
| Ap1s1 | 0.145528 | 7.371176 | 1.002119 | 0.337637 | 0.562308 | -5.76657 | Non-Significant | -0.05378 | 7.371176 | -0.37185 | 0.717009 | 0.830039 | -6.49207 | Non-Significant | FALSE |
| Ap2a1 | 0.230884 | 8.258809 | 2.066133 | 0.062974 | 0.221316 | -4.42868 | Non-Significant | 0.219231 | 8.258809 | 1.982089 | 0.072781 | 0.198479 | -4.82901 | Non-Significant | FALSE |
| Ap2a2 | 0.145018 | 9.438108 | 1.319662 | 0.213525 | 0.429616 | -5.52011 | Non-Significant | 0.308545 | 9.438108 | 2.793859 | 0.017334 | 0.086123 | -3.50773 | Non-Significant | FALSE |
| Ap2b1 | 0.088547 | 9.498202 | 0.853804 | 0.411265 | 0.625489 | -6.00239 | Non-Significant | 0.174391 | 9.498202 | 1.673835 | 0.122085 | 0.274973 | -5.32564 | Non-Significant | FALSE |
| Ap2m1 | 0.240125 | 9.713304 | 3.244324 | 0.00773 | 0.08771 | -2.50159 | Significant | 0.103161 | 9.713304 | 1.389107 | 0.192045 | 0.363974 | -5.71071 | Non-Significant | FALSE |
| Ap2s1 | 0.110525 | 7.563552 | 0.715972 | 0.488801 | 0.687144 | -6.03111 | Non-Significant | -0.16979 | 7.563552 | -1.10002 | 0.294612 | 0.474426 | -5.9717 | Non-Significant | FALSE |
| Ap3b2 | 0.106767 | 8.023528 | 1.396262 | 0.189938 | 0.396946 | -5.3677 | Non-Significant | 0.192387 | 8.023528 | 2.549966 | 0.026835 | 0.111216 | -3.88869 | Non-Significant | FALSE |
| Ap3m2 | 0.039382 | 8.717878 | 0.296774 | 0.77212 | 0.877275 | -6.31808 | Non-Significant | -0.09417 | 8.717878 | -0.71254 | 0.490844 | 0.65841 | -6.39373 | Non-Significant | FALSE |
| Apba2 | 0.245068 | 8.784453 | 2.864063 | 0.015281 | 0.11399 | -3.14118 | Non-Significant | 0.358332 | 8.784453 | 4.201049 | 0.001455 | 0.024052 | -1.05078 | Significant | TRUE |
| Apbb1 | 0.205936 | 10.29341 | 2.377542 | 0.036476 | 0.170477 | -3.96801 | Non-Significant | 0.209698 | 10.29341 | 2.401804 | 0.034939 | 0.128525 | -4.17304 | Non-Significant | FALSE |
| Apbb2 | 0.064964 | 7.202847 | 0.522779 | 0.611406 | 0.775849 | -6.10486 | Non-Significant | 0.045151 | 7.202847 | 0.367753 | 0.719975 | 0.832282 | -6.47617 | Non-Significant | FALSE |
| Apc | -0.5351 | 10.59942 | -2.8548 | 0.015537 | 0.114942 | -3.15996 | Non-Significant | -0.54705 | 10.59942 | -2.89644 | 0.014418 | 0.077428 | -3.31593 | Non-Significant | FALSE |
| Aplp1 | 0.426868 | 10.52777 | 5.245265 | 0.000267 | 0.027473 | 0.756456 | Significant | 0.289911 | 10.52777 | 3.541265 | 0.004562 | 0.042565 | -2.18701 | Non-Significant | FALSE |
| Apmap | 0.050625 | 7.196842 | 0.275225 | 0.788201 | 0.884291 | -6.21787 | Non-Significant | -0.23697 | 7.196842 | -1.29876 | 0.220365 | 0.396865 | -5.71754 | Non-Significant | FALSE |
| Apod | 0.233605 | 7.145703 | 1.885524 | 0.085796 | 0.25874 | -4.62242 | Non-Significant | 0.209711 | 7.145703 | 1.711884 | 0.114687 | 0.264131 | -5.1463 | Non-Significant | FALSE |
| Apoe | 0.162603 | 11.18167 | 1.061446 | 0.311042 | 0.533335 | -5.77776 | Non-Significant | 0.033606 | 11.18167 | 0.217858 | 0.831494 | 0.905105 | -6.60296 | Non-Significant | FALSE |
| Ar | -0.33883 | 6.906915 | -3.03224 | 0.011299 | 0.102571 | -2.76302 | Non-Significant | -0.39088 | 6.906915 | -3.59084 | 0.00418 | 0.040558 | -2.01732 | Significant | FALSE |
| Araf | -0.17514 | 8.058349 | -1.18117 | 0.262233 | 0.484651 | -5.63067 | Non-Significant | -0.30385 | 8.058349 | -2.06677 | 0.062904 | 0.182117 | -4.69276 | Non-Significant | FALSE |
| Arap2 | -0.35174 | 9.120374 | -3.8289 | 0.002756 | 0.052613 | -1.50479 | Significant | -0.27992 | 9.120374 | -3.05057 | 0.010933 | 0.065275 | -3.06358 | Non-Significant | FALSE |
| Arc | -0.54171 | 9.006055 | -2.51074 | 0.028781 | 0.150594 | -3.74657 | Non-Significant | -0.0567 | 9.006055 | -0.26401 | 0.796611 | 0.880811 | -6.62813 | Non-Significant | FALSE |
| Arcn1 | -0.06242 | 7.38448 | -0.78061 | 0.451362 | 0.659077 | -5.96411 | Non-Significant | -0.11174 | 7.38448 | -1.40882 | 0.186287 | 0.357178 | -5.59937 | Non-Significant | FALSE |
| Arel1 | -0.10252 | 10.04865 | -1.32689 | 0.211202 | 0.426063 | -5.51209 | Non-Significant | -0.0869 | 10.04865 | -1.11718 | 0.28752 | 0.467787 | -6.02668 | Non-Significant | FALSE |
| Arf1 | 0.280123 | 8.722428 | 3.034317 | 0.011257 | 0.102536 | -2.85032 | Non-Significant | 0.149286 | 8.722428 | 1.622144 | 0.132814 | 0.288933 | -5.38947 | Non-Significant | TRUE |
| Arf2 | -0.08098 | 6.839646 | -0.66485 | 0.519726 | 0.711204 | -5.97899 | Non-Significant | -0.18343 | 6.839646 | -1.53958 | 0.151676 | 0.314631 | -5.37281 | Non-Significant | FALSE |
| Arf3 | 0.144192 | 10.59997 | 2.126225 | 0.056738 | 0.208349 | -4.36719 | Non-Significant | 0.015746 | 10.59997 | 0.230801 | 0.821669 | 0.89851 | -6.61973 | Non-Significant | FALSE |
| Arf6 | 0.103036 | 7.272971 | 1.052071 | 0.315137 | 0.538225 | -5.70417 | Non-Significant | -0.00322 | 7.272971 | -0.03312 | 0.974167 | 0.989689 | -6.55483 | Non-Significant | FALSE |
| Arfgap2 | 0.014895 | 6.903709 | 0.152268 | 0.881709 | 0.937735 | -6.20053 | Non-Significant | -0.12513 | 6.903709 | -1.3032 | 0.218899 | 0.395554 | -5.67818 | Non-Significant | FALSE |
| Arfgef1 | -0.23658 | 8.816421 | -1.65243 | 0.126431 | 0.321796 | -5.0816 | Non-Significant | -0.31617 | 8.816421 | -2.21831 | 0.048305 | 0.157349 | -4.48791 | Non-Significant | FALSE |
| Arfgef2 | 0.002522 | 7.708104 | 0.022978 | 0.982076 | 0.989093 | -6.29975 | Non-Significant | 0.007625 | 7.708104 | 0.070079 | 0.945377 | 0.973063 | -6.5965 | Non-Significant | FALSE |
| Arfgef3 | -0.29901 | 8.887237 | -2.65618 | 0.02219 | 0.133016 | -3.50274 | Non-Significant | -0.2038 | 8.887237 | -1.81875 | 0.096014 | 0.238964 | -5.11636 | Non-Significant | FALSE |
| Arfip2 | 0.066309 | 7.427309 | 0.77412 | 0.455036 | 0.661916 | -5.96975 | Non-Significant | -0.02465 | 7.427309 | -0.28934 | 0.777653 | 0.868817 | -6.52756 | Non-Significant | FALSE |
| Arhgap10 | -0.08186 | 6.768364 | -0.77675 | 0.453545 | 0.66092 | -5.8859 | Non-Significant | -0.15405 | 6.768364 | -1.49837 | 0.161932 | 0.328527 | -5.41779 | Non-Significant | FALSE |
| Arhgap12 | -0.25381 | 7.003412 | -2.20345 | 0.04958 | 0.19584 | -4.13942 | Non-Significant | -0.3162 | 7.003412 | -2.8072 | 0.016924 | 0.084714 | -3.38774 | Non-Significant | FALSE |
| Arhgap20 | -0.32726 | 8.449989 | -2.77814 | 0.01783 | 0.124205 | -3.28186 | Non-Significant | -0.26828 | 8.449989 | -2.30507 | 0.041466 | 0.142367 | -4.33471 | Non-Significant | FALSE |
| Arhgap21 | -0.22449 | 8.86449 | -1.60386 | 0.136805 | 0.333968 | -5.14834 | Non-Significant | -0.11836 | 8.86449 | -0.84959 | 0.413508 | 0.595804 | -6.2902 | Non-Significant | FALSE |
| Arhgap23 | -0.17011 | 7.965728 | -1.53887 | 0.151848 | 0.355954 | -5.18369 | Non-Significant | -0.042 | 7.965728 | -0.38547 | 0.707173 | 0.824533 | -6.53782 | Non-Significant | FALSE |
| Arhgap32 | -0.15018 | 10.72039 | -1.41093 | 0.18568 | 0.393085 | -5.39339 | Non-Significant | -0.19211 | 10.72039 | -1.79093 | 0.100589 | 0.243817 | -5.13662 | Non-Significant | FALSE |
| Arhgap35 | -0.27878 | 10.34254 | -2.5331 | 0.027655 | 0.147531 | -3.71077 | Non-Significant | -0.15801 | 10.34254 | -1.42205 | 0.182504 | 0.353259 | -5.6587 | Non-Significant | FALSE |
| Arhgap39 | -0.16637 | 7.021611 | -1.38709 | 0.192642 | 0.400851 | -5.29264 | Non-Significant | 0.075803 | 7.021611 | 0.650793 | 0.528425 | 0.693181 | -6.31509 | Non-Significant | FALSE |
| Arhgap44 | 0.241229 | 8.705382 | 2.745862 | 0.018894 | 0.126343 | -3.33748 | Non-Significant | 0.414621 | 8.705382 | 4.742156 | 0.000593 | 0.017053 | -0.14859 | Significant | FALSE |
| Arhgap5 | -0.4083 | 8.752718 | -3.6621 | 0.003688 | 0.058824 | -1.78336 | Significant | -0.59967 | 8.752718 | -5.40398 | 0.000209 | 0.011476 | 0.887396 | Significant | FALSE |
| Arhgdia | 0.463436 | 7.916477 | 4.511357 | 0.000865 | 0.036195 | -0.35652 | Significant | 0.436235 | 7.916477 | 4.293043 | 0.001246 | 0.023246 | -0.86378 | Significant | FALSE |
| Arhgdig | 0.216672 | 7.1199 | 1.030025 | 0.324926 | 0.549685 | -5.71367 | Non-Significant | -0.17797 | 7.1199 | -0.85114 | 0.412683 | 0.595305 | -6.16563 | Non-Significant | FALSE |
| Arhgef10 | -0.18807 | 6.853114 | -1.5967 | 0.138395 | 0.336154 | -5.01258 | Non-Significant | -0.23845 | 6.853114 | -2.08031 | 0.061447 | 0.178476 | -4.58192 | Non-Significant | FALSE |
| Arhgef10l | -0.22813 | 7.84523 | -2.05405 | 0.064303 | 0.224067 | -4.43589 | Non-Significant | -0.34176 | 7.84523 | -3.09562 | 0.010085 | 0.062374 | -2.93497 | Non-Significant | FALSE |
| Arhgef11 | -0.0293 | 7.981001 | -0.2827 | 0.782612 | 0.882126 | -6.26722 | Non-Significant | 0.239372 | 7.981001 | 2.349112 | 0.03836 | 0.13564 | -4.22762 | Non-Significant | FALSE |
| Arhgef12 | -0.12218 | 9.845778 | -1.04835 | 0.316772 | 0.540328 | -5.82386 | Non-Significant | -0.00458 | 9.845778 | -0.03899 | 0.969593 | 0.985416 | -6.66191 | Non-Significant | FALSE |
| Arhgef17 | -0.03732 | 8.732676 | -0.38906 | 0.704593 | 0.839368 | -6.28371 | Non-Significant | 0.025066 | 8.732676 | 0.262736 | 0.797568 | 0.881505 | -6.62314 | Non-Significant | FALSE |
| Arhgef2 | 0.077657 | 8.081063 | 0.712567 | 0.490825 | 0.689157 | -6.05332 | Non-Significant | 0.177332 | 8.081063 | 1.649625 | 0.127011 | 0.281568 | -5.31848 | Non-Significant | FALSE |
| Arhgef25 | 0.300345 | 8.034777 | 2.969998 | 0.012634 | 0.107069 | -2.92838 | Non-Significant | 0.226781 | 8.034777 | 2.264893 | 0.044509 | 0.148925 | -4.36491 | Non-Significant | FALSE |
| Arhgef28 | -0.02088 | 7.034645 | -0.192 | 0.851209 | 0.918798 | -6.19747 | Non-Significant | 0.187048 | 7.034645 | 1.761536 | 0.105638 | 0.251817 | -5.0732 | Non-Significant | FALSE |
| Arhgef3 | 0.010911 | 7.993185 | 0.124063 | 0.903484 | 0.94958 | -6.30589 | Non-Significant | -0.03663 | 7.993185 | -0.42087 | 0.681888 | 0.806936 | -6.52095 | Non-Significant | FALSE |
| Arhgef4 | -0.05579 | 9.704161 | -0.74128 | 0.473916 | 0.675449 | -6.09418 | Non-Significant | -0.04034 | 9.704161 | -0.53341 | 0.604271 | 0.747454 | -6.51418 | Non-Significant | FALSE |
| Arhgef9 | 0.162061 | 9.489003 | 2.305776 | 0.041414 | 0.179461 | -4.08969 | Non-Significant | 0.082734 | 9.489003 | 1.173631 | 0.265117 | 0.444851 | -5.96832 | Non-Significant | FALSE |
| Arid1a | 0.005483 | 7.772385 | 0.032972 | 0.974282 | 0.985692 | -6.30227 | Non-Significant | 0.045421 | 7.772385 | 0.276069 | 0.787569 | 0.874786 | -6.56307 | Non-Significant | FALSE |
| Arid1b | -0.20722 | 7.793188 | -1.63816 | 0.129403 | 0.324781 | -5.04227 | Non-Significant | -0.08322 | 7.793188 | -0.66633 | 0.518811 | 0.683748 | -6.37466 | Non-Significant | TRUE |
| Arid2 | -0.31821 | 7.125244 | -2.16205 | 0.053304 | 0.200769 | -4.21403 | Non-Significant | -0.22033 | 7.125244 | -1.53337 | 0.153185 | 0.315637 | -5.42325 | Non-Significant | FALSE |
| Arih2 | -0.10206 | 7.146534 | -1.10991 | 0.290512 | 0.513674 | -5.63653 | Non-Significant | -0.14859 | 7.146534 | -1.64195 | 0.128609 | 0.283468 | -5.26768 | Non-Significant | FALSE |
| Arl1 | -0.1989 | 7.368163 | -1.36605 | 0.198967 | 0.407677 | -5.38317 | Non-Significant | -0.56525 | 7.368163 | -3.90278 | 0.002425 | 0.030641 | -1.51205 | Significant | FALSE |
| Arl2bp | -0.02664 | 7.685987 | -0.26167 | 0.798366 | 0.891587 | -6.26467 | Non-Significant | -0.08866 | 7.685987 | -0.87655 | 0.399309 | 0.583706 | -6.20029 | Non-Significant | FALSE |
| Arl4a | -0.37215 | 6.800948 | -3.82597 | 0.00277 | 0.052613 | -1.43438 | Significant | -0.66797 | 6.800948 | -6.93816 | 2.35E-05 | 0.003936 | 3.076153 | Significant | FALSE |
| Arl4c | 0.158597 | 7.070806 | 1.711187 | 0.114819 | 0.308101 | -4.87073 | Non-Significant | 0.093931 | 7.070806 | 1.028258 | 0.32572 | 0.507433 | -5.9913 | Non-Significant | FALSE |
| Arl6ip1 | -0.01495 | 9.051758 | -0.10238 | 0.920281 | 0.955922 | -6.36789 | Non-Significant | -0.1826 | 9.051758 | -1.25178 | 0.236393 | 0.413765 | -5.88069 | Non-Significant | FALSE |
| Arl6ip5 | 0.077797 | 6.873725 | 0.620367 | 0.547545 | 0.733162 | -6.0084 | Non-Significant | -0.07296 | 6.873725 | -0.59282 | 0.565187 | 0.722583 | -6.32908 | Non-Significant | FALSE |
| Arl8a | 0.568026 | 7.517051 | 5.734293 | 0.000127 | 0.023095 | 1.460046 | Significant | 0.390415 | 7.517051 | 3.953745 | 0.002221 | 0.02915 | -1.40691 | Significant | FALSE |
| Armcx1 | -0.13085 | 7.176631 | -1.05857 | 0.312292 | 0.534388 | -5.69891 | Non-Significant | -0.2773 | 7.176631 | -2.27436 | 0.043773 | 0.147267 | -4.30271 | Non-Significant | FALSE |
| Armcx2 | 0.232537 | 7.772107 | 2.619435 | 0.0237 | 0.136196 | -3.50833 | Non-Significant | 0.344598 | 7.772107 | 3.932922 | 0.002302 | 0.030069 | -1.47125 | Significant | FALSE |
| Armcx3 | -0.04468 | 7.56882 | -0.49057 | 0.633286 | 0.79146 | -6.16894 | Non-Significant | -0.15132 | 7.56882 | -1.66935 | 0.122985 | 0.275397 | -5.26228 | Non-Significant | FALSE |
| Arnt2 | 0.032877 | 9.1571 | 0.378779 | 0.711999 | 0.842338 | -6.29841 | Non-Significant | -0.01004 | 9.1571 | -0.11559 | 0.910042 | 0.955347 | -6.65778 | Non-Significant | FALSE |
| Arpc1a | 0.0945 | 8.224672 | 1.12539 | 0.284175 | 0.506445 | -5.69248 | Non-Significant | 0.110202 | 8.224672 | 1.327574 | 0.210982 | 0.386993 | -5.7515 | Non-Significant | FALSE |
| Arpc2 | 0.046936 | 8.008675 | 0.43454 | 0.672225 | 0.815686 | -6.21666 | Non-Significant | -0.06394 | 8.008675 | -0.59742 | 0.562217 | 0.721392 | -6.42701 | Non-Significant | FALSE |
| Arpc3 | -0.08725 | 7.61455 | -0.49746 | 0.628573 | 0.788809 | -6.16969 | Non-Significant | -0.20654 | 7.61455 | -1.18267 | 0.261658 | 0.441538 | -5.88916 | Non-Significant | FALSE |
| Arpc4 | 0.330414 | 7.790295 | 2.516047 | 0.02851 | 0.150001 | -3.68264 | Non-Significant | 0.227283 | 7.790295 | 1.744827 | 0.108608 | 0.25663 | -5.16192 | Non-Significant | FALSE |
| Arpc5l | -0.37801 | 7.153479 | -2.67529 | 0.021443 | 0.131219 | -3.38777 | Non-Significant | -0.4005 | 7.153479 | -2.89477 | 0.014461 | 0.077453 | -3.2553 | Non-Significant | FALSE |
| Arpp21 | -0.07664 | 9.681593 | -0.91606 | 0.379111 | 0.596509 | -5.95023 | Non-Significant | -0.06792 | 9.681593 | -0.80816 | 0.435981 | 0.615475 | -6.32506 | Non-Significant | FALSE |
| Arrb1 | -0.09582 | 9.771978 | -0.89063 | 0.392028 | 0.603727 | -5.97293 | Non-Significant | 0.125197 | 9.771978 | 1.154366 | 0.272604 | 0.453156 | -5.98766 | Non-Significant | FALSE |
| Arvcf | -0.18843 | 6.989042 | -1.43677 | 0.178372 | 0.384606 | -5.22621 | Non-Significant | 0.070637 | 6.989042 | 0.55538 | 0.589655 | 0.740664 | -6.37081 | Non-Significant | FALSE |
| Arxes2 | 0.347174 | 6.808834 | 2.652121 | 0.022352 | 0.13302 | -3.36647 | Non-Significant | 0.349265 | 6.808834 | 2.725497 | 0.019597 | 0.092566 | -3.47675 | Non-Significant | FALSE |
| Asap1 | -0.02282 | 7.932103 | -0.12937 | 0.899381 | 0.946941 | -6.30094 | Non-Significant | 0.057966 | 7.932103 | 0.332931 | 0.745392 | 0.849429 | -6.55426 | Non-Significant | FALSE |
| Ash1l | -0.33986 | 8.877302 | -2.71042 | 0.020134 | 0.128379 | -3.41175 | Non-Significant | -0.40214 | 8.877302 | -3.22068 | 0.008063 | 0.055466 | -2.76364 | Non-Significant | FALSE |
| Asic2 | -0.05821 | 7.89259 | -0.68078 | 0.509969 | 0.704267 | -6.06896 | Non-Significant | 0.037323 | 7.89259 | 0.442298 | 0.66677 | 0.794249 | -6.50792 | Non-Significant | FALSE |
| Asna1 | 0.113686 | 7.100771 | 0.808011 | 0.436063 | 0.646393 | -5.90283 | Non-Significant | 0.004214 | 7.100771 | 0.030357 | 0.976321 | 0.989803 | -6.53727 | Non-Significant | FALSE |
| Asns | 0.138299 | 7.829344 | 1.304127 | 0.218592 | 0.433147 | -5.46997 | Non-Significant | 0.290959 | 7.829344 | 2.78359 | 0.017657 | 0.086928 | -3.47852 | Non-Significant | FALSE |
| Asphd2 | 0.217927 | 8.406221 | 2.594377 | 0.024787 | 0.139371 | -3.58364 | Non-Significant | 0.031727 | 8.406221 | 0.379779 | 0.711277 | 0.826736 | -6.55992 | Non-Significant | FALSE |
| Asrgl1 | -0.08997 | 8.500099 | -1.03515 | 0.322629 | 0.54831 | -5.81382 | Non-Significant | -0.20998 | 8.500099 | -2.43229 | 0.033097 | 0.124929 | -4.11983 | Non-Significant | FALSE |
| Astn1 | 0.18027 | 8.797084 | 1.501958 | 0.161017 | 0.364501 | -5.28101 | Non-Significant | 0.086871 | 8.797084 | 0.725836 | 0.482967 | 0.65175 | -6.38293 | Non-Significant | FALSE |
| Asxl1 | -0.17433 | 6.976908 | -1.10318 | 0.293299 | 0.517087 | -5.62213 | Non-Significant | -0.21558 | 6.976908 | -1.39595 | 0.190029 | 0.361433 | -5.58116 | Non-Significant | FALSE |
| Atcay | 0.117566 | 8.865057 | 1.624826 | 0.132238 | 0.329426 | -5.11637 | Non-Significant | 0.256811 | 8.865057 | 3.558411 | 0.004426 | 0.041796 | -2.16496 | Non-Significant | FALSE |
| Ate1 | -0.03247 | 7.566421 | -0.38058 | 0.7107 | 0.842149 | -6.21869 | Non-Significant | -0.16872 | 7.566421 | -1.98488 | 0.072433 | 0.198138 | -4.79064 | Non-Significant | FALSE |
| Atf2 | 0.014588 | 7.529071 | 0.112839 | 0.912173 | 0.953474 | -6.28426 | Non-Significant | -0.15155 | 7.529071 | -1.17615 | 0.264148 | 0.443709 | -5.88803 | Non-Significant | FALSE |
| Atf4 | 0.124617 | 7.026635 | 0.781806 | 0.450687 | 0.658451 | -5.8974 | Non-Significant | 0.331902 | 7.026635 | 2.129047 | 0.05646 | 0.17098 | -4.49994 | Non-Significant | FALSE |
| Atf6 | -0.26285 | 7.889491 | -2.3169 | 0.040608 | 0.178092 | -4.0167 | Non-Significant | -0.1023 | 7.889491 | -0.91496 | 0.379663 | 0.565169 | -6.18201 | Non-Significant | FALSE |
| Atg13 | 0.232947 | 7.005697 | 2.60371 | 0.024376 | 0.138303 | -3.47544 | Non-Significant | 0.10569 | 7.005697 | 1.198537 | 0.255677 | 0.436117 | -5.80044 | Non-Significant | FALSE |
| Atg4b | -0.06965 | 7.19828 | -0.8431 | 0.416975 | 0.630754 | -5.88681 | Non-Significant | 0.011262 | 7.19828 | 0.138444 | 0.89237 | 0.947977 | -6.54269 | Non-Significant | FALSE |
| Atg9a | 0.204953 | 6.694488 | 1.937843 | 0.078499 | 0.247317 | -4.48487 | Non-Significant | 0.299605 | 6.694488 | 2.923477 | 0.013735 | 0.075681 | -3.13032 | Non-Significant | FALSE |
| Atmin | -0.16189 | 8.379606 | -1.51433 | 0.157892 | 0.362327 | -5.24504 | Non-Significant | -0.15948 | 8.379606 | -1.50837 | 0.15939 | 0.325093 | -5.53784 | Non-Significant | FALSE |
| Atp13a2 | 0.077737 | 8.735585 | 0.80422 | 0.438159 | 0.647013 | -6.0287 | Non-Significant | 0.138243 | 8.735585 | 1.437073 | 0.178289 | 0.350222 | -5.64293 | Non-Significant | FALSE |
| Atp1b2 | 0.542033 | 7.084665 | 3.118034 | 0.009688 | 0.09469 | -2.61068 | Significant | 0.443002 | 7.084665 | 2.571962 | 0.0258 | 0.108633 | -3.75837 | Non-Significant | FALSE |
| Atp1b3 | 0.060946 | 7.091651 | 0.587828 | 0.568417 | 0.746919 | -6.04754 | Non-Significant | 0.156558 | 7.091651 | 1.536957 | 0.152313 | 0.314975 | -5.3917 | Non-Significant | FALSE |
| Atp2b2 | -0.38459 | 11.76351 | -2.8862 | 0.014685 | 0.112151 | -3.07826 | Non-Significant | -0.12268 | 11.76351 | -0.90639 | 0.38399 | 0.570342 | -6.17565 | Non-Significant | TRUE |
| Atp2b4 | 0.261953 | 9.695109 | 2.177153 | 0.051915 | 0.197385 | -4.29671 | Non-Significant | 0.342125 | 9.695109 | 2.825562 | 0.016375 | 0.08275 | -3.45143 | Non-Significant | FALSE |
| Atp6ap1 | 0.237113 | 8.644672 | 2.733469 | 0.019318 | 0.126801 | -3.35857 | Non-Significant | 0.231914 | 8.644672 | 2.68693 | 0.021 | 0.096532 | -3.68182 | Non-Significant | FALSE |
| Atp6v0a1 | 0.175509 | 8.941405 | 1.699152 | 0.117117 | 0.309899 | -5.01556 | Non-Significant | 0.179159 | 8.941405 | 1.736769 | 0.110068 | 0.258336 | -5.23379 | Non-Significant | FALSE |
| Atp6v0a2 | -0.06256 | 7.296834 | -0.56739 | 0.581747 | 0.757069 | -6.0903 | Non-Significant | 0.116493 | 7.296834 | 1.072547 | 0.306245 | 0.486143 | -5.97711 | Non-Significant | FALSE |
| Atp6v0b | 0.018288 | 8.893038 | 0.119347 | 0.907134 | 0.950361 | -6.36322 | Non-Significant | -0.1185 | 8.893038 | -0.77514 | 0.454456 | 0.629934 | -6.35077 | Non-Significant | FALSE |
| Atp6v0d1 | 0.143728 | 8.248207 | 1.681958 | 0.120471 | 0.314445 | -5.00019 | Non-Significant | 0.135164 | 8.248207 | 1.598818 | 0.137923 | 0.295722 | -5.39685 | Non-Significant | FALSE |
| Atp6v0e2 | 0.333289 | 8.366164 | 3.285389 | 0.007183 | 0.084005 | -2.40382 | Significant | 0.137469 | 8.366164 | 1.362303 | 0.200112 | 0.373971 | -5.71224 | Non-Significant | FALSE |
| Atp6v1b2 | 0.189841 | 9.899514 | 2.186436 | 0.051079 | 0.196256 | -4.28224 | Non-Significant | 0.202002 | 9.899514 | 2.311592 | 0.040991 | 0.141644 | -4.32966 | Non-Significant | FALSE |
| Atp6v1c1 | -0.03163 | 9.519093 | -0.25384 | 0.804258 | 0.895263 | -6.34353 | Non-Significant | -0.03885 | 9.519093 | -0.31072 | 0.761775 | 0.86027 | -6.61405 | Non-Significant | FALSE |
| Atp6v1d | -0.03102 | 8.526943 | -0.26211 | 0.798037 | 0.891587 | -6.3174 | Non-Significant | -0.11504 | 8.526943 | -0.97857 | 0.348646 | 0.533834 | -6.15933 | Non-Significant | FALSE |
| Atp6v1e1 | 0.032243 | 9.579698 | 0.223833 | 0.826955 | 0.909149 | -6.35081 | Non-Significant | -0.03025 | 9.579698 | -0.20926 | 0.838036 | 0.908976 | -6.64203 | Non-Significant | FALSE |
| Atp6v1f | -0.02513 | 7.040813 | -0.14316 | 0.888732 | 0.941093 | -6.22582 | Non-Significant | -0.21242 | 7.040813 | -1.22807 | 0.244831 | 0.424649 | -5.78627 | Non-Significant | FALSE |
| Atp6v1g2 | 0.358368 | 7.042317 | 2.573104 | 0.025748 | 0.141478 | -3.52397 | Non-Significant | 0.246247 | 7.042317 | 1.789183 | 0.100883 | 0.244051 | -5.01907 | Non-Significant | FALSE |
| Atp6v1h | -0.0002 | 9.174555 | -0.00146 | 0.99886 | 0.999164 | -6.37428 | Non-Significant | -0.05389 | 9.174555 | -0.39507 | 0.700279 | 0.82108 | -6.58223 | Non-Significant | FALSE |
| Atp9a | 0.111074 | 10.53229 | 1.355479 | 0.20221 | 0.412673 | -5.46822 | Non-Significant | 0.141775 | 10.53229 | 1.714415 | 0.114209 | 0.263438 | -5.25348 | Non-Significant | FALSE |
| Atraid | -0.04849 | 6.64919 | -0.29061 | 0.776709 | 0.880248 | -6.13981 | Non-Significant | -0.35387 | 6.64919 | -2.1389 | 0.055499 | 0.16933 | -4.44979 | Non-Significant | FALSE |
| Atrnl1 | 0.016844 | 9.04368 | 0.23571 | 0.817952 | 0.903436 | -6.34327 | Non-Significant | 0.00205 | 9.04368 | 0.028709 | 0.977607 | 0.989803 | -6.66442 | Non-Significant | FALSE |
| Atxn1 | -0.2852 | 8.891513 | -3.23029 | 0.007926 | 0.089181 | -2.52193 | Significant | -0.19416 | 8.891513 | -2.20894 | 0.049106 | 0.158439 | -4.50359 | Non-Significant | TRUE |
| Atxn10 | 0.201383 | 8.330143 | 2.509451 | 0.028848 | 0.150594 | -3.71918 | Non-Significant | 0.107474 | 8.330143 | 1.349888 | 0.203943 | 0.378227 | -5.72839 | Non-Significant | FALSE |
| Atxn7l3b | 0.134078 | 9.171286 | 1.696689 | 0.117592 | 0.310257 | -5.02325 | Non-Significant | 0.005617 | 9.171286 | 0.071057 | 0.944617 | 0.973063 | -6.66215 | Non-Significant | FALSE |
| AU040320 | 0.159683 | 6.904426 | 1.253068 | 0.235941 | 0.455551 | -5.42724 | Non-Significant | 0.28478 | 6.904426 | 2.2909 | 0.042515 | 0.144162 | -4.22562 | Non-Significant | FALSE |
| Auts2 | -0.40598 | 6.965354 | -2.49178 | 0.029771 | 0.153779 | -3.66484 | Non-Significant | -0.25649 | 6.965354 | -1.6322 | 0.130665 | 0.286265 | -5.27603 | Non-Significant | FALSE |
| Avl9 | -0.24323 | 7.915135 | -3.13994 | 0.009316 | 0.093176 | -2.63623 | Significant | -0.07954 | 7.915135 | -1.04209 | 0.31954 | 0.500133 | -6.06082 | Non-Significant | FALSE |
| Azi2 | -0.029 | 7.382037 | -0.30486 | 0.766118 | 0.874537 | -6.21977 | Non-Significant | 0.082618 | 7.382037 | 0.878831 | 0.398123 | 0.583436 | -6.17098 | Non-Significant | FALSE |
| Azin2 | 0.093057 | 7.217152 | 0.57962 | 0.57375 | 0.750979 | -6.07792 | Non-Significant | -0.02893 | 7.217152 | -0.18191 | 0.858935 | 0.924842 | -6.53301 | Non-Significant | FALSE |
| B3galt2 | -0.02908 | 7.22926 | -0.32597 | 0.750514 | 0.868848 | -6.20442 | Non-Significant | -0.16878 | 7.22926 | -1.91215 | 0.082008 | 0.213967 | -4.87672 | Non-Significant | FALSE |
| B3gat1 | 0.301552 | 8.15674 | 2.796957 | 0.017238 | 0.123101 | -3.22469 | Non-Significant | 0.355901 | 8.15674 | 3.341192 | 0.006504 | 0.050503 | -2.51428 | Non-Significant | FALSE |
| B3gat3 | 0.371862 | 8.109096 | 2.865089 | 0.015253 | 0.11399 | -3.1094 | Non-Significant | 0.230534 | 8.109096 | 1.791013 | 0.100575 | 0.243817 | -5.11116 | Non-Significant | FALSE |
| B4galnt1 | 0.241808 | 7.043694 | 2.034063 | 0.06656 | 0.229024 | -4.37823 | Non-Significant | 0.464808 | 7.043694 | 3.986433 | 0.002099 | 0.028247 | -1.32178 | Significant | FALSE |
| B4galnt4 | 0.198868 | 7.539525 | 2.069907 | 0.062564 | 0.220745 | -4.3882 | Non-Significant | 0.069827 | 7.539525 | 0.729656 | 0.480719 | 0.650513 | -6.29989 | Non-Significant | FALSE |
| B4galt2 | 0.208588 | 7.840608 | 2.066692 | 0.062913 | 0.221316 | -4.41463 | Non-Significant | -0.08156 | 7.840608 | -0.81028 | 0.434809 | 0.614469 | -6.26108 | Non-Significant | FALSE |
| B4gat1 | 0.114349 | 7.896157 | 1.315849 | 0.21476 | 0.430324 | -5.4612 | Non-Significant | 0.146498 | 7.896157 | 1.706326 | 0.115742 | 0.265123 | -5.22885 | Non-Significant | FALSE |
| Bace1 | 0.080146 | 7.922766 | 0.776633 | 0.453611 | 0.66092 | -5.99995 | Non-Significant | 0.095656 | 7.922766 | 0.937932 | 0.368245 | 0.5534 | -6.15864 | Non-Significant | FALSE |
| Bag1 | -0.19571 | 7.756091 | -1.02275 | 0.328204 | 0.55244 | -5.78339 | Non-Significant | -0.52204 | 7.756091 | -2.72545 | 0.019598 | 0.092566 | -3.57295 | Non-Significant | FALSE |
| Bag4 | 0.210814 | 7.118966 | 2.410179 | 0.034424 | 0.165804 | -3.80634 | Non-Significant | 0.040298 | 7.118966 | 0.465651 | 0.650471 | 0.782552 | -6.41982 | Non-Significant | FALSE |
| Bag5 | -0.2742 | 7.187867 | -2.41853 | 0.033917 | 0.165358 | -3.81008 | Non-Significant | -0.2138 | 7.187867 | -1.92244 | 0.080586 | 0.21233 | -4.86603 | Non-Significant | FALSE |
| Bag6 | 0.152455 | 8.437707 | 1.558947 | 0.147054 | 0.349433 | -5.17633 | Non-Significant | 0.382863 | 8.437707 | 3.958354 | 0.002203 | 0.029062 | -1.45242 | Significant | FALSE |
| Bap1 | -0.07731 | 9.429698 | -1.18142 | 0.262137 | 0.484651 | -5.68159 | Non-Significant | 0.058271 | 9.429698 | 0.88709 | 0.39385 | 0.579892 | -6.25679 | Non-Significant | FALSE |
| Basp1 | 0.148141 | 10.05408 | 1.682993 | 0.120266 | 0.314218 | -5.04306 | Non-Significant | 0.116746 | 10.05408 | 1.317904 | 0.214094 | 0.389499 | -5.79514 | Non-Significant | FALSE |
| Bbs1 | -0.23606 | 7.19194 | -2.33629 | 0.03924 | 0.176064 | -3.93596 | Non-Significant | 0.014161 | 7.19194 | 0.143355 | 0.88858 | 0.945418 | -6.548 | Non-Significant | FALSE |
| BC005537 | 0.213632 | 8.372228 | 1.444966 | 0.176107 | 0.38217 | -5.33132 | Non-Significant | -0.03495 | 8.372228 | -0.23752 | 0.816585 | 0.895501 | -6.60501 | Non-Significant | FALSE |
| BC031181 | -0.09186 | 7.237989 | -0.61483 | 0.551064 | 0.73513 | -6.07287 | Non-Significant | -0.32256 | 7.237989 | -2.18047 | 0.051615 | 0.162006 | -4.461 | Non-Significant | FALSE |
| Bcan | 0.166205 | 8.639166 | 1.752374 | 0.107257 | 0.293271 | -4.92626 | Non-Significant | 0.139451 | 8.639166 | 1.478076 | 0.1672 | 0.335904 | -5.58337 | Non-Significant | FALSE |
| Bcas1 | -0.37669 | 7.37177 | -1.99446 | 0.071252 | 0.237473 | -4.51116 | Non-Significant | -0.7633 | 7.37177 | -4.06786 | 0.001825 | 0.026423 | -1.23277 | Significant | FALSE |
| Bcas3 | 0.168969 | 6.928154 | 1.785495 | 0.101506 | 0.285376 | -4.74663 | Non-Significant | 0.097568 | 6.928154 | 1.050721 | 0.31573 | 0.49755 | -5.95175 | Non-Significant | FALSE |
| Bcl11b | -0.03544 | 8.478851 | -0.31903 | 0.755631 | 0.870875 | -6.28953 | Non-Significant | 0.163102 | 8.478851 | 1.485071 | 0.165368 | 0.333728 | -5.56887 | Non-Significant | FALSE |
| Bcr | -0.01323 | 8.503939 | -0.0898 | 0.93005 | 0.962924 | -6.33744 | Non-Significant | 0.287299 | 8.503939 | 1.973873 | 0.073812 | 0.199434 | -4.85942 | Non-Significant | FALSE |
| Becn1 | 0.067911 | 8.241622 | 0.704179 | 0.495833 | 0.692139 | -6.07149 | Non-Significant | 0.038653 | 8.241622 | 0.405088 | 0.693112 | 0.815892 | -6.54149 | Non-Significant | FALSE |
| Begain | 0.049879 | 7.070869 | 0.472372 | 0.645815 | 0.799033 | -6.11565 | Non-Significant | -0.00856 | 7.070869 | -0.08244 | 0.935762 | 0.969053 | -6.53324 | Non-Significant | FALSE |
| Bend6 | 0.141514 | 7.249722 | 1.413395 | 0.184972 | 0.392204 | -5.29443 | Non-Significant | -0.08015 | 7.249722 | -0.80615 | 0.437089 | 0.615742 | -6.21593 | Non-Significant | FALSE |
| Bex2 | -0.11935 | 7.749918 | -0.92292 | 0.375681 | 0.594958 | -5.87486 | Non-Significant | -0.32882 | 7.749918 | -2.54924 | 0.026869 | 0.111216 | -3.87581 | Non-Significant | FALSE |
| Bex3 | 0.111044 | 7.12841 | 0.518808 | 0.614084 | 0.777917 | -6.11112 | Non-Significant | -0.29971 | 7.12841 | -1.40823 | 0.186458 | 0.357178 | -5.57324 | Non-Significant | FALSE |
| Bfar | -0.14709 | 6.80619 | -1.19443 | 0.257214 | 0.480014 | -5.49407 | Non-Significant | -0.08633 | 6.80619 | -0.72397 | 0.484066 | 0.65175 | -6.24104 | Non-Significant | FALSE |
| Bhlhe22 | 0.338891 | 7.139638 | 2.734908 | 0.019269 | 0.126801 | -3.26414 | Non-Significant | 0.262647 | 7.139638 | 2.139783 | 0.055414 | 0.16933 | -4.48794 | Non-Significant | FALSE |
| Bhlhe40 | 0.29325 | 7.304749 | 1.506821 | 0.159783 | 0.364501 | -5.16643 | Non-Significant | 0.260902 | 7.304749 | 1.350665 | 0.203702 | 0.378041 | -5.64353 | Non-Significant | FALSE |
| Bin1 | 0.15765 | 8.398769 | 1.897427 | 0.084083 | 0.255299 | -4.69539 | Non-Significant | 0.222421 | 8.398769 | 2.702909 | 0.020407 | 0.09519 | -3.6426 | Non-Significant | FALSE |
| Birc2 | -0.48585 | 7.119312 | -4.35843 | 0.001116 | 0.036452 | -0.58346 | Significant | -0.37275 | 7.119312 | -3.4411 | 0.005446 | 0.046269 | -2.30127 | Non-Significant | FALSE |
| Blcap | 0.336392 | 6.954227 | 4.019879 | 0.001982 | 0.044463 | -1.11105 | Significant | 0.24642 | 6.954227 | 2.990748 | 0.012172 | 0.070404 | -3.03676 | Non-Significant | FALSE |
| Blmh | -0.02647 | 7.525548 | -0.18652 | 0.855404 | 0.920879 | -6.27124 | Non-Significant | -0.12042 | 7.525548 | -0.85295 | 0.41172 | 0.594387 | -6.20806 | Non-Significant | FALSE |
| Bmpr1a | 0.012097 | 7.277107 | 0.102945 | 0.919844 | 0.955922 | -6.25387 | Non-Significant | -0.01994 | 7.277107 | -0.17152 | 0.866904 | 0.930116 | -6.5439 | Non-Significant | FALSE |
| Bmpr2 | -0.09322 | 9.259447 | -0.70825 | 0.493399 | 0.690708 | -6.11618 | Non-Significant | -0.19888 | 9.259447 | -1.5102 | 0.15893 | 0.324649 | -5.55589 | Non-Significant | FALSE |
| Bod1 | 0.12472 | 7.774294 | 0.988696 | 0.343879 | 0.564628 | -5.81138 | Non-Significant | -0.02436 | 7.774294 | -0.19416 | 0.849556 | 0.918124 | -6.58044 | Non-Significant | FALSE |
| Bpgm | -0.4132 | 7.048015 | -2.9061 | 0.01417 | 0.111252 | -2.99091 | Non-Significant | -0.4361 | 7.048015 | -3.14553 | 0.009223 | 0.059948 | -2.81056 | Non-Significant | FALSE |
| Brap | -0.2036 | 8.139345 | -2.2424 | 0.046305 | 0.188638 | -4.14946 | Non-Significant | -0.10153 | 8.139345 | -1.134 | 0.280696 | 0.461452 | -5.97758 | Non-Significant | FALSE |
| Brd1 | -0.10065 | 7.62325 | -1.20778 | 0.252242 | 0.474235 | -5.57319 | Non-Significant | 0.021942 | 7.62325 | 0.266461 | 0.79477 | 0.879953 | -6.55784 | Non-Significant | FALSE |
| Bri3bp | 0.172848 | 8.369675 | 1.904205 | 0.083121 | 0.254108 | -4.68307 | Non-Significant | 0.227032 | 8.369675 | 2.526172 | 0.028 | 0.113993 | -3.9439 | Non-Significant | FALSE |
| Brinp1 | 0.037306 | 9.4177 | 0.371909 | 0.716965 | 0.844713 | -6.3034 | Non-Significant | -0.02382 | 9.4177 | -0.23686 | 0.817082 | 0.89568 | -6.63525 | Non-Significant | FALSE |
| Brinp2 | -0.05065 | 9.155273 | -0.53736 | 0.601631 | 0.771263 | -6.22307 | Non-Significant | -0.01906 | 9.155273 | -0.20219 | 0.843426 | 0.913705 | -6.64313 | Non-Significant | FALSE |
| Brinp3 | -0.13604 | 7.441287 | -1.49421 | 0.163002 | 0.36744 | -5.22036 | Non-Significant | -0.33205 | 7.441287 | -3.66498 | 0.003669 | 0.037794 | -1.92139 | Significant | FALSE |
| Brk1 | -0.48934 | 7.648346 | -3.44193 | 0.005438 | 0.072094 | -2.11839 | Significant | -0.63863 | 7.648346 | -4.51049 | 0.000867 | 0.019835 | -0.4975 | Significant | FALSE |
| Brsk1 | 0.031316 | 8.835664 | 0.382201 | 0.70953 | 0.842149 | -6.29091 | Non-Significant | 0.026206 | 8.835664 | 0.320879 | 0.754264 | 0.854811 | -6.60735 | Non-Significant | FALSE |
| Brsk2 | 0.059433 | 8.426912 | 0.639937 | 0.535203 | 0.722416 | -6.12658 | Non-Significant | 0.148884 | 8.426912 | 1.619662 | 0.13335 | 0.289863 | -5.38044 | Non-Significant | FALSE |
| Bsdc1 | 0.158898 | 7.225935 | 1.31686 | 0.214432 | 0.430324 | -5.40561 | Non-Significant | 0.044204 | 7.225935 | 0.369544 | 0.718678 | 0.831369 | -6.47505 | Non-Significant | FALSE |
| Bsg | 0.281363 | 9.308071 | 3.152023 | 0.009116 | 0.092417 | -2.65885 | Significant | 0.218883 | 9.308071 | 2.446649 | 0.032263 | 0.123782 | -4.10546 | Non-Significant | FALSE |
| Btbd1 | -0.11816 | 8.924202 | -1.28262 | 0.225768 | 0.443584 | -5.56054 | Non-Significant | -0.09351 | 8.924202 | -1.01785 | 0.330427 | 0.512599 | -6.13432 | Non-Significant | FALSE |
| Btbd10 | -0.33902 | 7.700064 | -2.47911 | 0.030451 | 0.15533 | -3.74725 | Non-Significant | -0.42206 | 7.700064 | -3.1034 | 0.009946 | 0.06194 | -2.91537 | Non-Significant | FALSE |
| Btbd2 | 0.258708 | 7.190802 | 2.789028 | 0.017485 | 0.123548 | -3.17755 | Non-Significant | 0.297381 | 7.190802 | 3.240636 | 0.007781 | 0.0543 | -2.62171 | Non-Significant | FALSE |
| Bub3 | 0.130969 | 6.943071 | 1.223668 | 0.246424 | 0.467038 | -5.47131 | Non-Significant | 0.182492 | 6.943071 | 1.743739 | 0.108805 | 0.256721 | -5.08589 | Non-Significant | FALSE |
| Bzw1 | -0.07542 | 8.795152 | -0.68905 | 0.504945 | 0.700281 | -6.12145 | Non-Significant | -0.08358 | 8.795152 | -0.76701 | 0.459086 | 0.633824 | -6.35579 | Non-Significant | FALSE |
| C1ql3 | 0.323339 | 7.416356 | 3.257857 | 0.007545 | 0.086349 | -2.40067 | Significant | 0.432342 | 7.416356 | 4.402339 | 0.001037 | 0.021699 | -0.64953 | Significant | FALSE |
| C1qtnf4 | 0.334249 | 9.004173 | 2.229537 | 0.047363 | 0.190019 | -4.20819 | Non-Significant | 0.242416 | 9.004173 | 1.617501 | 0.133818 | 0.28994 | -5.40374 | Non-Significant | FALSE |
| C2cd2l | 0.51119 | 8.183698 | 5.652931 | 0.000143 | 0.023095 | 1.351023 | Significant | 0.528047 | 8.183698 | 5.899837 | 9.96E-05 | 0.007744 | 1.647056 | Significant | FALSE |
| Cab39 | -0.22667 | 10.1314 | -2.79353 | 0.017344 | 0.123509 | -3.27381 | Non-Significant | -0.1425 | 10.1314 | -1.74202 | 0.109115 | 0.257 | -5.22298 | Non-Significant | FALSE |
| Cables1 | -0.20333 | 6.822287 | -1.85248 | 0.090721 | 0.267694 | -4.64855 | Non-Significant | -0.24525 | 6.822287 | -2.29437 | 0.042256 | 0.14379 | -4.23133 | Non-Significant | FALSE |
| Cabp1 | 0.112948 | 8.174084 | 0.723661 | 0.48425 | 0.684023 | -6.05425 | Non-Significant | -0.02593 | 8.174084 | -0.16755 | 0.869955 | 0.931896 | -6.60837 | Non-Significant | FALSE |
| Cacna1i | -0.27506 | 8.338531 | -1.82536 | 0.094955 | 0.274018 | -4.80751 | Non-Significant | -0.15735 | 8.338531 | -1.05874 | 0.312222 | 0.494166 | -6.07318 | Non-Significant | FALSE |
| Cacnb1 | 0.184542 | 7.653694 | 2.127013 | 0.05666 | 0.208349 | -4.29855 | Non-Significant | 0.448792 | 7.653694 | 5.257245 | 0.000262 | 0.012089 | 0.699018 | Significant | TRUE |
| Cacnb2 | 0.112036 | 6.739256 | 1.118252 | 0.287083 | 0.510181 | -5.56374 | Non-Significant | 0.017559 | 6.739256 | 0.179398 | 0.860859 | 0.925856 | -6.47513 | Non-Significant | FALSE |
| Cacnb3 | 0.154297 | 8.462052 | 1.560107 | 0.146781 | 0.34938 | -5.17978 | Non-Significant | 0.258213 | 8.462052 | 2.634746 | 0.023059 | 0.101212 | -3.76322 | Non-Significant | TRUE |
| Cacng7 | 0.360096 | 6.75938 | 2.994092 | 0.012099 | 0.105315 | -2.79945 | Non-Significant | 0.179829 | 6.75938 | 1.520142 | 0.156443 | 0.321179 | -5.36763 | Non-Significant | FALSE |
| Cacng8 | 0.305377 | 6.971734 | 2.374479 | 0.036675 | 0.170529 | -3.83872 | Non-Significant | 0.354438 | 6.971734 | 2.805065 | 0.016989 | 0.08488 | -3.35849 | Non-Significant | FALSE |
| Cacul1 | -0.18774 | 7.75068 | -2.23617 | 0.046815 | 0.189667 | -4.14196 | Non-Significant | -0.10632 | 7.75068 | -1.28068 | 0.226426 | 0.402773 | -5.7868 | Non-Significant | FALSE |
| Cadm1 | 0.232093 | 8.585879 | 2.834052 | 0.016127 | 0.118001 | -3.18356 | Non-Significant | 0.292367 | 8.585879 | 3.592616 | 0.004167 | 0.040558 | -2.0931 | Significant | FALSE |
| Cadm4 | 0.054322 | 9.363061 | 0.46931 | 0.647934 | 0.799616 | -6.2607 | Non-Significant | -0.20511 | 9.363061 | -1.77097 | 0.103993 | 0.249364 | -5.18855 | Non-Significant | FALSE |
| Cadps2 | -0.03174 | 9.027675 | -0.43088 | 0.674805 | 0.817705 | -6.27501 | Non-Significant | -0.02897 | 9.027675 | -0.39379 | 0.701195 | 0.821673 | -6.58244 | Non-Significant | FALSE |
| Calm1 | 0.155602 | 12.6999 | 1.61495 | 0.134372 | 0.330986 | -5.0552 | Non-Significant | 0.101989 | 12.6999 | 1.048879 | 0.31654 | 0.498184 | -6.00474 | Non-Significant | FALSE |
| Calm3 | 0.572607 | 11.83573 | 4.561847 | 0.000796 | 0.036195 | -0.28182 | Significant | 0.434584 | 11.83573 | 3.437869 | 0.005477 | 0.046269 | -2.333 | Non-Significant | FALSE |
| Calr | -0.12866 | 9.596463 | -1.2378 | 0.241338 | 0.460808 | -5.61834 | Non-Significant | 0.539879 | 9.596463 | 5.134051 | 0.000317 | 0.013486 | 0.472723 | Significant | FALSE |
| Caly | 0.268323 | 7.551884 | 2.053699 | 0.064342 | 0.224067 | -4.41303 | Non-Significant | 0.120228 | 7.551884 | 0.923565 | 0.375358 | 0.56079 | -6.13811 | Non-Significant | FALSE |
| Camk1 | -0.08315 | 7.58735 | -0.64417 | 0.532556 | 0.720807 | -6.07751 | Non-Significant | -0.00559 | 7.58735 | -0.04372 | 0.965906 | 0.983195 | -6.59129 | Non-Significant | FALSE |
| Camk2a | 0.033949 | 10.59472 | 0.276431 | 0.787298 | 0.883648 | -6.32072 | Non-Significant | -0.29125 | 10.59472 | -2.36678 | 0.037179 | 0.134184 | -4.22846 | Non-Significant | TRUE |
| Camk2n1 | 0.232229 | 11.55612 | 2.071845 | 0.062354 | 0.220606 | -4.42352 | Non-Significant | -0.00528 | 11.55612 | -0.04693 | 0.963407 | 0.981736 | -6.61416 | Non-Significant | FALSE |
| Camkk1 | 0.035957 | 8.515936 | 0.450572 | 0.660974 | 0.807891 | -6.24251 | Non-Significant | 0.042437 | 8.515936 | 0.535926 | 0.602586 | 0.74638 | -6.49565 | Non-Significant | FALSE |
| Camkv | 0.243064 | 10.58084 | 2.257934 | 0.045057 | 0.186209 | -4.15863 | Non-Significant | 0.346661 | 10.58084 | 3.185252 | 0.00859 | 0.05751 | -2.80901 | Non-Significant | FALSE |
| Camsap1 | -0.13092 | 8.611321 | -1.12524 | 0.284237 | 0.506445 | -5.72071 | Non-Significant | 0.125807 | 8.611321 | 1.091894 | 0.298018 | 0.477328 | -6.04792 | Non-Significant | FALSE |
| Camsap2 | -0.07877 | 9.858221 | -0.83658 | 0.420478 | 0.633318 | -6.01865 | Non-Significant | 0.011363 | 9.858221 | 0.119845 | 0.906748 | 0.953935 | -6.65483 | Non-Significant | FALSE |
| Camta2 | 0.143839 | 7.916491 | 0.745794 | 0.471295 | 0.673494 | -6.02156 | Non-Significant | 0.212774 | 7.916491 | 1.117712 | 0.287304 | 0.46772 | -5.97768 | Non-Significant | FALSE |
| Cand1 | 0.008127 | 8.927489 | 0.080093 | 0.93759 | 0.966206 | -6.3672 | Non-Significant | -0.0002 | 8.927489 | -0.00194 | 0.998488 | 0.999936 | -6.66418 | Non-Significant | FALSE |
| Cant1 | 0.232897 | 7.09695 | 2.365603 | 0.037256 | 0.170843 | -3.86931 | Non-Significant | 0.257839 | 7.09695 | 2.655101 | 0.022233 | 0.099233 | -3.63076 | Non-Significant | FALSE |
| Cap1 | 0.071939 | 8.756545 | 0.757847 | 0.464332 | 0.669515 | -6.06686 | Non-Significant | 0.102699 | 8.756545 | 1.086626 | 0.300241 | 0.47917 | -6.05736 | Non-Significant | FALSE |
| Cap2 | 0.050655 | 8.447556 | 0.471285 | 0.646566 | 0.799033 | -6.22033 | Non-Significant | 0.300933 | 8.447556 | 2.833969 | 0.016129 | 0.082394 | -3.4189 | Non-Significant | FALSE |
| Capns1 | 0.28141 | 7.10485 | 2.208155 | 0.049173 | 0.195379 | -4.12274 | Non-Significant | 0.228517 | 7.10485 | 1.81412 | 0.096762 | 0.240156 | -4.99177 | Non-Significant | FALSE |
| Caprin1 | -0.07267 | 8.81917 | -0.81991 | 0.429527 | 0.640848 | -6.02121 | Non-Significant | 0.07116 | 8.81917 | 0.806662 | 0.436808 | 0.615742 | -6.32332 | Non-Significant | FALSE |
| Capzb | 0.207625 | 7.201636 | 2.570095 | 0.025887 | 0.141478 | -3.55585 | Non-Significant | -0.08027 | 7.201636 | -1.00058 | 0.33835 | 0.522167 | -6.03572 | Non-Significant | FALSE |
| Car10 | -0.06111 | 7.795845 | -0.64484 | 0.532135 | 0.720807 | -6.09178 | Non-Significant | -0.15716 | 7.795845 | -1.66947 | 0.122961 | 0.275397 | -5.27599 | Non-Significant | FALSE |
| Car11 | 0.119822 | 7.914196 | 0.774765 | 0.454671 | 0.661916 | -6.00382 | Non-Significant | -0.07574 | 7.914196 | -0.49268 | 0.631844 | 0.767731 | -6.47984 | Non-Significant | FALSE |
| Car2 | 0.153908 | 7.891748 | 1.204164 | 0.253582 | 0.47489 | -5.5915 | Non-Significant | 0.083194 | 7.891748 | 0.657125 | 0.524495 | 0.689881 | -6.38189 | Non-Significant | TRUE |
| Carmil1 | -0.23048 | 6.919003 | -1.97719 | 0.073394 | 0.240574 | -4.4741 | Non-Significant | -0.21398 | 6.919003 | -1.88655 | 0.085647 | 0.220261 | -4.8903 | Non-Significant | FALSE |
| Casc4 | -0.00083 | 7.297009 | -0.00872 | 0.993198 | 0.996547 | -6.27197 | Non-Significant | -0.26639 | 7.297009 | -2.83294 | 0.016159 | 0.082394 | -3.36104 | Non-Significant | FALSE |
| Casd1 | 0.047503 | 7.413995 | 0.490516 | 0.633324 | 0.79146 | -6.15269 | Non-Significant | -0.0829 | 7.413995 | -0.86057 | 0.407685 | 0.591232 | -6.19015 | Non-Significant | FALSE |
| Cask | -0.04867 | 8.372808 | -0.60064 | 0.560146 | 0.742444 | -6.15691 | Non-Significant | -0.20897 | 8.372808 | -2.59795 | 0.024629 | 0.105518 | -3.82997 | Non-Significant | FALSE |
| Cbarp | 0.262607 | 8.818765 | 1.708617 | 0.115306 | 0.308173 | -4.9967 | Non-Significant | 0.207559 | 8.818765 | 1.353693 | 0.202763 | 0.376892 | -5.74856 | Non-Significant | FALSE |
| Cbl | -0.15581 | 7.012278 | -0.89697 | 0.388777 | 0.601818 | -5.826 | Non-Significant | -0.22264 | 7.012278 | -1.30996 | 0.216679 | 0.392336 | -5.69278 | Non-Significant | FALSE |
| Cbln2 | -0.19567 | 7.167965 | -1.38408 | 0.193536 | 0.401464 | -5.31437 | Non-Significant | 0.10291 | 7.167965 | 0.744952 | 0.471783 | 0.64297 | -6.2617 | Non-Significant | FALSE |
| Cbx4 | 0.325703 | 6.714269 | 3.19357 | 0.008463 | 0.090802 | -2.45361 | Significant | 0.435241 | 6.714269 | 4.395834 | 0.001048 | 0.021766 | -0.6148 | Significant | FALSE |
| Ccar2 | 0.16069 | 7.641188 | 1.866328 | 0.088626 | 0.265185 | -4.70308 | Non-Significant | 0.1782 | 7.641188 | 2.088303 | 0.060601 | 0.177869 | -4.62878 | Non-Significant | FALSE |
| Ccdc136 | 0.161022 | 7.077401 | 0.994399 | 0.341217 | 0.563019 | -5.73932 | Non-Significant | -0.08911 | 7.077401 | -0.55605 | 0.58921 | 0.74063 | -6.37218 | Non-Significant | FALSE |
| Ccdc149 | -0.1843 | 7.295453 | -1.99411 | 0.071295 | 0.237473 | -4.4945 | Non-Significant | -0.25623 | 7.295453 | -2.80351 | 0.017036 | 0.08496 | -3.41519 | Non-Significant | FALSE |
| Ccdc32 | -0.15046 | 6.886689 | -0.91643 | 0.378926 | 0.596509 | -5.78983 | Non-Significant | -0.19649 | 6.886689 | -1.22665 | 0.245344 | 0.425263 | -5.77437 | Non-Significant | FALSE |
| Ccdc85b | -0.04591 | 7.338609 | -0.21942 | 0.830309 | 0.910786 | -6.257 | Non-Significant | -0.41787 | 7.338609 | -2.00612 | 0.06984 | 0.193954 | -4.74674 | Non-Significant | FALSE |
| Ccdc91 | -0.09318 | 7.275767 | -0.67743 | 0.512011 | 0.705529 | -6.03172 | Non-Significant | -0.21692 | 7.275767 | -1.59295 | 0.139236 | 0.29806 | -5.34897 | Non-Significant | FALSE |
| Cck | 0.095947 | 7.86212 | 0.593081 | 0.565018 | 0.745548 | -6.12674 | Non-Significant | -0.0754 | 7.86212 | -0.46902 | 0.648136 | 0.780796 | -6.48949 | Non-Significant | FALSE |
| Ccm2 | -0.09195 | 7.559784 | -0.81116 | 0.434329 | 0.646185 | -5.95549 | Non-Significant | -0.07445 | 7.559784 | -0.66231 | 0.521293 | 0.686006 | -6.36029 | Non-Significant | FALSE |
| Ccnd2 | -0.08029 | 6.824064 | -0.49048 | 0.633345 | 0.79146 | -6.08311 | Non-Significant | -0.27398 | 6.824064 | -1.70586 | 0.11583 | 0.265123 | -5.1405 | Non-Significant | FALSE |
| Ccng1 | -0.05746 | 6.991427 | -0.37955 | 0.711442 | 0.842282 | -6.1447 | Non-Significant | -0.01027 | 6.991427 | -0.06954 | 0.945799 | 0.973063 | -6.53036 | Non-Significant | FALSE |
| Ccni | 0.308024 | 8.289543 | 2.886399 | 0.01468 | 0.112151 | -3.08318 | Non-Significant | 0.0292 | 8.289543 | 0.274847 | 0.788484 | 0.875439 | -6.5863 | Non-Significant | FALSE |
| Ccny | 0.013517 | 8.982963 | 0.200199 | 0.844947 | 0.916533 | -6.35094 | Non-Significant | -0.06671 | 8.982963 | -0.9897 | 0.343409 | 0.527928 | -6.16202 | Non-Significant | FALSE |
| Ccp110 | -0.12361 | 6.779631 | -0.91127 | 0.381522 | 0.597417 | -5.77235 | Non-Significant | -0.10381 | 6.779631 | -0.79043 | 0.445841 | 0.622504 | -6.18646 | Non-Significant | FALSE |
| Ccsap | 0.230508 | 7.095812 | 2.355571 | 0.037924 | 0.172722 | -3.88987 | Non-Significant | 0.129275 | 7.095812 | 1.337417 | 0.207852 | 0.384412 | -5.64581 | Non-Significant | FALSE |
| Ccser2 | -0.10184 | 8.32796 | -1.26398 | 0.232142 | 0.451143 | -5.55271 | Non-Significant | -0.17289 | 8.32796 | -2.16715 | 0.052831 | 0.163374 | -4.54851 | Non-Significant | FALSE |
| Cct2 | -0.01227 | 8.674911 | -0.11639 | 0.909421 | 0.952084 | -6.35327 | Non-Significant | -0.00123 | 8.674911 | -0.01172 | 0.990858 | 0.996065 | -6.65755 | Non-Significant | FALSE |
| Cct4 | -0.15343 | 7.870727 | -1.70921 | 0.115192 | 0.308173 | -4.94597 | Non-Significant | -0.0463 | 7.870727 | -0.52265 | 0.61149 | 0.751523 | -6.4669 | Non-Significant | FALSE |
| Cct5 | -0.04531 | 8.521824 | -0.4088 | 0.690461 | 0.829692 | -6.2587 | Non-Significant | 0.161445 | 8.521824 | 1.472176 | 0.168758 | 0.337265 | -5.58824 | Non-Significant | FALSE |
| Cct8 | -0.02521 | 8.635043 | -0.21824 | 0.831203 | 0.910786 | -6.33438 | Non-Significant | -0.07179 | 8.635043 | -0.6253 | 0.544421 | 0.705691 | -6.45081 | Non-Significant | FALSE |
| Cd200 | 0.182621 | 7.663886 | 1.370664 | 0.197566 | 0.406048 | -5.38405 | Non-Significant | 0.068595 | 7.663886 | 0.517602 | 0.614897 | 0.753981 | -6.45006 | Non-Significant | FALSE |
| Cd2bp2 | 0.100409 | 7.266658 | 1.010414 | 0.333821 | 0.55734 | -5.74162 | Non-Significant | 0.066196 | 7.266658 | 0.672467 | 0.515045 | 0.680794 | -6.31612 | Non-Significant | FALSE |
| Cd81 | 0.378516 | 9.112256 | 5.00897 | 0.000386 | 0.031358 | 0.39859 | Significant | 0.369641 | 9.112256 | 4.88563 | 0.00047 | 0.016359 | 0.075608 | Significant | FALSE |
| Cd99l2 | 0.079781 | 7.716439 | 0.828911 | 0.424625 | 0.636344 | -5.94781 | Non-Significant | 0.108799 | 7.716439 | 1.141565 | 0.27767 | 0.458729 | -5.94015 | Non-Significant | FALSE |
| Cdc16 | 0.159413 | 6.861521 | 1.623225 | 0.132585 | 0.329669 | -4.96144 | Non-Significant | 0.188385 | 6.861521 | 1.964352 | 0.075025 | 0.202061 | -4.74431 | Non-Significant | FALSE |
| Cdc23 | -0.06421 | 6.952467 | -0.67478 | 0.513629 | 0.706471 | -5.97893 | Non-Significant | 0.008073 | 6.952467 | 0.087095 | 0.932148 | 0.967153 | -6.52302 | Non-Significant | FALSE |
| Cdc34 | 0.025564 | 6.801213 | 0.214254 | 0.834235 | 0.913362 | -6.16862 | Non-Significant | -0.01991 | 6.801213 | -0.17104 | 0.867273 | 0.930139 | -6.48996 | Non-Significant | FALSE |
| Cdc37 | -0.09542 | 6.91222 | -0.99169 | 0.342479 | 0.563756 | -5.72398 | Non-Significant | -0.18665 | 6.91222 | -1.98221 | 0.072766 | 0.198479 | -4.73827 | Non-Significant | FALSE |
| Cdc42 | 0.099548 | 8.769194 | 0.772836 | 0.455765 | 0.662616 | -6.05898 | Non-Significant | -0.09724 | 8.769194 | -0.75713 | 0.464744 | 0.636947 | -6.36131 | Non-Significant | FALSE |
| Cdc42bpa | -0.27207 | 9.559751 | -2.51717 | 0.028453 | 0.149995 | -3.74405 | Non-Significant | -0.30224 | 9.559751 | -2.78771 | 0.017527 | 0.086919 | -3.52418 | Non-Significant | FALSE |
| Cdc42bpb | -0.13608 | 9.009495 | -1.26703 | 0.231091 | 0.450082 | -5.57985 | Non-Significant | -0.05261 | 9.009495 | -0.49069 | 0.633202 | 0.767987 | -6.53743 | Non-Significant | FALSE |
| Cdc42se1 | 0.299437 | 6.999585 | 2.872424 | 0.015053 | 0.113333 | -3.02332 | Non-Significant | 0.22377 | 6.999585 | 2.179628 | 0.051691 | 0.162006 | -4.4145 | Non-Significant | FALSE |
| Cdc42se2 | 0.322974 | 8.796225 | 3.817087 | 0.002813 | 0.052686 | -1.5188 | Significant | 0.054518 | 8.796225 | 0.645605 | 0.531568 | 0.694866 | -6.43809 | Non-Significant | FALSE |
| Cdh10 | -0.1202 | 6.803491 | -0.92764 | 0.373331 | 0.591937 | -5.75993 | Non-Significant | -0.07459 | 6.803491 | -0.59503 | 0.563761 | 0.721679 | -6.32773 | Non-Significant | FALSE |
| Cdh13 | 0.454846 | 7.036858 | 4.690063 | 0.000645 | 0.036195 | -0.05656 | Significant | 0.508995 | 7.036858 | 5.316973 | 0.000239 | 0.011748 | 0.826383 | Significant | FALSE |
| Cdh2 | 0.117222 | 7.539292 | 0.883074 | 0.395924 | 0.60779 | -5.88722 | Non-Significant | 0.116111 | 7.539292 | 0.881632 | 0.39667 | 0.582392 | -6.1799 | Non-Significant | FALSE |
| Cdh22 | -0.01951 | 7.698432 | -0.17856 | 0.861502 | 0.924833 | -6.27953 | Non-Significant | 0.167291 | 7.698432 | 1.553216 | 0.148409 | 0.310112 | -5.43447 | Non-Significant | FALSE |
| Cdh8 | -0.14789 | 7.300046 | -1.91534 | 0.081563 | 0.251922 | -4.6126 | Non-Significant | -0.20666 | 7.300046 | -2.70627 | 0.020284 | 0.094968 | -3.58203 | Non-Significant | TRUE |
| Cdip1 | 0.138356 | 8.232964 | 1.634077 | 0.130266 | 0.325578 | -5.06576 | Non-Significant | 0.151186 | 8.232964 | 1.805931 | 0.098098 | 0.241458 | -5.09822 | Non-Significant | FALSE |
| Cdipt | 0.23307 | 7.937473 | 2.661891 | 0.021964 | 0.132777 | -3.44723 | Non-Significant | 0.151091 | 7.937473 | 1.742391 | 0.109048 | 0.257 | -5.17683 | Non-Significant | FALSE |
| Cdk10 | -0.02549 | 6.787507 | -0.28188 | 0.783228 | 0.882402 | -6.16456 | Non-Significant | -0.3217 | 6.787507 | -1.68134 | 0.120594 | 0.272761 | -5.16658 | Non-Significant | FALSE |
| Cdk13 | -0.1851 | 7.892617 | -1.92377 | 0.080403 | 0.24981 | -4.63223 | Non-Significant | 0.016013 | 7.892617 | 0.169043 | 0.868804 | 0.931036 | -6.59835 | Non-Significant | FALSE |
| Cdk14 | 0.129412 | 8.509825 | 1.163122 | 0.269181 | 0.495101 | -5.67619 | Non-Significant | -0.03477 | 8.509825 | -0.31419 | 0.759201 | 0.858904 | -6.6097 | Non-Significant | FALSE |
| Cdk16 | 0.007337 | 9.036315 | 0.081478 | 0.936513 | 0.966206 | -6.36898 | Non-Significant | 0.006158 | 9.036315 | 0.06845 | 0.946645 | 0.973063 | -6.66231 | Non-Significant | FALSE |
| Cdk2ap1 | 0.241514 | 6.689053 | 1.448532 | 0.175128 | 0.38217 | -5.16172 | Non-Significant | 0.2835 | 6.689053 | 1.752118 | 0.107303 | 0.254524 | -5.03806 | Non-Significant | FALSE |
| Cdk5 | 0.124804 | 7.594573 | 0.978435 | 0.348708 | 0.57011 | -5.80776 | Non-Significant | 0.035354 | 7.594573 | 0.278676 | 0.785618 | 0.874433 | -6.54496 | Non-Significant | TRUE |
| Cdk5r1 | 0.155598 | 9.781102 | 2.179809 | 0.051675 | 0.19713 | -4.2936 | Non-Significant | 0.208033 | 9.781102 | 2.896071 | 0.014427 | 0.077428 | -3.32955 | Non-Significant | FALSE |
| Cdk5r2 | 0.392782 | 7.667723 | 4.126134 | 0.001652 | 0.042216 | -0.97022 | Significant | 0.279017 | 7.667723 | 2.949021 | 0.013119 | 0.073408 | -3.17285 | Non-Significant | FALSE |
| Cdkl5 | -0.17578 | 9.387306 | -1.81625 | 0.096417 | 0.275875 | -4.85456 | Non-Significant | -0.33098 | 9.387306 | -3.4165 | 0.005689 | 0.047313 | -2.42106 | Non-Significant | FALSE |
| Cds1 | -0.02459 | 7.674987 | -0.24274 | 0.812633 | 0.900153 | -6.26667 | Non-Significant | 0.029439 | 7.674987 | 0.293496 | 0.774559 | 0.867532 | -6.55194 | Non-Significant | FALSE |
| Cdv3 | -0.20678 | 8.231265 | -2.34129 | 0.038895 | 0.175946 | -4.00015 | Non-Significant | -0.35203 | 8.231265 | -4.02093 | 0.001978 | 0.027167 | -1.3464 | Significant | FALSE |
| Celf4 | -0.0979 | 9.264953 | -1.21487 | 0.249633 | 0.470456 | -5.64304 | Non-Significant | -0.05831 | 9.264953 | -0.72257 | 0.484893 | 0.652286 | -6.39199 | Non-Significant | FALSE |
| Celf5 | 0.138148 | 7.336478 | 1.605609 | 0.136418 | 0.333328 | -5.04521 | Non-Significant | 0.151742 | 7.336478 | 1.779184 | 0.10258 | 0.247708 | -5.07294 | Non-Significant | FALSE |
| Celsr2 | 0.01313 | 9.386163 | 0.086144 | 0.932887 | 0.964211 | -6.37194 | Non-Significant | 0.029475 | 9.386163 | 0.192855 | 0.850557 | 0.918835 | -6.64471 | Non-Significant | FALSE |
| Cend1 | 0.350009 | 7.565998 | 3.543563 | 0.004543 | 0.064036 | -1.93431 | Significant | 0.161081 | 7.565998 | 1.635375 | 0.129991 | 0.285346 | -5.29675 | Non-Significant | FALSE |
| Cenpb | -0.06527 | 9.546476 | -0.45704 | 0.656457 | 0.803966 | -6.26864 | Non-Significant | -0.44932 | 9.546476 | -3.14658 | 0.009205 | 0.059948 | -2.89691 | Non-Significant | FALSE |
| Cep170 | -0.18013 | 7.739059 | -1.68475 | 0.119921 | 0.313622 | -4.97555 | Non-Significant | -0.10712 | 7.739059 | -1.01288 | 0.332693 | 0.515597 | -6.07881 | Non-Significant | FALSE |
| Cep170b | -0.15598 | 9.571407 | -1.18008 | 0.262646 | 0.48508 | -5.6851 | Non-Significant | -0.0467 | 9.571407 | -0.35164 | 0.731698 | 0.839477 | -6.59962 | Non-Significant | FALSE |
| Cep19 | -0.27491 | 7.453374 | -2.32246 | 0.040211 | 0.177124 | -3.98964 | Non-Significant | -0.2481 | 7.453374 | -2.11675 | 0.057681 | 0.173562 | -4.58047 | Non-Significant | FALSE |
| Cfl1 | 0.221701 | 9.326435 | 1.761293 | 0.10568 | 0.290567 | -4.93136 | Non-Significant | 0.030473 | 9.326435 | 0.241807 | 0.81334 | 0.893448 | -6.63341 | Non-Significant | FALSE |
| Cggbp1 | 0.139273 | 6.664729 | 1.092749 | 0.297659 | 0.52066 | -5.5743 | Non-Significant | 0.09429 | 6.664729 | 0.76009 | 0.463044 | 0.636187 | -6.17908 | Non-Significant | FALSE |
| Chd3 | -0.23801 | 10.96852 | -2.95357 | 0.013012 | 0.108158 | -2.98564 | Non-Significant | 0.036458 | 10.96852 | 0.445679 | 0.664399 | 0.79219 | -6.62585 | Non-Significant | FALSE |
| Chga | -0.02012 | 7.711986 | -0.08463 | 0.934063 | 0.965054 | -6.30084 | Non-Significant | -0.23372 | 7.711986 | -1.649 | 0.12714 | 0.281622 | -5.29854 | Non-Significant | FALSE |
| Chgb | -0.24407 | 8.308331 | -2.11322 | 0.058036 | 0.210274 | -4.37326 | Non-Significant | -0.46876 | 8.308331 | -4.08765 | 0.001765 | 0.026016 | -1.23743 | Significant | FALSE |
| Chid1 | -0.07416 | 7.732984 | -0.63837 | 0.536184 | 0.722723 | -6.09666 | Non-Significant | -0.38918 | 7.732984 | -3.34796 | 0.006426 | 0.050402 | -2.48498 | Non-Significant | FALSE |
| Chmp1a | 0.175607 | 8.211571 | 1.38181 | 0.194214 | 0.401935 | -5.39764 | Non-Significant | 0.045113 | 8.211571 | 0.357973 | 0.727079 | 0.836391 | -6.55613 | Non-Significant | FALSE |
| Chmp2a | -0.12844 | 7.044616 | -0.80085 | 0.440031 | 0.647581 | -5.92037 | Non-Significant | -0.42867 | 7.044616 | -2.70205 | 0.020439 | 0.09519 | -3.56928 | Non-Significant | FALSE |
| Chmp3 | 0.021145 | 7.839635 | 0.143734 | 0.888288 | 0.941093 | -6.29808 | Non-Significant | -0.14012 | 7.839635 | -0.95802 | 0.358461 | 0.54235 | -6.13353 | Non-Significant | FALSE |
| Chmp4b | 0.134366 | 8.340129 | 0.93503 | 0.369674 | 0.588537 | -5.89148 | Non-Significant | 0.022665 | 8.340129 | 0.15894 | 0.876572 | 0.936362 | -6.62051 | Non-Significant | FALSE |
| Chmp7 | 0.036917 | 8.187275 | 0.339382 | 0.740658 | 0.862689 | -6.2656 | Non-Significant | -0.06592 | 8.187275 | -0.61172 | 0.553052 | 0.714844 | -6.429 | Non-Significant | FALSE |
| Chp1 | 0.121831 | 7.913815 | 1.380714 | 0.194541 | 0.402302 | -5.38357 | Non-Significant | 0.124611 | 7.913815 | 1.428679 | 0.180634 | 0.351809 | -5.60922 | Non-Significant | FALSE |
| Chpf | 0.03132 | 7.589971 | 0.330383 | 0.747264 | 0.867473 | -6.22915 | Non-Significant | 0.191447 | 7.589971 | 2.045701 | 0.065237 | 0.186849 | -4.69585 | Non-Significant | FALSE |
| Chpf2 | 0.24751 | 7.789295 | 2.392469 | 0.035523 | 0.168672 | -3.89171 | Non-Significant | -0.08262 | 7.789295 | -0.79943 | 0.440819 | 0.618069 | -6.26592 | Non-Significant | FALSE |
| Chrm1 | 0.188118 | 8.349895 | 2.200971 | 0.049796 | 0.19584 | -4.22355 | Non-Significant | 0.123695 | 8.349895 | 1.459239 | 0.172219 | 0.341631 | -5.59046 | Non-Significant | TRUE |
| Chrm3 | -0.1404 | 8.039811 | -1.5181 | 0.156951 | 0.361094 | -5.22058 | Non-Significant | -0.33829 | 8.039811 | -3.68268 | 0.003557 | 0.037213 | -1.91772 | Significant | TRUE |
| Chst1 | 0.237662 | 8.975012 | 3.0359 | 0.011225 | 0.102536 | -2.85474 | Non-Significant | 0.288963 | 8.975012 | 3.693235 | 0.003492 | 0.037027 | -1.93036 | Significant | FALSE |
| Chst10 | 0.109243 | 6.860943 | 1.145461 | 0.27612 | 0.498282 | -5.54184 | Non-Significant | 0.237237 | 6.860943 | 2.557396 | 0.026481 | 0.110289 | -3.77925 | Non-Significant | FALSE |
| Chst15 | -0.2719 | 7.987232 | -3.65823 | 0.003713 | 0.058838 | -1.759 | Significant | -0.12796 | 7.987232 | -1.74728 | 0.108167 | 0.25612 | -5.17877 | Non-Significant | FALSE |
| Chst2 | 0.027186 | 9.618585 | 0.321461 | 0.753835 | 0.870875 | -6.32302 | Non-Significant | -0.01787 | 9.618585 | -0.21048 | 0.837108 | 0.908336 | -6.64161 | Non-Significant | FALSE |
| Chtf8 | 0.029052 | 7.657354 | 0.329458 | 0.747944 | 0.867473 | -6.23204 | Non-Significant | 0.287899 | 7.657354 | 3.318893 | 0.006767 | 0.050947 | -2.53408 | Non-Significant | FALSE |
| Chtop | -0.02178 | 8.621429 | -0.20311 | 0.842723 | 0.915914 | -6.33591 | Non-Significant | -0.017 | 8.621429 | -0.1596 | 0.876064 | 0.93626 | -6.64198 | Non-Significant | FALSE |
| Ciao1 | 0.014336 | 6.821991 | 0.101215 | 0.921186 | 0.95603 | -6.19075 | Non-Significant | -0.02455 | 6.821991 | -0.17766 | 0.86219 | 0.926543 | -6.49199 | Non-Significant | FALSE |
| Cic | 0.094393 | 7.684121 | 0.567314 | 0.581796 | 0.757069 | -6.12906 | Non-Significant | 0.114592 | 7.684121 | 0.695101 | 0.501288 | 0.667885 | -6.34261 | Non-Significant | FALSE |
| Cinp | -0.33618 | 6.955307 | -2.60566 | 0.024291 | 0.138232 | -3.48089 | Non-Significant | -0.33728 | 6.955307 | -2.68916 | 0.020916 | 0.096419 | -3.58993 | Non-Significant | FALSE |
| Cipc | 0.104601 | 7.500455 | 1.322836 | 0.212502 | 0.427879 | -5.43715 | Non-Significant | -0.21916 | 7.500455 | -2.77132 | 0.01805 | 0.088045 | -3.47802 | Non-Significant | FALSE |
| Ckap5 | -0.07924 | 9.206064 | -0.79348 | 0.444135 | 0.651722 | -6.0504 | Non-Significant | 0.0307 | 9.206064 | 0.307044 | 0.764495 | 0.862028 | -6.6143 | Non-Significant | FALSE |
| Ckb | 0.136848 | 11.81119 | 1.345966 | 0.205166 | 0.41687 | -5.43986 | Non-Significant | 0.279444 | 11.81119 | 2.711723 | 0.020087 | 0.094373 | -3.59799 | Non-Significant | FALSE |
| Clasp1 | -0.28614 | 8.608206 | -2.91999 | 0.013821 | 0.111238 | -3.04738 | Non-Significant | -0.2432 | 8.608206 | -2.50373 | 0.029143 | 0.116929 | -4.00676 | Non-Significant | FALSE |
| Clasp2 | -0.07038 | 10.21483 | -0.78365 | 0.44965 | 0.658013 | -6.05693 | Non-Significant | 0.005014 | 10.21483 | 0.055353 | 0.956841 | 0.977277 | -6.65455 | Non-Significant | FALSE |
| Clcn3 | -0.03622 | 9.180653 | -0.54885 | 0.593981 | 0.765855 | -6.21731 | Non-Significant | -0.12451 | 9.180653 | -1.88659 | 0.08564 | 0.220261 | -5.01403 | Non-Significant | FALSE |
| Clcn6 | 0.115336 | 7.307662 | 1.339251 | 0.207274 | 0.419878 | -5.38449 | Non-Significant | 0.143003 | 7.307662 | 1.676748 | 0.121504 | 0.274357 | -5.21919 | Non-Significant | FALSE |
| Clcn7 | 0.211348 | 6.643882 | 2.373857 | 0.036715 | 0.170529 | -3.8094 | Non-Significant | 0.118427 | 6.643882 | 1.360284 | 0.200731 | 0.374304 | -5.56131 | Non-Significant | FALSE |
| Cldn11 | -0.04774 | 8.155594 | -0.22408 | 0.826769 | 0.909149 | -6.29537 | Non-Significant | 0.037464 | 8.155594 | 0.178279 | 0.861717 | 0.926406 | -6.60319 | Non-Significant | FALSE |
| Cldn5 | -0.2399 | 6.930546 | -2.00488 | 0.069989 | 0.235761 | -4.42814 | Non-Significant | -0.07001 | 6.930546 | -0.60538 | 0.557101 | 0.718013 | -6.34201 | Non-Significant | FALSE |
| Clec16a | -0.18609 | 7.740197 | -1.95665 | 0.076019 | 0.244703 | -4.57608 | Non-Significant | 0.021662 | 7.740197 | 0.231231 | 0.821344 | 0.89851 | -6.57702 | Non-Significant | FALSE |
| Clec2l | 0.178071 | 6.774219 | 1.34543 | 0.205334 | 0.416894 | -5.31518 | Non-Significant | 0.01715 | 6.774219 | 0.131882 | 0.897438 | 0.949937 | -6.48503 | Non-Significant | FALSE |
| Clmn | 0.044277 | 6.802954 | 0.219152 | 0.83051 | 0.910786 | -6.17189 | Non-Significant | -0.1108 | 6.802954 | -0.5592 | 0.587135 | 0.739988 | -6.34034 | Non-Significant | FALSE |
| Clock | -0.37658 | 8.117132 | -3.64633 | 0.003791 | 0.059032 | -1.78558 | Significant | -0.24716 | 8.117132 | -2.42923 | 0.033278 | 0.12499 | -4.10716 | Non-Significant | FALSE |
| Clptm1 | 0.157034 | 8.392707 | 1.828759 | 0.094414 | 0.273709 | -4.79758 | Non-Significant | 0.196769 | 8.392707 | 2.312979 | 0.040891 | 0.14148 | -4.30433 | Non-Significant | FALSE |
| Clptm1l | 0.25347 | 7.756072 | 2.960927 | 0.012841 | 0.107803 | -2.93155 | Non-Significant | 0.313238 | 7.756072 | 3.702175 | 0.003438 | 0.037027 | -1.86774 | Significant | FALSE |
| Clstn1 | 0.213362 | 11.11175 | 2.242312 | 0.046312 | 0.188638 | -4.16915 | Non-Significant | 0.191858 | 11.11175 | 1.997963 | 0.070825 | 0.195351 | -4.80826 | Non-Significant | TRUE |
| Clta | 0.257248 | 8.843079 | 1.761911 | 0.105572 | 0.290567 | -4.923 | Non-Significant | 0.103466 | 8.843079 | 0.710102 | 0.492294 | 0.658749 | -6.39515 | Non-Significant | FALSE |
| Cltb | -0.07612 | 8.383413 | -0.43379 | 0.672754 | 0.815958 | -6.24848 | Non-Significant | -0.27598 | 8.383413 | -1.58288 | 0.141512 | 0.30102 | -5.43575 | Non-Significant | FALSE |
| Clu | 0.514066 | 9.17219 | 3.667415 | 0.003654 | 0.058824 | -1.77479 | Significant | 0.566288 | 9.17219 | 4.02966 | 0.001949 | 0.02701 | -1.34687 | Significant | FALSE |
| Cmas | -0.1765 | 8.033312 | -1.14695 | 0.27553 | 0.497872 | -5.66556 | Non-Significant | -0.26005 | 8.033312 | -1.70608 | 0.115789 | 0.265123 | -5.23922 | Non-Significant | FALSE |
| Cmip | 0.183866 | 8.28523 | 1.149815 | 0.274397 | 0.497727 | -5.66593 | Non-Significant | 0.315539 | 8.28523 | 1.997947 | 0.070827 | 0.195351 | -4.80579 | Non-Significant | FALSE |
| Cnbp | -0.31146 | 8.205002 | -1.95661 | 0.076024 | 0.244703 | -4.6124 | Non-Significant | -0.60666 | 8.205002 | -3.82966 | 0.002753 | 0.033055 | -1.67626 | Significant | FALSE |
| Cnih2 | 0.347275 | 7.379735 | 2.973458 | 0.012556 | 0.106744 | -2.88136 | Non-Significant | 0.336615 | 7.379735 | 2.902586 | 0.01426 | 0.07699 | -3.22462 | Non-Significant | TRUE |
| Cnih3 | 0.274033 | 8.148302 | 2.979293 | 0.012425 | 0.105968 | -2.91669 | Non-Significant | 0.163652 | 8.148302 | 1.795309 | 0.099856 | 0.243325 | -5.10753 | Non-Significant | TRUE |
| Cnnm1 | 0.163409 | 7.309019 | 1.466086 | 0.17038 | 0.378026 | -5.22405 | Non-Significant | 0.211214 | 7.309019 | 1.913618 | 0.081803 | 0.213967 | -4.86665 | Non-Significant | FALSE |
| Cnot1 | -0.02929 | 8.523257 | -0.21037 | 0.837188 | 0.914292 | -6.32907 | Non-Significant | -0.07971 | 8.523257 | -0.57671 | 0.575648 | 0.731924 | -6.47513 | Non-Significant | FALSE |
| Cnot7 | -0.0449 | 6.768611 | -0.42035 | 0.682254 | 0.824188 | -6.10475 | Non-Significant | -0.22327 | 6.768611 | -2.22859 | 0.047441 | 0.155505 | -4.32532 | Non-Significant | FALSE |
| Cnot8 | 0.150425 | 7.020235 | 1.22045 | 0.247594 | 0.46793 | -5.49027 | Non-Significant | 0.078835 | 7.020235 | 0.650165 | 0.528816 | 0.693181 | -6.30458 | Non-Significant | FALSE |
| Cnp | -0.07938 | 9.613425 | -0.72195 | 0.485259 | 0.68432 | -6.10874 | Non-Significant | -0.08295 | 9.613425 | -0.75148 | 0.468002 | 0.63977 | -6.37058 | Non-Significant | FALSE |
| Cnpy3 | 0.288662 | 6.949176 | 2.369023 | 0.037031 | 0.170529 | -3.85425 | Non-Significant | 0.077877 | 6.949176 | 0.647433 | 0.530517 | 0.694053 | -6.29224 | Non-Significant | FALSE |
| Cnst | 0.005028 | 7.636354 | 0.064866 | 0.949435 | 0.973425 | -6.29146 | Non-Significant | 0.080916 | 7.636354 | 1.054852 | 0.313918 | 0.49568 | -6.02746 | Non-Significant | FALSE |
| Cntnap5a | 0.065045 | 7.250171 | 0.550115 | 0.59314 | 0.765139 | -6.09953 | Non-Significant | -0.05069 | 7.250171 | -0.43264 | 0.673564 | 0.799559 | -6.45591 | Non-Significant | FALSE |
| Cobl | -0.02549 | 7.367159 | -0.14663 | 0.886057 | 0.940491 | -6.24013 | Non-Significant | 0.428319 | 7.367159 | 2.520659 | 0.028277 | 0.114734 | -3.89229 | Non-Significant | FALSE |
| Cog3 | -0.11567 | 7.17421 | -1.37687 | 0.195693 | 0.403438 | -5.33475 | Non-Significant | -0.12884 | 7.17421 | -1.55759 | 0.147374 | 0.308818 | -5.38618 | Non-Significant | FALSE |
| Cog6 | -0.0612 | 7.326882 | -0.66525 | 0.519478 | 0.711204 | -6.03267 | Non-Significant | 0.12391 | 7.326882 | 1.366045 | 0.198969 | 0.372615 | -5.6439 | Non-Significant | FALSE |
| Cog7 | 0.067235 | 6.788 | 0.797868 | 0.441686 | 0.649196 | -5.86323 | Non-Significant | 0.06649 | 6.788 | 0.810798 | 0.434526 | 0.614394 | -6.16116 | Non-Significant | FALSE |
| Col25a1 | 0.073537 | 7.370401 | 0.616847 | 0.549782 | 0.734407 | -6.07122 | Non-Significant | 0.038032 | 7.370401 | 0.321442 | 0.753849 | 0.854811 | -6.51018 | Non-Significant | FALSE |
| Copa | -0.0354 | 8.943233 | -0.37664 | 0.71354 | 0.843516 | -6.29559 | Non-Significant | 0.091492 | 8.943233 | 0.975559 | 0.35007 | 0.534617 | -6.17429 | Non-Significant | FALSE |
| Copb2 | -0.08446 | 7.476276 | -0.85242 | 0.412002 | 0.625489 | -5.91175 | Non-Significant | -0.00269 | 7.476276 | -0.02744 | 0.978599 | 0.989803 | -6.58169 | Non-Significant | FALSE |
| Cope | 0.103455 | 7.162747 | 0.644237 | 0.532512 | 0.720807 | -6.03625 | Non-Significant | -0.14207 | 7.162747 | -0.89308 | 0.390773 | 0.576896 | -6.13787 | Non-Significant | FALSE |
| Copg1 | 0.135738 | 9.19995 | 1.902803 | 0.083319 | 0.254423 | -4.7217 | Non-Significant | 0.142884 | 9.19995 | 2.000246 | 0.070548 | 0.195351 | -4.83619 | Non-Significant | FALSE |
| Copg2 | 0.019885 | 7.924062 | 0.216463 | 0.832555 | 0.911894 | -6.28699 | Non-Significant | -0.08632 | 7.924062 | -0.94762 | 0.363504 | 0.548431 | -6.14863 | Non-Significant | FALSE |
| Cops2 | -0.48618 | 6.936694 | -3.92638 | 0.002328 | 0.049477 | -1.27684 | Significant | -0.659 | 6.936694 | -5.43457 | 0.000199 | 0.011476 | 0.986153 | Significant | FALSE |
| Cops3 | -0.08742 | 7.20676 | -0.74264 | 0.473125 | 0.675388 | -5.96578 | Non-Significant | 0.030853 | 7.20676 | 0.26634 | 0.794861 | 0.879953 | -6.51575 | Non-Significant | FALSE |
| Cops4 | -0.41789 | 7.680762 | -3.04992 | 0.010946 | 0.102057 | -2.78463 | Non-Significant | -0.43989 | 7.680762 | -3.23379 | 0.007876 | 0.054786 | -2.68701 | Non-Significant | FALSE |
| Cops7a | 0.392759 | 7.311149 | 4.443847 | 0.000968 | 0.036195 | -0.44609 | Significant | 0.328384 | 7.311149 | 3.737702 | 0.003231 | 0.03575 | -1.76471 | Significant | FALSE |
| Cops8 | 0.158661 | 7.191156 | 1.416244 | 0.184156 | 0.391405 | -5.27531 | Non-Significant | 0.158081 | 7.191156 | 1.427093 | 0.18108 | 0.35201 | -5.54362 | Non-Significant | FALSE |
| Coq8a | 0.127017 | 6.708644 | 0.677141 | 0.512185 | 0.705529 | -5.95249 | Non-Significant | -0.16747 | 6.708644 | -0.89949 | 0.387496 | 0.573104 | -6.06727 | Non-Significant | FALSE |
| Coro1a | 0.249715 | 8.293065 | 1.607104 | 0.136089 | 0.333009 | -5.10931 | Non-Significant | 0.043145 | 8.293065 | 0.279288 | 0.78516 | 0.874287 | -6.58634 | Non-Significant | FALSE |
| Cpd | 0.011703 | 8.370769 | 0.082628 | 0.935619 | 0.965916 | -6.33245 | Non-Significant | 0.080024 | 8.370769 | 0.57143 | 0.579098 | 0.733944 | -6.46569 | Non-Significant | FALSE |
| Cpe | 0.160103 | 11.34973 | 2.036719 | 0.066256 | 0.228875 | -4.48815 | Non-Significant | 0.251949 | 11.34973 | 3.167139 | 0.008873 | 0.058597 | -2.81972 | Non-Significant | FALSE |
| Cplx1 | 0.473379 | 8.34208 | 3.654047 | 0.00374 | 0.058923 | -1.77216 | Significant | 0.488132 | 8.34208 | 3.798275 | 0.002907 | 0.034091 | -1.7207 | Significant | TRUE |
| Cpne4 | 0.465008 | 6.914239 | 3.622573 | 0.003953 | 0.060111 | -1.75219 | Significant | 0.590657 | 6.914239 | 4.689876 | 0.000645 | 0.017563 | -0.14467 | Significant | FALSE |
| Cpne5 | 0.199321 | 6.91877 | 1.980617 | 0.072964 | 0.240048 | -4.45395 | Non-Significant | 0.214471 | 6.91877 | 2.175955 | 0.052024 | 0.162006 | -4.41471 | Non-Significant | FALSE |
| Cpne6 | 0.585619 | 7.079748 | 4.169986 | 0.001534 | 0.041073 | -0.87023 | Significant | 0.541719 | 7.079748 | 3.89285 | 0.002467 | 0.030805 | -1.47397 | Significant | FALSE |
| Cpsf2 | 0.100977 | 7.10997 | 1.24922 | 0.237292 | 0.457172 | -5.4651 | Non-Significant | 0.162185 | 7.10997 | 2.038649 | 0.066036 | 0.187334 | -4.65616 | Non-Significant | FALSE |
| Creb3 | 0.026652 | 7.022924 | 0.13727 | 0.893276 | 0.942182 | -6.22748 | Non-Significant | -0.26127 | 7.022924 | -1.3599 | 0.200849 | 0.374304 | -5.62548 | Non-Significant | FALSE |
| Creg2 | 0.009646 | 7.837747 | 0.10605 | 0.917436 | 0.955922 | -6.30451 | Non-Significant | -0.25784 | 7.837747 | -2.84221 | 0.015893 | 0.081962 | -3.3742 | Non-Significant | FALSE |
| Creld1 | 0.142499 | 7.074164 | 1.490142 | 0.164051 | 0.368565 | -5.15973 | Non-Significant | 0.328605 | 7.074164 | 3.501103 | 0.004897 | 0.043715 | -2.16202 | Significant | FALSE |
| Crim1 | 0.236518 | 7.655202 | 1.978302 | 0.073254 | 0.240411 | -4.53465 | Non-Significant | 0.221569 | 7.655202 | 1.868477 | 0.088305 | 0.225367 | -4.96939 | Non-Significant | FALSE |
| Crip2 | 0.291736 | 6.786986 | 2.244813 | 0.046109 | 0.188519 | -4.02218 | Non-Significant | 0.388073 | 6.786986 | 3.067424 | 0.010608 | 0.063695 | -2.88653 | Non-Significant | FALSE |
| Crmp1 | 0.257423 | 7.791993 | 2.406994 | 0.034619 | 0.166146 | -3.86342 | Non-Significant | 0.213028 | 7.791993 | 2.010562 | 0.069308 | 0.193543 | -4.76162 | Non-Significant | FALSE |
| Cry2 | 0.111012 | 7.695492 | 1.31387 | 0.215403 | 0.430324 | -5.45676 | Non-Significant | -0.03128 | 7.695492 | -0.37203 | 0.716878 | 0.830039 | -6.52187 | Non-Significant | FALSE |
| Crym | 0.09828 | 7.545429 | 0.56141 | 0.585677 | 0.760642 | -6.11948 | Non-Significant | 0.186163 | 7.545429 | 1.074461 | 0.305423 | 0.485346 | -5.99646 | Non-Significant | FALSE |
| Cse1l | 0.077479 | 6.930348 | 0.833224 | 0.42229 | 0.634782 | -5.85787 | Non-Significant | 0.012093 | 6.930348 | 0.132751 | 0.896767 | 0.94989 | -6.50964 | Non-Significant | FALSE |
| Csnk1d | -0.08342 | 8.110498 | -1.02324 | 0.327985 | 0.552417 | -5.79149 | Non-Significant | 0.039169 | 8.110498 | 0.487341 | 0.635501 | 0.770078 | -6.49756 | Non-Significant | FALSE |
| Csnk1e | 0.03733 | 7.829493 | 0.421788 | 0.681235 | 0.823633 | -6.21115 | Non-Significant | 0.145758 | 7.829493 | 1.668845 | 0.123086 | 0.275397 | -5.28033 | Non-Significant | FALSE |
| Csnk1g2 | -0.20901 | 8.607118 | -2.76711 | 0.018187 | 0.124205 | -3.30724 | Non-Significant | -0.16746 | 8.607118 | -2.23547 | 0.046872 | 0.153973 | -4.45428 | Non-Significant | FALSE |
| Csnk1g3 | -0.09217 | 6.907891 | -0.95724 | 0.358837 | 0.579918 | -5.75804 | Non-Significant | -0.24463 | 6.907891 | -2.59125 | 0.024925 | 0.106327 | -3.74129 | Non-Significant | FALSE |
| Csnk2a1 | -0.04679 | 8.601846 | -0.60299 | 0.558638 | 0.741794 | -6.17035 | Non-Significant | -0.16677 | 8.601846 | -2.16159 | 0.053346 | 0.164207 | -4.5721 | Non-Significant | FALSE |
| Csnk2b | 0.11874 | 6.883567 | 0.747185 | 0.470488 | 0.673419 | -5.92291 | Non-Significant | -0.06067 | 6.883567 | -0.38808 | 0.705296 | 0.823677 | -6.43289 | Non-Significant | FALSE |
| Cspg5 | 0.253806 | 9.660205 | 2.628497 | 0.023318 | 0.135721 | -3.55375 | Non-Significant | 0.049998 | 9.660205 | 0.5166 | 0.615574 | 0.754121 | -6.52306 | Non-Significant | TRUE |
| Csrnp2 | -0.18503 | 7.133027 | -1.986 | 0.072294 | 0.238723 | -4.48581 | Non-Significant | -0.15372 | 7.133027 | -1.6826 | 0.120344 | 0.272442 | -5.2116 | Non-Significant | FALSE |
| Csrnp3 | -0.18315 | 8.04362 | -1.98156 | 0.072847 | 0.239957 | -4.55732 | Non-Significant | -0.33032 | 8.04362 | -3.60271 | 0.004094 | 0.040303 | -2.05793 | Significant | FALSE |
| Csrp1 | 0.336414 | 7.895677 | 2.766426 | 0.018209 | 0.124205 | -3.2698 | Non-Significant | 0.161045 | 7.895677 | 1.333997 | 0.208935 | 0.385616 | -5.72376 | Non-Significant | FALSE |
| Cstf2t | -0.03981 | 7.793123 | -0.5177 | 0.614828 | 0.778492 | -6.1639 | Non-Significant | 0.066732 | 7.793123 | 0.878907 | 0.398084 | 0.583436 | -6.20685 | Non-Significant | FALSE |
| Ctbp1 | 0.283172 | 9.177792 | 4.085175 | 0.001772 | 0.043538 | -1.0756 | Significant | 0.286975 | 9.177792 | 4.133571 | 0.001632 | 0.025226 | -1.17225 | Significant | FALSE |
| Ctcf | -0.10008 | 7.00763 | -0.96941 | 0.352994 | 0.574312 | -5.75758 | Non-Significant | -0.18283 | 7.00763 | -1.807 | 0.097923 | 0.24125 | -5.01514 | Non-Significant | FALSE |
| Ctdsp2 | 0.024188 | 7.242175 | 0.242649 | 0.812704 | 0.900153 | -6.22954 | Non-Significant | -0.14355 | 7.242175 | -1.45386 | 0.173675 | 0.344009 | -5.52755 | Non-Significant | FALSE |
| Ctnnbip1 | 0.167745 | 6.636605 | 1.313547 | 0.215508 | 0.430324 | -5.33435 | Non-Significant | -0.01849 | 6.636605 | -0.14706 | 0.885719 | 0.943694 | -6.46019 | Non-Significant | FALSE |
| Ctps | 0.034959 | 6.97286 | 0.343047 | 0.737974 | 0.861881 | -6.15365 | Non-Significant | 0.019513 | 6.97286 | 0.195529 | 0.848513 | 0.917678 | -6.50409 | Non-Significant | FALSE |
| Ctsb | 0.255622 | 10.35714 | 2.589706 | 0.024995 | 0.139793 | -3.614 | Non-Significant | 0.247044 | 10.35714 | 2.483028 | 0.030239 | 0.119972 | -4.03457 | Non-Significant | FALSE |
| Ctsf | 0.290717 | 6.938081 | 3.088395 | 0.010217 | 0.097482 | -2.65406 | Significant | 0.263853 | 6.938081 | 2.853675 | 0.015569 | 0.080957 | -3.27358 | Non-Significant | FALSE |
| Ctsl | -0.03617 | 7.630714 | -0.26428 | 0.796408 | 0.890847 | -6.26182 | Non-Significant | -0.14978 | 7.630714 | -1.09943 | 0.294858 | 0.474537 | -5.9802 | Non-Significant | FALSE |
| Cttn | -0.05 | 8.037064 | -0.46872 | 0.648341 | 0.79975 | -6.20061 | Non-Significant | 0.002234 | 8.037064 | 0.021215 | 0.983451 | 0.991087 | -6.61837 | Non-Significant | FALSE |
| Ctxn1 | 0.240862 | 7.068732 | 1.983309 | 0.072628 | 0.239531 | -4.48277 | Non-Significant | -0.21461 | 7.068732 | -1.77598 | 0.103129 | 0.248533 | -5.05511 | Non-Significant | FALSE |
| Cuedc2 | 0.074839 | 7.702492 | 0.399671 | 0.696984 | 0.833099 | -6.22142 | Non-Significant | -0.27108 | 7.702492 | -1.44597 | 0.175832 | 0.347042 | -5.57114 | Non-Significant | FALSE |
| Cul1 | -0.02449 | 8.653388 | -0.28806 | 0.778609 | 0.881275 | -6.31615 | Non-Significant | -0.03675 | 8.653388 | -0.43503 | 0.671883 | 0.798432 | -6.55667 | Non-Significant | FALSE |
| Cul2 | -0.19302 | 6.861137 | -2.17663 | 0.051963 | 0.197385 | -4.16262 | Non-Significant | -0.28315 | 6.861137 | -3.27036 | 0.007379 | 0.053261 | -2.56428 | Non-Significant | FALSE |
| Cul3 | -0.25986 | 7.607884 | -3.04265 | 0.01109 | 0.102057 | -2.79594 | Non-Significant | -0.46027 | 7.607884 | -5.40322 | 0.000209 | 0.011476 | 0.920136 | Significant | FALSE |
| Cul4a | -0.22985 | 8.084624 | -2.83239 | 0.016175 | 0.118031 | -3.16971 | Non-Significant | -0.26621 | 8.084624 | -3.31718 | 0.006788 | 0.050947 | -2.56125 | Non-Significant | FALSE |
| Cul9 | -0.10114 | 7.813732 | -0.99865 | 0.339243 | 0.562899 | -5.80322 | Non-Significant | -0.05905 | 7.813732 | -0.58934 | 0.567438 | 0.723988 | -6.42409 | Non-Significant | FALSE |
| Cux1 | -0.0895 | 7.624973 | -0.74729 | 0.470428 | 0.673419 | -6.01304 | Non-Significant | -0.26331 | 7.624973 | -2.2047 | 0.049472 | 0.158665 | -4.44444 | Non-Significant | FALSE |
| Cux2 | -0.04145 | 7.807309 | -0.3496 | 0.733187 | 0.857871 | -6.24289 | Non-Significant | -0.12455 | 7.807309 | -1.05824 | 0.312439 | 0.494218 | -6.03408 | Non-Significant | FALSE |
| Cx3cl1 | 0.255917 | 9.760126 | 2.540136 | 0.02731 | 0.146861 | -3.70188 | Non-Significant | 0.651443 | 9.760126 | 6.391904 | 4.94E-05 | 0.00588 | 2.330517 | Significant | TRUE |
| Cxcl14 | 0.467615 | 7.238514 | 3.377797 | 0.006094 | 0.077341 | -2.18705 | Significant | 0.356015 | 7.238514 | 2.586351 | 0.025145 | 0.107056 | -3.75138 | Non-Significant | FALSE |
| Cxxc5 | 0.27584 | 7.048545 | 2.555944 | 0.02655 | 0.143927 | -3.55056 | Non-Significant | 0.33348 | 7.048545 | 3.138036 | 0.009347 | 0.060464 | -2.78795 | Non-Significant | FALSE |
| Cyb561 | 0.156525 | 7.554841 | 1.940129 | 0.078193 | 0.247216 | -4.5892 | Non-Significant | 0.076836 | 7.554841 | 0.957521 | 0.358701 | 0.542405 | -6.11127 | Non-Significant | FALSE |
| Cyb5r4 | -0.41355 | 7.350245 | -3.73329 | 0.003256 | 0.057361 | -1.614 | Significant | -0.30366 | 7.350245 | -2.78494 | 0.017614 | 0.086928 | -3.45686 | Non-Significant | FALSE |
| Cyfip1 | -0.08305 | 7.594579 | -0.73003 | 0.480499 | 0.681195 | -6.01819 | Non-Significant | -0.00263 | 7.594579 | -0.02338 | 0.981761 | 0.991042 | -6.59263 | Non-Significant | FALSE |
| Cyfip2 | 0.098213 | 10.4884 | 0.737662 | 0.476029 | 0.677008 | -6.08638 | Non-Significant | 0.176485 | 10.4884 | 1.312576 | 0.215824 | 0.391053 | -5.79156 | Non-Significant | FALSE |
| Cyhr1 | -0.05156 | 9.106892 | -0.59923 | 0.561055 | 0.743079 | -6.18605 | Non-Significant | 0.06059 | 9.106892 | 0.704133 | 0.49586 | 0.661971 | -6.40436 | Non-Significant | FALSE |
| Cyld | -0.18666 | 9.031132 | -2.43353 | 0.033025 | 0.163432 | -3.87797 | Non-Significant | -0.2394 | 9.031132 | -3.12663 | 0.00954 | 0.060607 | -2.92917 | Non-Significant | FALSE |
| Cyp46a1 | 0.43233 | 7.997097 | 5.601758 | 0.000155 | 0.023095 | 1.278666 | Significant | 0.446771 | 7.997097 | 5.856514 | 0.000106 | 0.007744 | 1.586935 | Significant | FALSE |
| Cyp51 | -0.13791 | 7.348339 | -1.45151 | 0.174314 | 0.382008 | -5.25982 | Non-Significant | -0.07496 | 7.348339 | -0.79827 | 0.441464 | 0.618327 | -6.24155 | Non-Significant | FALSE |
| Cyth2 | 0.141354 | 7.561008 | 1.060238 | 0.311567 | 0.533489 | -5.72062 | Non-Significant | 0.1928 | 7.561008 | 1.459826 | 0.17206 | 0.341631 | -5.54201 | Non-Significant | FALSE |
| Cyth3 | 0.078813 | 7.034786 | 0.845006 | 0.415955 | 0.630405 | -5.86048 | Non-Significant | 0.050782 | 7.034786 | 0.554285 | 0.590379 | 0.741226 | -6.36764 | Non-Significant | FALSE |
| D17Wsu92[illegible] | -0.02742 | 9.757705 | -0.34757 | 0.734663 | 0.858997 | -6.31412 | Non-Significant | 0.018912 | 9.757705 | 0.238379 | 0.815932 | 0.895501 | -6.63333 | Non-Significant | FALSE |
| D1Ertd622[illegible] | -0.24945 | 7.398983 | -2.44826 | 0.03217 | 0.161032 | -3.78678 | Non-Significant | -0.52932 | 7.398983 | -5.22681 | 0.000274 | 0.012459 | 0.653567 | Significant | FALSE |
| D430019H[illegible] | 0.134527 | 9.010516 | 1.506428 | 0.159882 | 0.364501 | -5.28197 | Non-Significant | 0.014773 | 9.010516 | 0.16559 | 0.871458 | 0.932759 | -6.64981 | Non-Significant | FALSE |
| D430041D[illegible] | -0.18625 | 9.705602 | -1.383 | 0.193859 | 0.401617 | -5.44657 | Non-Significant | -0.01611 | 9.705602 | -0.11881 | 0.907547 | 0.953935 | -6.65656 | Non-Significant | FALSE |
| D630045J[illegible] | -0.28447 | 7.499954 | -2.70594 | 0.020296 | 0.128495 | -3.35429 | Non-Significant | -0.03988 | 7.499954 | -0.38545 | 0.707187 | 0.824533 | -6.5116 | Non-Significant | FALSE |
| Dact2 | 0.221896 | 7.04084 | 1.94498 | 0.077549 | 0.246581 | -4.53557 | Non-Significant | -0.10923 | 7.04084 | -0.96598 | 0.354636 | 0.539304 | -6.05257 | Non-Significant | FALSE |
| Dact3 | 0.103177 | 8.989188 | 0.724386 | 0.483821 | 0.684023 | -6.10128 | Non-Significant | -0.13448 | 8.989188 | -0.94547 | 0.364552 | 0.549702 | -6.2045 | Non-Significant | FALSE |
| Dag1 | 0.057708 | 7.36516 | 0.575077 | 0.576713 | 0.753017 | -6.09352 | Non-Significant | 0.102086 | 7.36516 | 1.027219 | 0.326188 | 0.507867 | -6.02593 | Non-Significant | FALSE |
